# The Phylogeny and the Evolution of Parasitic Strategies in Trematoda

**DOI:** 10.1101/2024.08.09.607286

**Authors:** Chuan-Yu Xiang, Ivan Jakovlić, Tong Ye, Rui Song, Hong Zou, Gui-Tang Wang, Wen-Xiang Li, Dong Zhang

## Abstract

Trematodes are obligatory parasites that generally must transmit between hosts to complete their life cycle. They parasitize varying numbers of intermediate hosts (0, 1 or 2), but the evolutionary history of these strategies and the ancestral states remain unknown. We conducted the ancestral state reconstruction of the number of intermediate hosts using mitogenomic (Trematoda) and nuclear-genomic (Neodermata) topologies. Aspidogastrea was identified as the sister-group (“basal”) to all other Trematoda using a range of approaches, so it is crucial for studying the evolutionary history of trematodes. However, there is only one transcriptome available for this lineage, and mitochondrial genomes (mitogenomes) remain unavailable. Herein, we sequenced mitogenomes of two aspidogastreans: *Aspidogaster ijimai* and *Aspidogaster conchicola*. As the ancestral state reconstruction analysis is topology-sensitive, we tested multiple phylogenetic strategies, comprising the outgroup selection, phylogenetic models, partitioning strategies, and topological constraints. These mitogenomic phylogenies exhibited pronounced topological instability, with Aspidogastrea resolved as the “basal” radiation in most, but not all, topologies. Based on our analyses, Cestoda was the optimal outgroup choice, and the “heterogeneous” CAT-GTR model in PhyloBayes was the optimal model choice. We inferred the time tree and conducted ancestral state reconstruction analyses using this “optimal” topology, as well as constrained mitogenomic and nuclear genomic topologies. Results were ambiguous for some lineages, but scenario that received the strongest support is the direct life cycle (no intermediate hosts) in the ancestors of Trematoda (proto-trematodes) and Aspidogastrea (proto-aspidogastreans), while the ancestor of Digenea (proto-digeneans) had two intermediate hosts. The inferred scenario indicates that host strategies are relatively plastic among trematodes, putatively comprising several independent host gains, and multiple host losses. We propose a timeline for these events and discuss the role that alternating sexual and asexual generations putatively played in the evolution of complex parasitic life histories in digeneans.

## 1. Introduction

Neodermata, a subphylum of Platyhelminthes, is the most speciose obligatory parasitic clade of animals on the planet (Carlson, et al. 2020). Traditionally, it was divided into three classes: Monogenea, Trematoda (flukes), and Cestoda (tapeworms), but Monogenea is almost certainly paraphyletic and split into two classes: Monopisthocotylea and Polyopisthocotylea (Justine 1998; Brabec, et al. 2023). Despite the paraphyly, members of both “monogenean” classes have similar life history characteristics: primarily ectoparasites (parasitizing external surfaces) of fish, amphibians, and reptiles, with no intermediate hosts (Reed, et al. 2009). Monogeneans can reproduce both asexually or sexually, but they do not alternate between the two strategies (Zhang 1999). As common parasites of all domestic animals and humans, the remaining two classes have major medical and veterinary importance. Cestodes are endoparasites, i.e parasitic within the bodies of hosts (e.g. the abdominal cavity or intestinal tract), exhibit a life cycle with alternating sexual and asexual generations, and typically have one or two intermediate hosts (Smyth and McManus 1989).

Trematoda are also endoparasitic and divided into two subclasses: the smaller Aspidogastrea, and Digenea, which comprises most of the known trematodes. Aspidogastreans have either none or one intermediate host (mollusks), whereas digeneans have one (mollusks; e.g. *Schistosoma* and *Fischoederius*) or two (mollusks, Arthropoda, or lower vertebrates; e.g. *Paragonimus*) intermediate hosts (Cribb, et al. 2001). However, the mollusk hosts of the two taxa are different: aspidogastreans mainly parasitize bivalves, while digeneans mainly parasitise gastropods (Zhang 1999). Regarding the definitive hosts, aspidogastreans without an intermediate host mainly parasitise crustaceans, mollusks, or lower vertebrates, whereas aspidogastreans with one intermediate host mainly parasitize lower vertebrates (Alves, et al. 2015). For digeneans, the definitive hosts are primarily higher vertebrates such as birds and mammals (rarely reptiles) (for details, see Dataset S2: sheet “Host”).

Regarding life history, all digeneans have asexual generations (Alves, et al. 2015). For species with one intermediate host (such as *Schistosoma*), after the intermediate host swallows a miracidium (hatched from an egg in water), it grows and develops inside the host. After a period of time, the miracidium starts to produce a large number of cercaria (up to several hundred), which are commonly released via the host’s digestive tract into the environment. Finally, cercariae must infect the definitive host in order to reproduce sexually (Wright 1967; Jourdane and Mingyi 1987). For species with two intermediate hosts (such as *Paragonimus westermani*): after developing into cercariae, they enter the second intermediate host via predation and develop into metacercariae. Finally, metacercariae must also infect the definitive host (via predation of the 2^nd^ intermediate host) in order to reproduce sexually (Kim 1984; Bengtsson 2003). These interchanging asexual and sexual reproduction strategies maximize the quantity of offspring (Helden and Dixon 2002).

Aspidogastreans do not have an asexual generation, and they potentially hold a key place in the evolution of Trematoda. This small subclass contains species that mainly parasitize the upper gastrointestinal tract of fishes, mollusks and turtles (Alves, et al. 2015). Morphologically, they are characterized by the ventral attachment alveoli and possess many “primitive” life-history traits, so this lineage is traditionally regarded as the sister lineage to all other Trematoda (Olson, et al. 2003). Specifically, it has been proposed that aspidogastreans are poorly adapted to parasitism, with examples including low host specificity and a complex nervous system (Rohde 1971), the ability of some species to lay eggs and complete the entire ontogenetic development in molluscan hosts without a undergoing larval stage (Tang Chongti 2015), and the ability of some species to survive for a long time outside the host (Ferguson, et al. 1999). Due to their putative key position in the phylogeny of Trematoda, Aspidogastrea is also crucial for understanding the evolution of parasitic strategies in this class.

Currently, the number of hosts (both intermediate and definitive) is known for the majority of extant Trematoda species. However, the numbers of intermediate hosts in the ancestors of Trematoda, Aspidogastrea and Digenea (proto-trematodes, proto-aspidogastreans and proto-digeneans) remain unknown. In the light of mixed strategies of contemporary Aspidogastrea, the ancestral state for this lineage could be both 0 and 1 intermediate hosts. Following this reasoning, the ancestral state for Digenea could be both 1 and 2. Finally, due to the unresolved relationships among the major lineages of Neodermata (Brabec, et al. 2023), it is difficult to infer the ancestral state for Trematoda, as it could be 0 or 1. This allows multiple scenarios among the three lineages. Importantly, some scenarios require only gains of hosts (an increase in the number of hosts); for example, 0 as the ancestral state for Trematoda and Aspidogastrea, and 1 for Digenea, followed by independent gains of an additional intermediate host in certain aspidogastrean and digenean lineages. Other types of scenarios require both gains and losses of hosts; for example, 1 as the ancestral state for Aspidogastrea, and 2 for Digenea would both require independent gains of hosts in the ancestral lineages, followed by losses of hosts in some contemporary lineages of these two subclasses. To shed light on the evidently complex evolution of this unique ecological strategy, we set out herein to infer the ancestral host number states for Trematoda, Digenea, and Aspidogastrea.

Molecular dating (also known as molecular clocks or time trees), is commonly used to unravel the evolutionary history between different species and estimate the timing of lineage divergence (Lanfear, et al. 2010). Besides, the ancestral state reconstruction is also an important way to explore the evolutionary history of species (Meade and Pagel 2022). Both types of analyses rely on reliable phylogenetic tree topology, but some questions remain open in the phylogeny of Trematoda, largely due to the scarcity of markers with sufficient resolution. Morphological data in small parasitic animals often carry a limited amount of signal, exhibit convergent evolution, and host-induced morphological variation. For example, the internal relationships of the subclass Digenea remained unresolved even after 203 morphological characters were applied simultaneously to resolve them (Brooks, et al. 1985). Therefore, molecular data are necessary to resolve the phylogenetic relationships of Trematoda and the position of Aspidogastrea. The commonly-used single-locus molecular markers largely support the Aspidogastrea as the sister lineage to the remaining Trematoda (henceforth referred to as “basal Aspidogastrea” hypothesis), but they also often produce variable phylogenetic relationships across studies and datasets (Giese, et al. 2015; Atopkin, et al. 2018) (for details, see Text S1: “Phylogenetic analysis based on commonly-used single-locus molecular markers”).

Compared to the morphological data and single-locus molecular markers, genomic data offer much higher resolving power (Huyse, et al. 2008; Yuan, et al. 2020). However, likely due to their lesser veterinary/medical importance, aspidogastreans remained non-represented in terms of genomic data until very recently, when Brabec et al. sequenced the transcriptome of *Aspidogaster limacoides*, and found support for the “basal Aspidogastrea” hypothesis (Brabec, et al. 2023). While the nuclear genome contains more phylogenetic signal than the mitochondrial genome (mitogenome), the lineage coverage of Trematoda remains much better for mitogenomes (51 mitogenomes vs. 23 nuclear genomes, most of which belong to *Schistosoma*). Several studies that relied on mitogenomes for the phylogenetic reconstruction in Trematoda provided considerable novel insights into their phylogeny (Huyse, et al. 2008; Suleman, et al. 2019; Suleman, et al. 2021). However, mitogenomes are also not flawless molecular markers, as they are susceptible to compositional heterogeneity, substitutional saturation, and several other potential sources of error in phylogenetic reconstruction (Rubinoff and Holland 2005; Zhang, et al. 2019). Indeed, previous mitogenomic studies did not fully resolve the topological instability in Trematoda. For example, a mitochondrial phylogenomic study found that the Bayesian Inference (BI) method produced a paraphyletic order Diplostomida and monophyletic order Plagiorchiida, while the Maximum Likelihood (ML) method produced paraphyletic Plagiorchiida and monophyletic Diplostomida (Li, et al. 2020). Furthermore, Kenny, et al. (2019) found that topologies differed depending on whether the standard “homogeneous” (assuming compositional homogeneity) evolutionary models, such as ML or standard BI as employed by MrBayes, or models accounting for compositional heterogeneity, such as CAT-GTR in PhyloBayes, were used. As a result of this topological instability and/or poor lineage coverage, there are multiple open questions in the phylogeny of Trematoda. Furthermore, despite the critical role that Aspidogastrea probably played in the evolutionary history of Trematoda, there are currently no mitogenomes available for this lineage.

In order to investigate the phylogeny and evolution of Trematoda, for this study we sequenced mitogenomes of two Aspidogastrea species: *Aspidogaster ijimai* and *Aspidogaster conchicola*. We also assembled the mitogenome of *Aspidogaster limacoides* from the raw genome sequencing data generated by Brabec, et al. (2023). Along with other Trematoda mitogenomes available in public databases, we applied these data to accomplish two main objectives. First, we reconstructed the phylogeny of Trematoda. In the light of the topological instability observed in previous studies, we thoroughly tested the impacts of different datasets and methodological approaches: 1. outgroup selection (monogeneans or cestodes); 2. homogeneous and site-heterogeneous mixture phylogenetic models; 3. partitioning strategies; and 4. removal of partitions that violate the assumptions of stationarity and homogeneity. Our analyses indicated that Cestoda was the optimal outgroup choice and that the site-heterogeneous mixture phylogenetic model in PhyloBayes (CAT-GTR) was the optimal phylogenetic algorithm. We tested the performance of these optimal parameters in combination with both nucleotide and amino acid alignments (NUCPBc and AAPBc, “optimal combination” topologies), and used the NUCPBc to infer the time tree. In addition, we also integrated the results of previous studies to reconstruct a constrained tree fixed at the superfamily relationships level. Following this, we used the two “optimal combination” topologies and the constrained tree to study the evolution of intermediate host numbers in Trematoda. For this, we inferred data about the intermediate host numbers for the 53 Trematoda species included in the mitogenomic dataset, as well as all other Neodermata species included in the study of Brabec, et al. (2023). We also incorporated key geological events. Finally, we performed the ancestral state reconstruction and inferred the likelihood of different host loss and gain scenarios in the evolutionary history of Trematoda.

## 2. Materials and methods

### 2.1. Specimen collection and identification

*Aspidogaster ijimai* was collected from the intestine of a common carp (*Cyprinus carpio*) specimen from Boyang Lake (115°47′-116°45′ E, 28°22′-29°45′ N) in Jiangxi Province (China) and *Aspidogaster conchicola* from the intestine of a black carp (*Mylopharyngodon piceus*) specimen from Wuhu Lake (113°47′24′′ E, 30°11′24′′ N), Hubei Province (China). The two parasitic species were identified based on their morphological characteristics (Wu et al., 1991) and ITS sequences (SRR28778872 and SRR28779009), which exhibited >99.0% similarity with *A. ijimai* (DQ345322.1) and *A. conchicola* (DQ345317.1 and HE863965.1) (Chen, et al. 2010), respectively. DNA extraction, and mitogenome amplification, sequencing, annotation, and comparative analyses, were conducted according to Zhang, et al. (2019), so details are given in Text S1.

### 2.2. Phylogenetic analyses

The methods of phylogenetic tree reconstruction based on Trematoda mitogenome with homogeneity model and heterogeneity model are shown in Text S1 (Text S1: “phylogenetic analyzes”). To test our hypothesis that the underlying reason why *Schistosoma* rather than Aspidogastrea was resolved at the base of multiple topologies was long-branch attraction, we inferred long-branch scores, identified spurious species using the TreeSuite function in PhyloSuite v1.2.3, and performed selection pressure analyses using RELAX and aBSREL (adaptive Branch-Site Random Effects Likelihood) HyPhy tools (Kosakovsky Pond, et al. 2019). Selection analyses were performed on NUCm and NUCc datasets, where *Schistosoma* species and *Trichobilharzia regenti* (Schistosomatidae) were selected as the foreground branches because these lineages had the longest branches and highest long branch scores in all topologies.

As compositional heterogeneity may increase the susceptibility to long-branch attraction phenomena in phylogenetic reconstruction (Uribe, et al. 2019), we assessed the presence of compositional heterogeneity in the NUC and AA datasets using the relative composition variability (RCV) test in TreeSuite (PhyloSuite) and symmetry tests in IQ-TREE (Minh, et al. 2020). Before the test, each NUC dataset was divided into 60 partitions (12 PCGs split by codon position = 12 × 3 = 36 + 24 RNAs), and the AA dataset was divided into 12 partitions. The homogeneous model was used to reconstruct phylogenetic trees with the ultrafast bootstrap with 5000 repetitions (Minh, et al. 2020). The symmetry test was used to detect and remove “bad” partitions against the assumptions of stationarity and homogeneity. The remaining partitions were named “passed” partitions, and used in combination with partitioning strategies to reconstruct phylogenetic trees. Furthermore, we also used “passed” partitions to reconstruct trees with homogeneous and heterogeneous models in the IQ-TREE (AA-passed datasets only) respectively. The results of symmetry tests can be divided into three topology groups based on three different strategies. Herein, we collectively refer to the datasets that passed the symmetry tests (NUCm-passed, AAm-passed, NUCc-passed and AAc-passed datasets) as “passed datasets”. The strategies for reconstructing phylogenetic trees were: 1. using the “passed” datasets, partitioned data, and homogeneous ML model of IQ-TREE (NUCSYMm, AASYMm, NUCSYMc and AASYMc, Figures S1-S4); 2. using the “passed” datasets, nonpartitioned dataset, and homogeneous ML model of IQ-TREE (NUCSYMm-Ho, NUCSYMc-Ho, AASYMm-Ho and AASYMc-Ho; Figures S4-S8); 3. using the “passed” datasets, nonpartitioned dataset, and heterogeneous ML model of IQ-TREE (AASYMm-He and AASYMc-He; Figures S9-S10).

In response to the diverse mitogenome-based topologies obtained and the uncertain phylogenetic position of Aspidogastrea, we collected Trematoda topologies obtained in previous studies and constructed trees constrained at the superfamily level using results from previous studies (details of reference topologies are summarized in the “References about topologies and hosts” of Text S1 and Dataset S2, sheet “constraint tree”). The GTR+F+R6 (for the NUCm dataset), mtInv+C50+F+R7 (for the AAm dataset), GTR+F+R7 (for the NUCc dataset), and mtInv+C50+F+R6 (for the AAc dataset) models were selected by ModelFinder (Kalyaanamoorthy, et al. 2017) and used to reconstruct the constraint trees in IQ-TREE (-g parameter): NUCm-cons, AAm-cons, NUCc-cons, and AAc-cons (Figures S11-S14). Furthermore, based on the concatenated sequences, we calculated the gene concordance factor (gCF) and the site concordance factor (sCF) of NUCPBm, AAPBm, NUCPBc and AAPBc topologies using IQ-TREE (Figures S15-S18). Abbreviations used for topologies are presented in Table 1. The MAST algorithm, implemented in IQ-TREE 2.2.0.7.mix, was used to test the weight values of phylogenetic trees. For this analysis, all topologies were divided into four test groups based on the permutations of outgroups (monogeneans or cestodes) and datasets (NUC or AA): monogeneans-NUC, monogeneans-AA, cestodes-NUC, and cestodes-AA. As a heterogeneous model was only available for AA datasets and did not support the CAT-GTR model implemented in PhyloBayes, we used the NUCIQ model (GTR+F+R6 and GTR+F+R7) for the NUCPBm and NUCPBc topologies; and the AAIQ-He model (mtInv+C50+F+R7 and mtInv+C50+F+R6) for the AAPBm and AAPBc topologies.

**Table 1.**
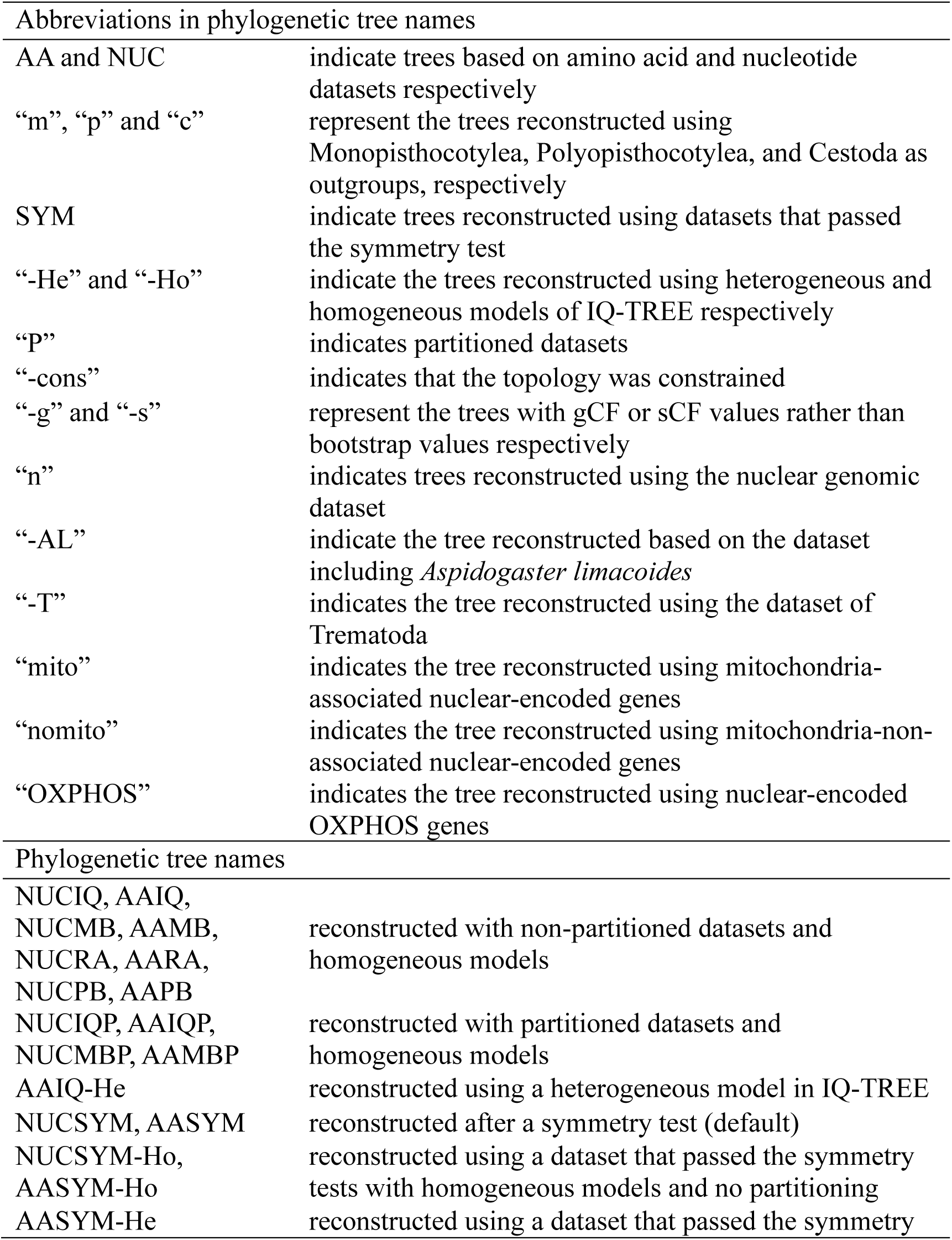
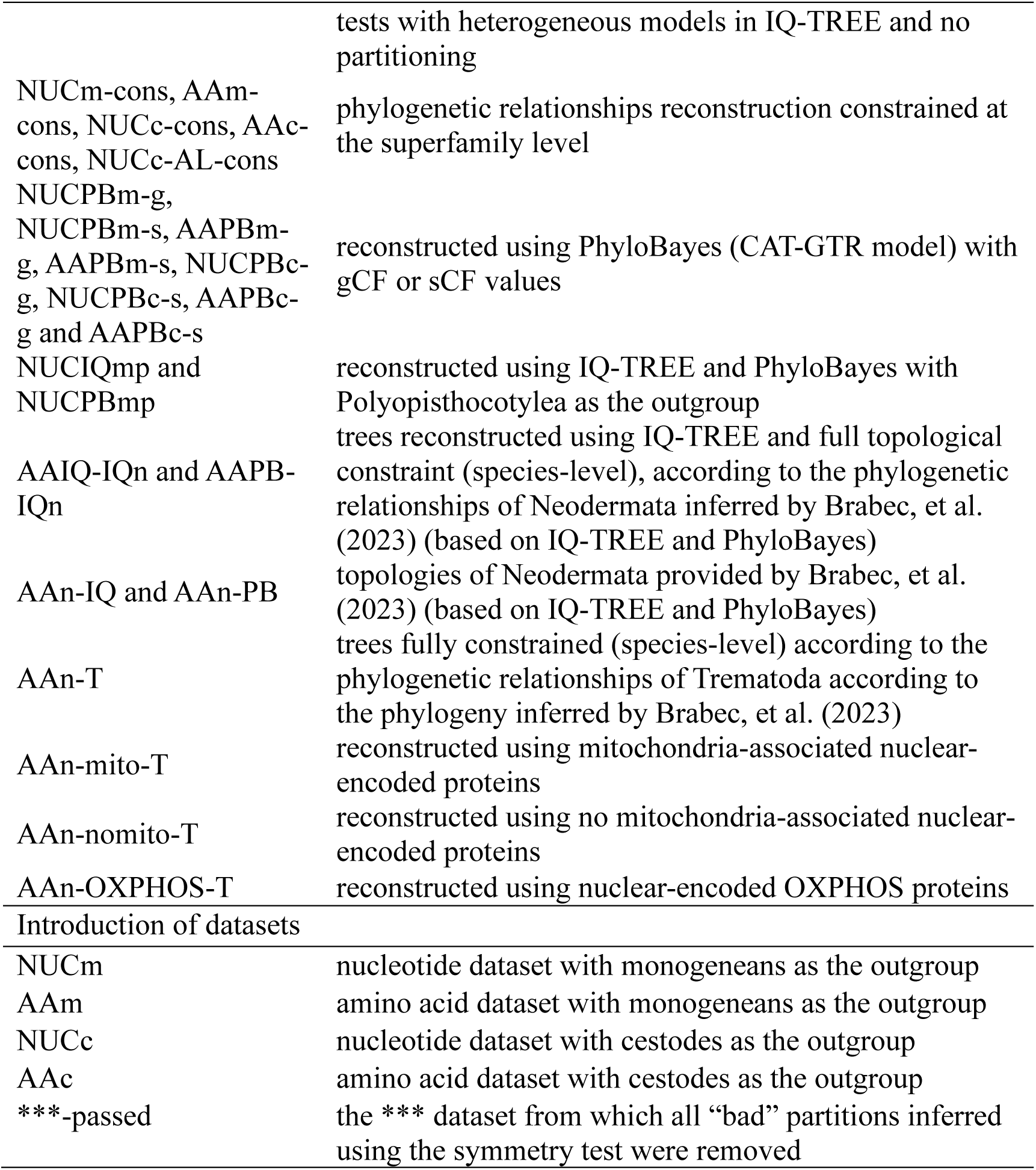
The abbreviations in phylogenetic tree names.

To infer the evolutionary history of intermediate host numbers in the class Trematoda, we performed molecular dating analysis using topologies of mitogenome-based constrained trees with Cestoda as an outgroup, and topology based on nuclear genomic data inferred by Brabec, et al. (2023). The approximate divergence times among the Neodermata classes were estimated using MCMCTree (dos Reis and Yang 2017). We used the GTR model for mitochondrial nucleotides (selected by ModelFinder). Two nodes were chosen as calibration points: the estimate of the origin of *Schistosoma* was between the Early Cretaceous and Paleocene (66-145 Ma) (Peterson, et al. 2004; Parfrey, et al. 2011), whereas the origin of the genus *Paragonimus* is presumed to have taken place between the Eocene and Pliocene (7-44 Ma) (Vainutis, et al. 2022). For molecular dating analysis using genomic protein sequences, we constrained the phylogenetic relationships in Trematoda based on the topology provided by Brabec, et al. (2023) and used the LG+F+C60 model (selected by ModelFinder and IQ-TREE). We used two calibration points: the origin of *Schistosoma* was the same as mentioned above (66-145 Ma), and the origin of Fasciolidae was set between 45 and 55 Ma according to (Lotfy, et al. 2008). Runs were conducted with two MCMC chains, and comprised 2.5 million generations sampled every 100 generations (the first 5000 trees of each run were discarded as burnin). We assessed the convergence of MCMCTree via two indicators: effective sample size (ESS) value above 200, and the posterior mean time values of the two MCMC chains forming a diagonal (dos Reis and Yang 2017).

Finally, to explore whether the topology is correlated with the gene group function, we used a subset of the nuclear genomic (and transcriptomic) dataset used by Brabec, et al. (2023), comprising all 19 species of Trematoda and 3 species of Polyopisthocotylea. PhyloFisher (Tice, et al. 2021) was used to categorize and filter homologous proteins into three groups as described in Zhang, et al. (2024) (for details, see Text S1: “Categorize and filter homologous nuclear genome proteins by PhyloFisher”). ModelFinder and IQ-TREE were used to select the optimal model for phylogenetic tree reconstruction: 1. mitochondria-associated-directly group with LG+C30+F+R6 (AAn-mito-T topology), OXPHOS-associated group with LG+C20+F+R6 (AAn-OXPHOS-T topology), and mitochondria-non-associated group with JTTDCMut+F+R4+C60 (AAn-nomito-T topology).

### 2.3. Ancestral state reconstruction

We collected the number of intermediate hosts and definitive host species of these 53 species of Trematodes by referring to a wide range of scientific literature. For details, see “References about topologies and hosts” in Text S1 and Dataset S2: sheet “Host”. Besides, we also collected the number of intermediate hosts and definitive host species for the remaining Neodermata species included in Brabec, et al. (2023) from a range of sources (please see “References about topologies and hosts” in Text S1, and Dataset S2: sheet “Host_Neodermata”). Taking the number of intermediate hosts as the target feature, the maximum likelihood method (ML) of BayesTraitsV3 (Meade and Pagel 2022) was used to conduct the ancestral state reconstruction based on the constraint tree with cestodes as the outgroup (NUCc-cons and AAc-cons). To increase the number of available Aspidogastrea, we extracted the incomplete mitogenome of *Aspidogaster limacoides* (containing nine protein-coding genes and *rrnS*) from the previously sequenced transcriptome (Brabec, et al. 2023), reconstructed a constrained tree (NUCc-AL-cons; Figure S19) based on the NUCc-cons topology, and performed the ancestral state reconstruction using BayesTraitsV3 (see Dataset S2: sheet “ancestor”). Among the three species, *A. limacoides* and *A.conchicola* both have one intermediate host, while *A.ijimai* does not have an intermediate host.

The results of ancestral state reconstruction are easily affected by the topology of the phylogenetic tree (Hanson-Smith, et al. 2010; Litsios and Salamin 2012). Brabec, et al. (2023) found that relationships within (but not among) the four classes of Neodermata were stable between a homogeneous model and a heterogeneous model-based phylogenetic reconstruction (47 species in total, including 19 Trematoda species). Accordingly, we selected these 47 species of Neodermata (plus one species of Bothrioplanida as the outgroup) to construct phylogenetic trees for the ancestral state reconstruction. Furthermore, PhyloFisher was used to obtain the homologous proteins of two datasets: 1. Neodermata (48 species); 2. Trematoda (19 Trematoda + 3 Polyopisthocotylea species as outgroups). The former dataset was filtered to retain protein sequences identified in more than 40 species (205 protein sequence files in total), and the latter was filtered to retain 220 protein sequences. As Brabec, et al. (2023) found that nuclear genomic data produce two different topologies of Neodermata, we used these two topologies as constraints for phylogenetic analyses: “constraint 1” was Monopisthocotylea+(Cestoda+(Trematoda+Polyopisthocotylea)), and “constraint 2” was (Monopisthocotylea+Cestoda)+(Trematoda+Polyopisthocotylea). The constrained trees (AAIQ-IQn, AAPB-IQn) were reconstructed using IQ-TREE and 205 homologous protein sequence files mentioned above. The ancestral state reconstruction analyses were based on these two topologies. In addition, because Aspidogastrea was represented by only one species in the nuclear genomic dataset (*A. limacoides*), we tested scenarios where this species had both 0 and 1 intermediate hosts. To further test the sensitivity of these results to topology, we also conducted the ancestral state reconstruction analyses using the topology produced using nuclear genomes in a previous study Brabec, et al. (2023). First, we produced a constrained topology of Trematoda using IQ-TREE analysis and 220 homologous protein sequences mentioned above. To better calibrate the time tree within Trematoda, in addition to mitogenomic data, the nuclear genome dataset was used for the molecular dating analysis.

According to the results that we obtained, Cestoda appears to be a more suitable outgroup for Trematoda than “monogenea” (Polyopisthocotylea and Monopisthocotylea) for mitogenome-based phylogenetic reconstruction. When the ancestral state was reconstructed based on the results of PhyloBayes with cestodes as the outgroup (NUCPBc and AAPBc), the results showed that the ancestral state of Trematoda was two intermediate hosts (for details, see Dataset S2: sheet “filtered out”). However, the ancestor of Aspidogastrea, which was resolved as the earliest radiation of Trematoda, had no intermediate hosts. According to this contradictory finding, we speculated that NUCc-cons and AAc-cons topologies may be more suitable for the ancestral state reconstruction analysis than NUCPBc and AAPBc topologies.

## 3. Results

### 3.1. Topological instability in Trematoda

Partitioning, outgroups, and evolutionary models all affected the topology. Regarding the order-level phylogeny: with Monopisthocotylea (“monogenea”) as the outgroup and standard (“homogeneous”) models, the topologies (AAIQm, NUCIQm, AARAm, NUCRAm, AAMBm and NUCMBm; for phylogram name abbreviations see Table 1) consistently resolved the two Aspidogastrea species sequenced for this study nested within the Diplostomida clade, thus rendering this order paraphyletic (Figures S20-S24, Figure 1a). Contrary to this, the results of “heterogeneous” models (CAT-GTR in PhyloBayes and mtInv+C50+F+R7 in IQ-TREE) mostly placed Aspidogastrea at the root of Trematoda, thus producing monophyletic Diplostomida (AAPBm, NUCPBm, and AAIQm-He; Figure S25, 1b and S26). The only exception was AAIQm-He topology, where Diplostomida was rendered paraphyletic by the two Aspidogastrea species nested within the clade.

**Figure 1.**
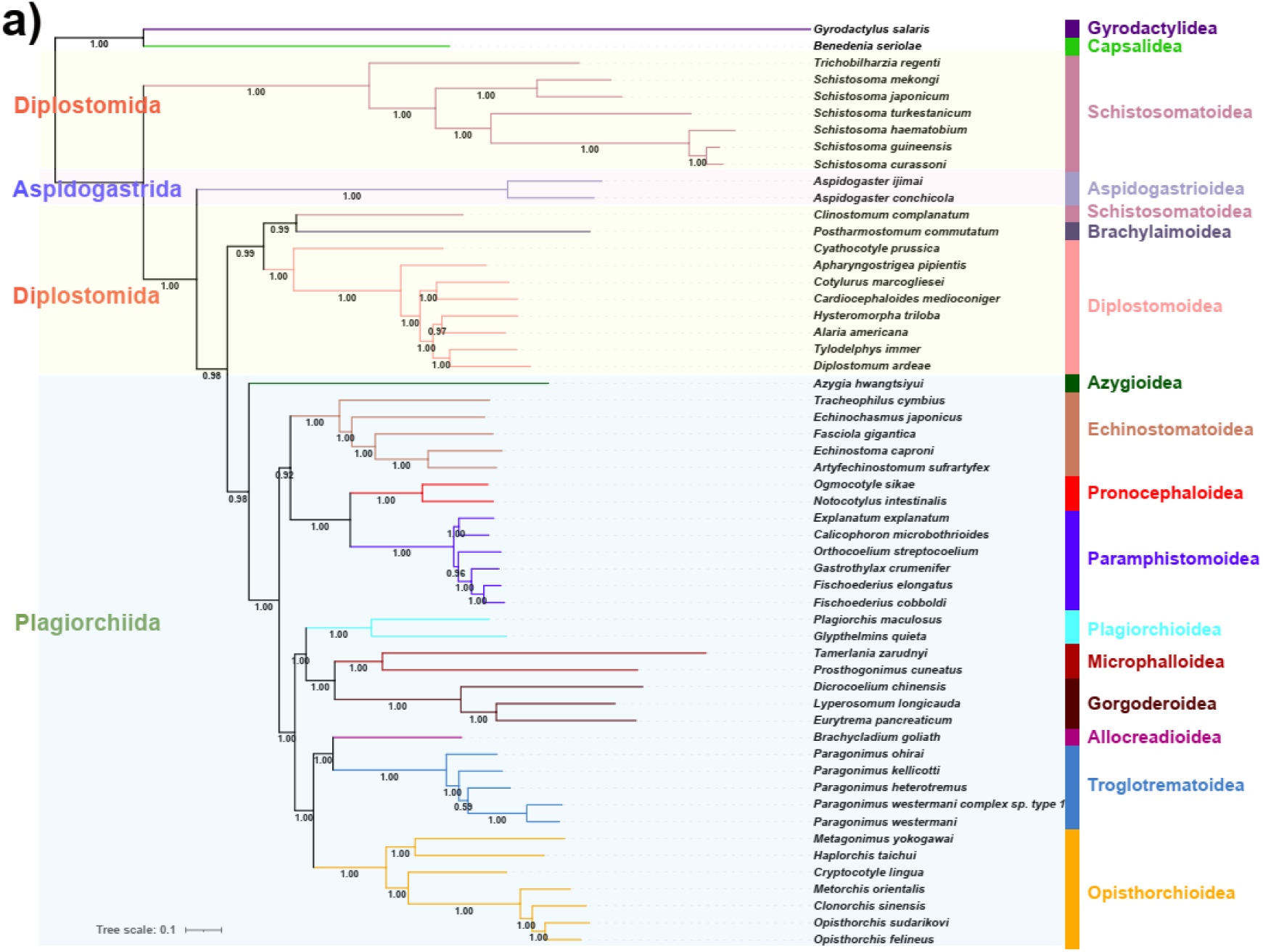
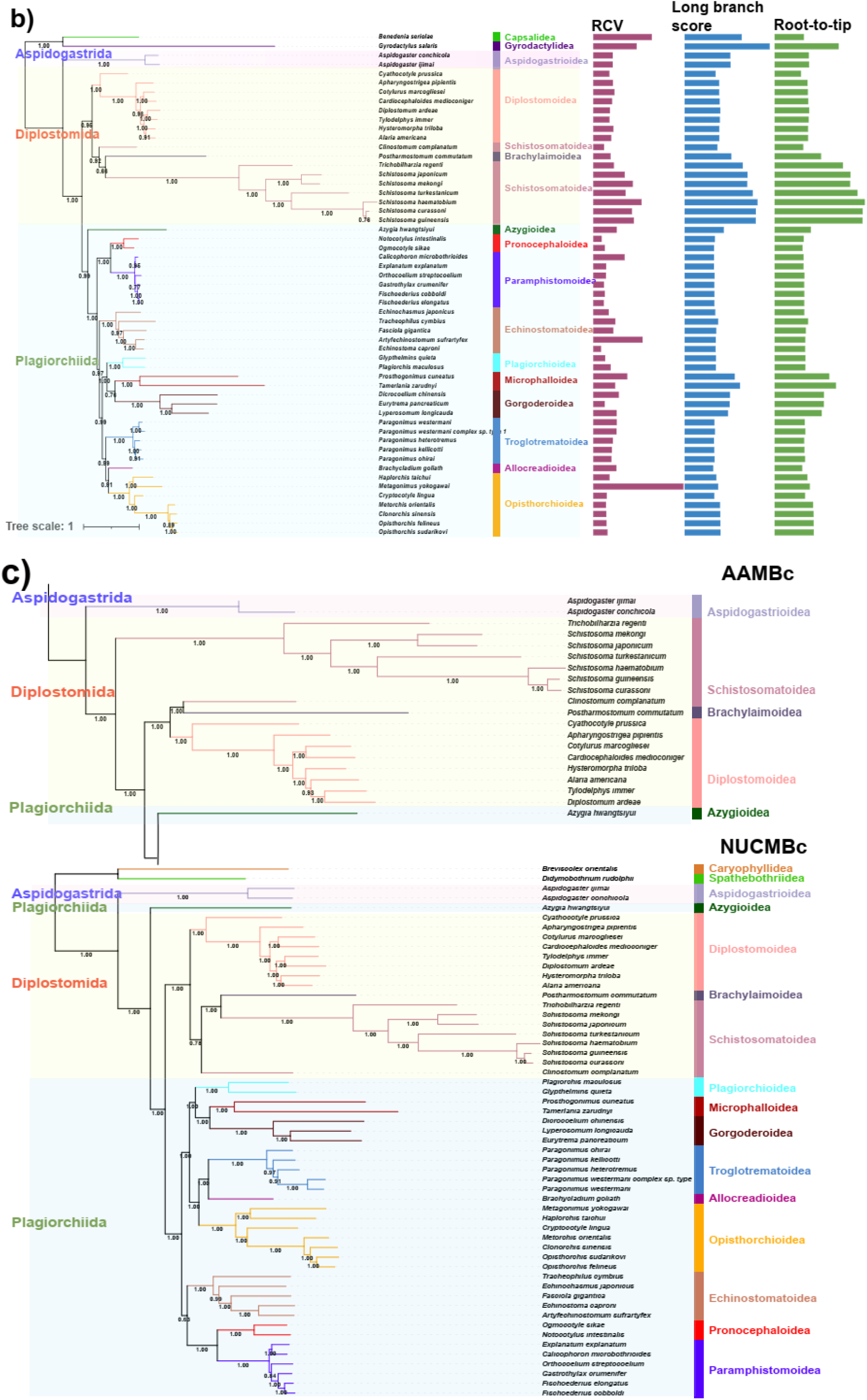
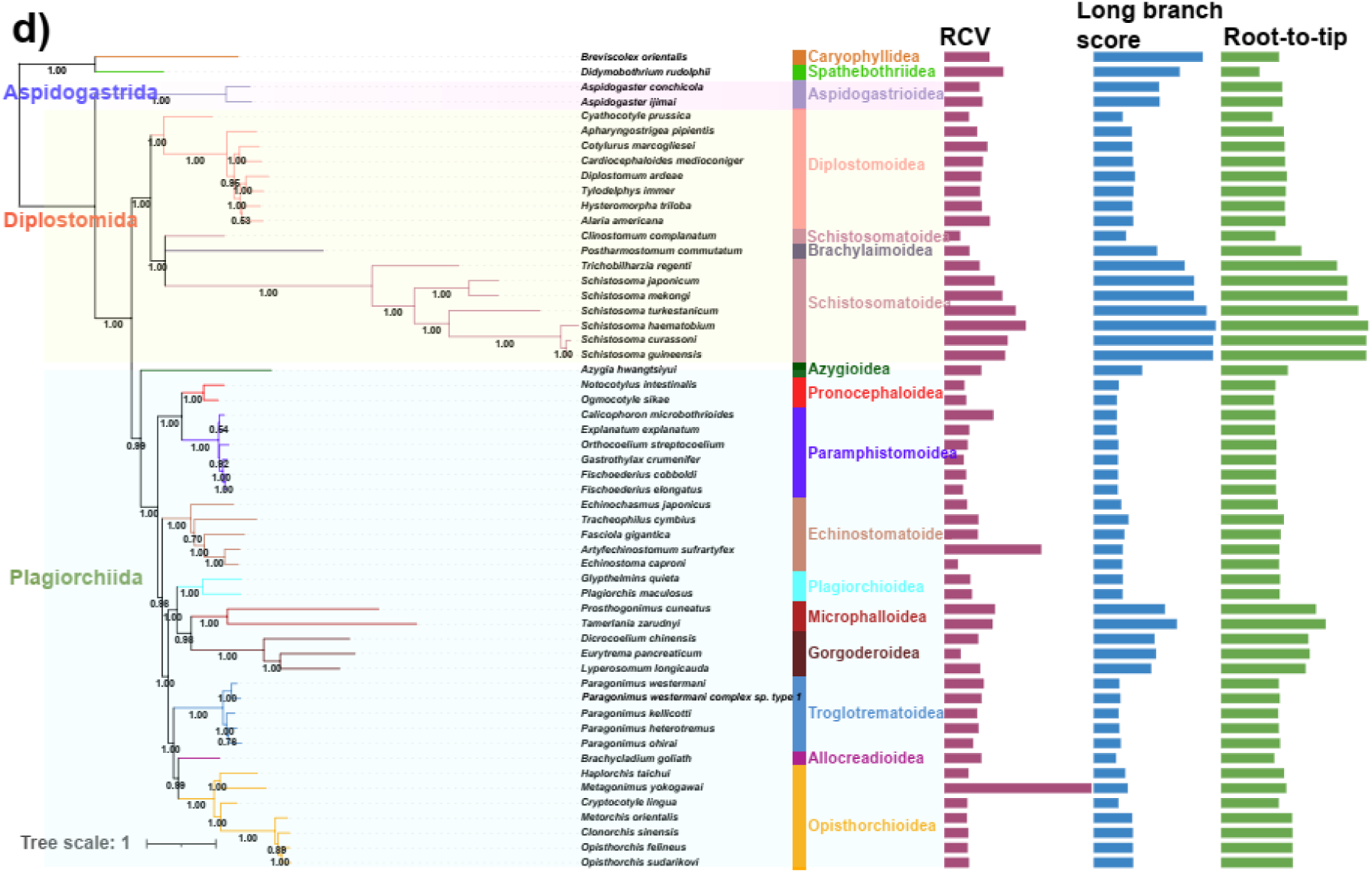
Phylogenetic trees reconstructed using the mitogenomes of Trematoda. With Monopisthocotylea as the outgroup and with (a) the “homogeneous” model of MrBayes, and (b) the “heterogeneous” Bayesian CAT-GTR model implemented in PhyloBayes. With Cestoda as the outgroup and with (c) MrBayes using both nucleotide (AAMBc) and amino acid (NUCMBc) datasets, and (d) the CAT-GTR model of PhyloBayes. If not specified, the nucleotide dataset was used to infer phylogenies. Only a fragment of the AAMBc topology is shown in the panel c; the complete topology is available as Supplementary Figure S31. Bayesian posterior probability values are shown under the branches. RCV is the relative compositional variability.

Interestingly, with Cestoda as the outgroup, the results consistently placed Aspidogastrea at the root of Trematoda, including when homogeneous models were used (AAIQc, NUCIQc, AARAc, NUCRAc, AAMBc, and NUCMBc; Figures S27-S31, Figure 1c). However, topologies differed between the AA dataset and the NUC dataset. For example, the AAMBc analysis produced monophyletic Plagiorchiida and paraphyletic Diplostomida orders, while in the NUCMBc topology, Diplostomida was monophyletic but Plagiorchiida was paraphyletic. Heterogeneous models (AAPBc, NUCPBc and AAIQc-He) partially managed to reduce the paraphyly, with all three orders (Aspidogastrida, Diplostomida and Plagiorchiida) monophyletic in AAPBc and NUCPBc topologies (Figure S32 and 1d), but Diplostomida remaining paraphyletic in the AAIQc-He topology (Figure S10).

As Polyopisthocotylea, rather than Cestoda, was resolved as a sister clade to Trematoda in a recent nuclear phylogenomic study (Brabec, et al. 2023), we also tested the performance of this lineage as the outgroup for mitogenome-based phylogenetic analyses (NUCIQmp and NUCPBmp; Figures S13 and S14; for details, see Text S1: “Phylogenetic analysis with Polyopisthocotylea as the outgroup”). The results produced two different topologies: Diplostomida was paraphyletic in the NUCIQmp, and Trematoda was polytomous in the NUCPBmp topology. Finally, partitioning strategies also affected the topology (for details, see Text S1: “Partitioning”).

### 3.2. Topology testing and evaluation

The long-branch scores, selection analyses, and spurious species identification were used to assess the likelihood of the long-branch attraction phenomena. Among the Trematoda, species from the superfamily Schistosomatoidea exhibited significantly higher long branch scores than other lineages (p<0.01, Figure 1; Dataset S2: sheet “LBS”). Besides, based on the NUCm and NUCc datasets, *Schistosoma* species and *Trichobilharzia regenti* (Schistosomatidae) are evolving under relatively relaxed selection pressures (p<0.05 and k<1; Dataset S2: “hyphy” sheet). Regarding the different outgroups, only a monogenean outgroup species *Gyrodactylus salaris* (Monopisthocotylea) was identified as a spurious species (only in AAPBm and NUCPBm topologies). Interestingly, no spurious species were detected in analyses with cestodes as the outgroup.

Relative composition variability (RCV) and symmetry tests can be used to predict the heterogeneity level of a dataset and its impact on the topology. Species in the superfamily Schistosomatoidea exhibited significantly higher RCV values (p<0.01, Figure 1; Dataset S2: sheet “RCV”). The NUCm-passed (“-passed” in the name of the dataset indicates that it has passed the symmetry test; NUCSYMm and NUCSYMm-Ho, Figures S1 and S6), AAm-passed (AASYMm, AASYMm-Ho and AASYMm-He, Figures S2, S5, and S9) and NUCc-passed datasets (NUCSYMc and NUCSYMc-Ho, Figures S3 and S7) all resolved Aspidogastrea as the sister lineage to Digenea, and all three orders (Aspidogastrida, Diplostomida and Plagiorchiida) were monophyletic. In contrast, the AAc-passed dataset (AASYMc, AASYMc-Ho, and AASYMc-He, Figures S4, S8, and S33) produced paraphyletic Plagiorchiida. For details, see Table 2.

**Table 2.**
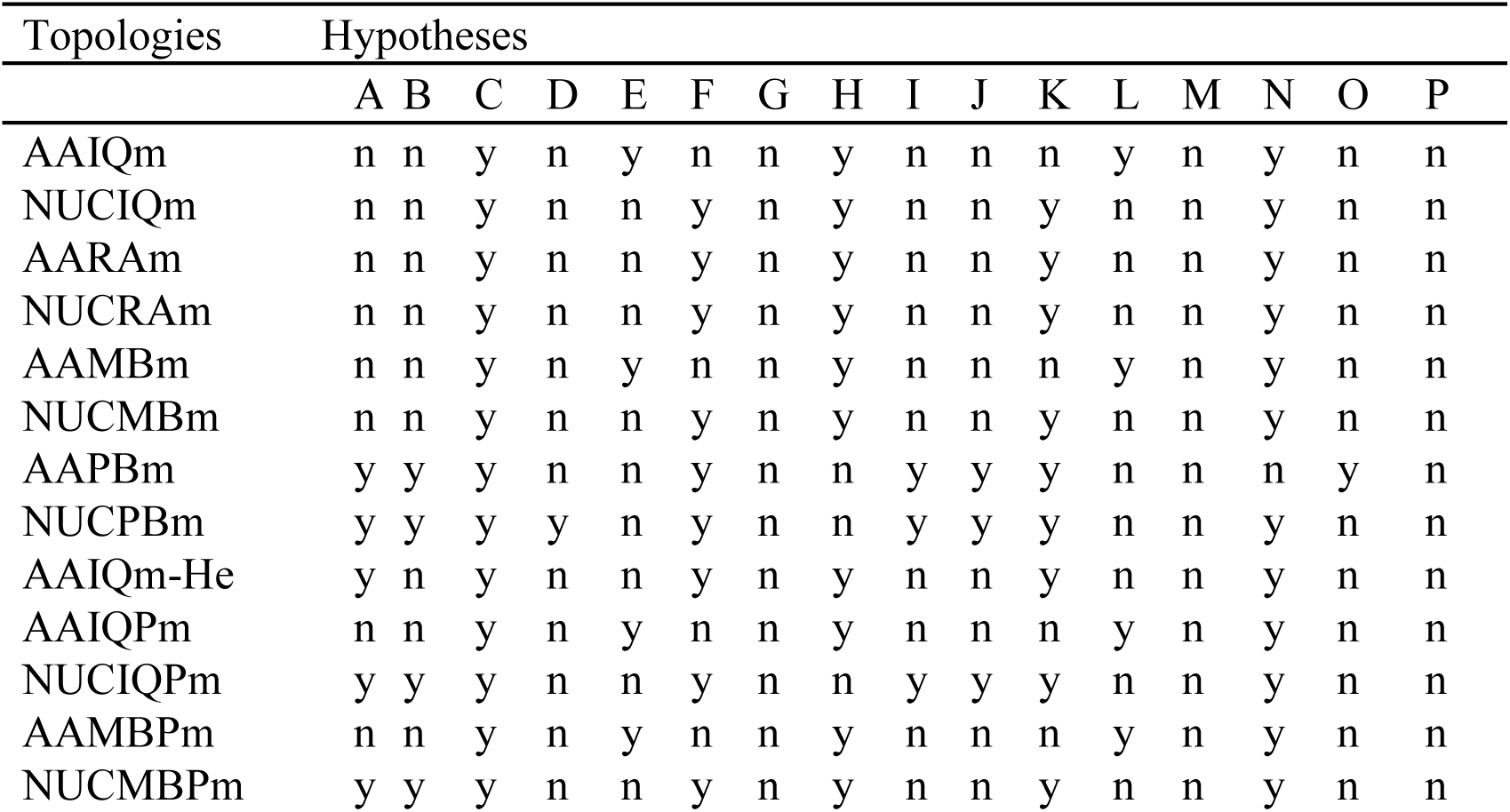
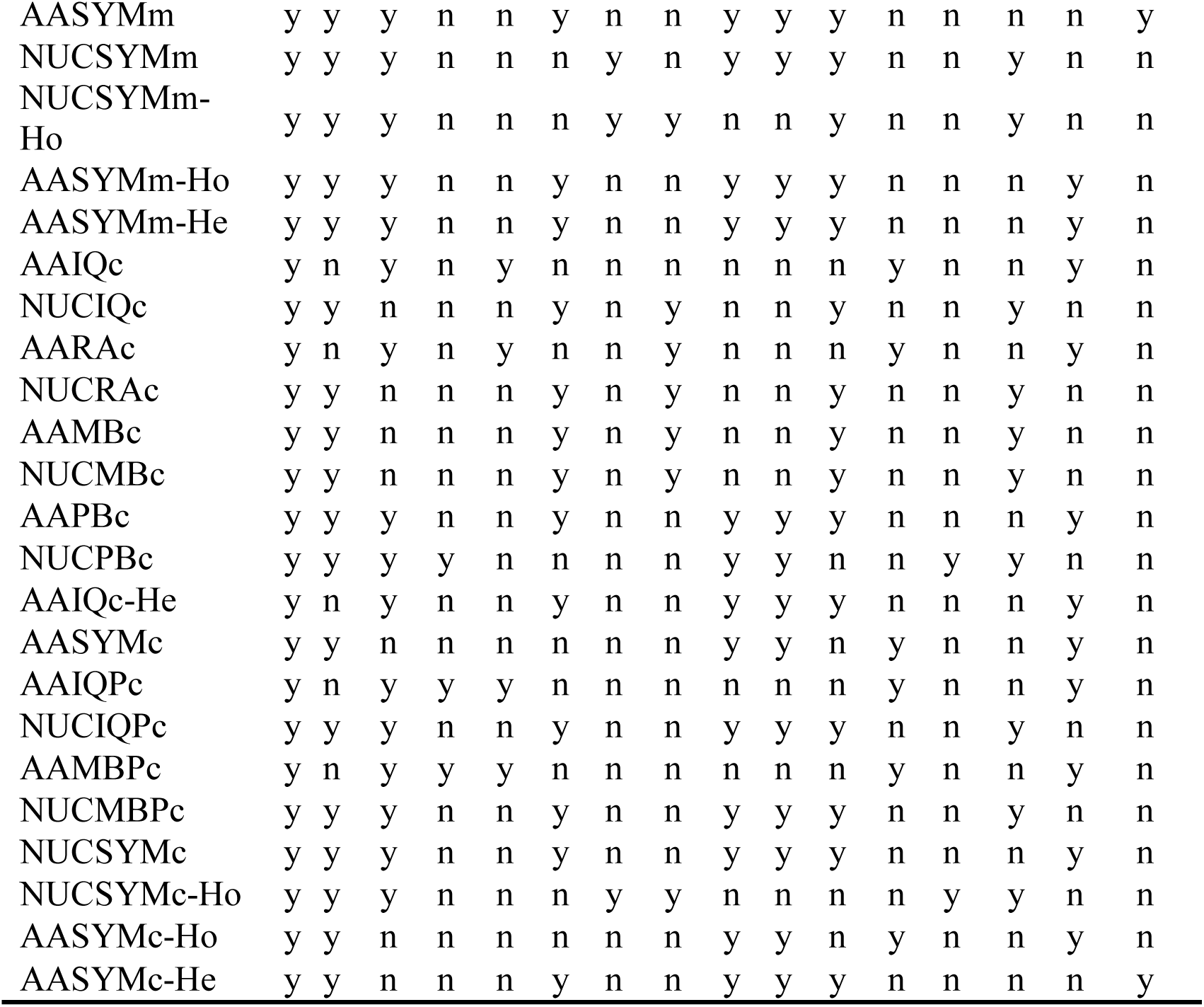
Summary of phylogenetic analyses. Rows show topologies. Columns indicate different phylogenetic hypotheses: (A) monophyletic subclass Digenea (B), monophyletic order Diplostomida, (C) monophyletic order Plagiorchiida, (D) monophyletic superfamily Schistosomatoidea, (E) Gorgoderoidea+Microphalloidea sister clade is the basal radiation in the order Plagiorchiida except for Azygioidea, (F) Gorgoderoidea+Microphalloidea is sister clade to the superfamily Plagiorchioidea, (G) Plagiorchioidea+Microphalloidea is sister-clade to Gorgoderoidea, (H) Echinostomatoidea is sister-clade to Pronocephaloidea+Paramphistomoidea, (I) Echinostomatoidea is the basal Plagiorchiida radiation except for Azygioidea, Pronocephaloidea and Paramphistomoidea, (J) Pronocephaloidea+Paramphistomoidea is the basal Plagiorchiida radiation except for Azygioidea, (K) Plagiorchioidea+Microphalloidea+Gorgoderoidea is the sister clade to Allocreadioidea+Troglotrematoidea+Opisthorchioidea, (L) Plagiorchioidea is sister-clade to Allocreadioidea+Troglotrematoidea+Opisthorchioidea, (M) Allocreadioidea is sister-clade to Troglotrematoidea, (N) Allocreadioidea is sister-clade to Opisthorchioidea, (O) Troglotrematoidea is sister-clade to Opisthorchioidea, (P) Microphalloidea+Gorgoderoidea is sister-clade to Plagiorchioidea. y: yes (hypothesis supported); n: no (hypothesis not supported).

The MAST analysis (Wong, et al. 2022) was used to evaluate the “weight” of the multiple topologies in our analyses. With monogeneans as the outgroup, topologies inferred using the CAT-GTR model of PhyloBayes (NUCPBm and AAPBm) exhibited higher weight values than topologies inferred using homogeneous models. Interestingly, with cestodes as the outgroup, heterogeneous PhyloBayes model NUC topology (NUCPBc) and heterogeneous IQ-TREE model AA topology (AAIQc-He), but not PhyloBayes AA topology (AAPBc), exhibited the highest weight values (detailed MAST analyses results in Dataset S2: sheet “MAST”).

Concordance factor analyses were used to evaluate the different phylogenetic positions of Aspidogastrea obtained in the results of homogeneous and heterogeneous models. The topologies reconstructed using the NUC dataset in combination with PhyloBayes (NUCPBm and NUCPBc) and IQ-TREE (NUCIQm and NUCIQc) were used for concordance factor analyses (the higher the value, the more genes/sites support the branch). The sCF and gCF values (higher value = higher support) supporting the “basal” position of Aspidogastrea were 100 and 73.73 respectively in the NUCPBm topology (Figures S15 and S16); both were 100 in the NUCPBc (Figures S17 and S18); and they were 100 and 46.98 respectively in the NUCIQc topology (Figures S34 and S35). Regarding the sCF and gCF values supporting the Aspidogastrea superfamily nested within the order Diplostomida, they were 39.50 and 27.27 respectively in the NUCIQm topology (Figures S36 and S37). Finally, we confirmed that nuclear data consistently produced the “basal” Aspidogastrea and monophyletic orders (further details in Text S1: “Evolutionary scenarios for Trematoda inferred using the nuclear genome data”).

### 3.3. Molecular dating

Based on the above results, we inferred that Cestoda performed better as an outgroup than “monogenea”, and that PhyloBayes’ CAT-GTR model outperformed other models regardless of the outgroup used. Since the inference of divergence time relies on a reliable phylogenetic tree, and molecular dating is typically performed using nucleotide data instead of amino acid data (Guindon 2020), we selected the topology inferred using PhyloBayes with Cestoda as the outgroup (NUCPBc) for the time tree analysis (NUCPBc-time; Figure 2). To assess the sensitivity to topology, the time tree analysis was also conducted using the nuclear genome-based AAn-T topology (AAn-T-time; Figure 3). The results indicated that Aspidogastrea and Digenea subclasses separated between the Mid-Triassic and Late Jurassic (95% confidence interval: 231-144 Ma) in the mitogenomic topology, and between the early Carboniferous and Late Jurassic (353-149 Ma) in the nuclear topology. Diplostomida and Plagiorchiida diverged between 129-204 Ma in the mitogenomic, and between 119-277 Ma in the nuclear topology.

**Figure 2.**
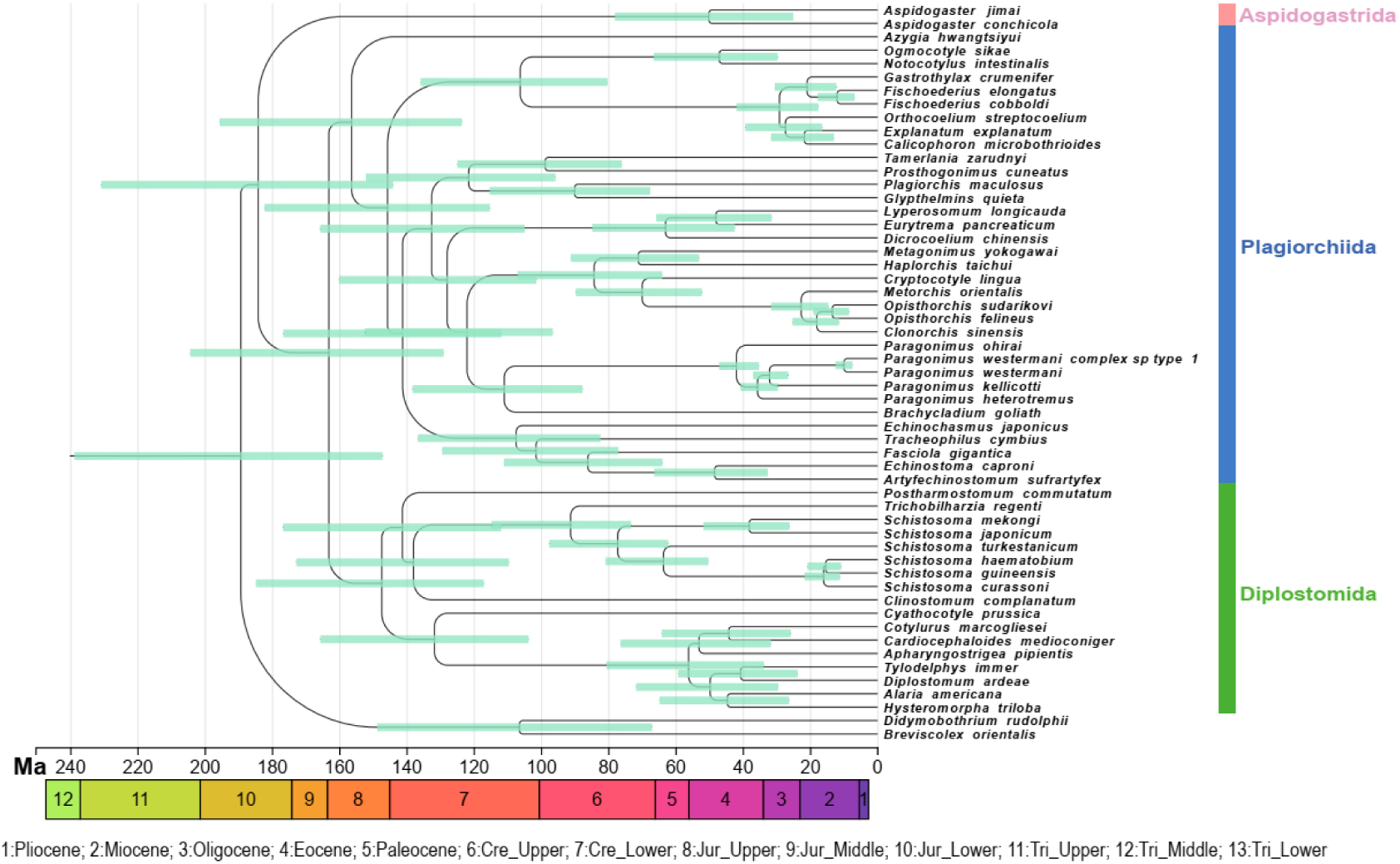
The time tree reconstructed using the NUCPBc topology, inferred using the mitochondrial nucleotide dataset of Trematoda, with Cestoda as the outgroup, and the CAT-GTR model in PhyloBayes. Two calibration points are the origin of *Schistosoma* (66 -145 Ma) and *Paragonimus* genera (7 - 44 Ma) (Vainutis, et al. 2022). The 95% confidence intervals for the time points when the two sister branches diverged are shown as shaded lines at nodes. The colored legend below the timeline corresponds to major geological eras (Cre is Cretaceous; Jur is Jurrasic, Tri is Triassic).

**Figure 3.**
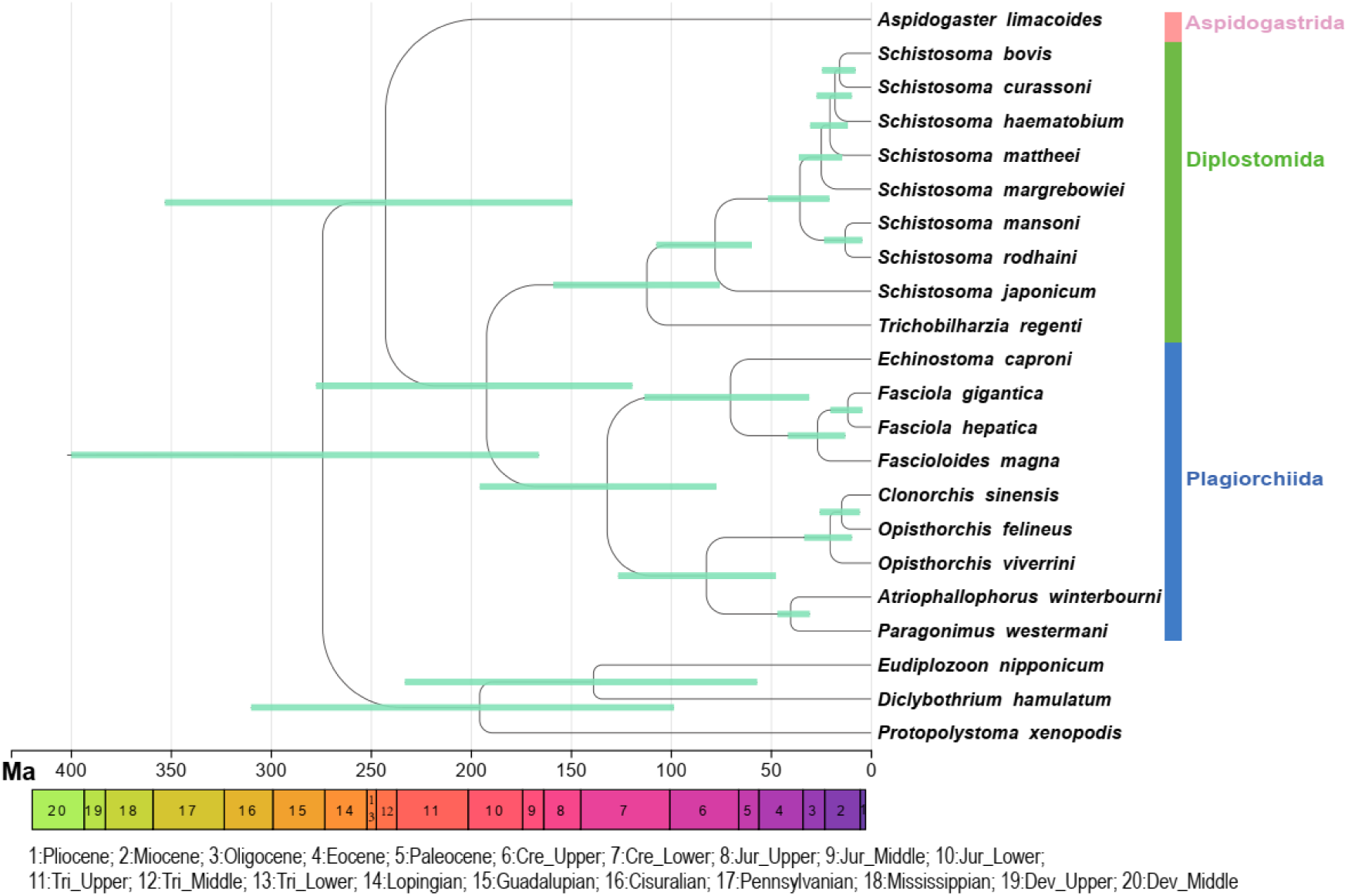
The time tree reconstructed using the AAn-T topology (the dataset comprising nuclear genomes of Trematoda). Two calibration points are the origin of *Schistosoma* (66-145 Ma) and Fasciolidae (50 Ma). The 95% confidence intervals for the time points when the two sister branches diverged are shown as shaded lines at nodes. The colored legend below the timeline corresponds to major geological eras (Cre is Cretaceous; Jur is Jurrasic, Tri is Triassic).

### 3.4. Ancestral state reconstruction

First, we tested the performance of the mitogenomic dataset topologies. The “optimal combination” topologies (NUCPBc and AAPBc) produced a somewhat contradictory evolutionary scenario: two intermediate hosts state was the most probable (0.46 – 0.50 probability) state for proto-trematodes, closely followed by 0 (0.35 – 0.40), but 0 hosts was resolved as by far the most likely ancestral state for proto-aspidogastreans (Figure 4; Table 3). Constrained mitogenomic trees with cestodes as the outgroup (NUCc-cons and AAc-cons; Figure 4 and Figure S38) produced a similar result, but in this case, a direct life-cycle (0 intermediate hosts) was a marginally more likely state than the 2 intermediate hosts for proto-trematodes (Table 3). Two intermediate hosts state was the most likely ancestral state for proto-digeneans, and the ancestors of the Diplostomida and Plagiorchiida orders. However, there was also a 30-40% probability of proto-digeneans having a direct life cycle. In the light of contradictory findings for proto-trematodes, we attempted to further increase the coverage of Aspidogastrea by incorporating the mitogenome of *A. limacoides* extracted from its transcriptome into the existing dataset of 55 species, and reconstructed a new constrained tree (NUCc-AL-cons, Figure S19). This analysis corroborated the above result (for details, see Text S1: “Ancestral state reconstruction”).

**Figure 4.**
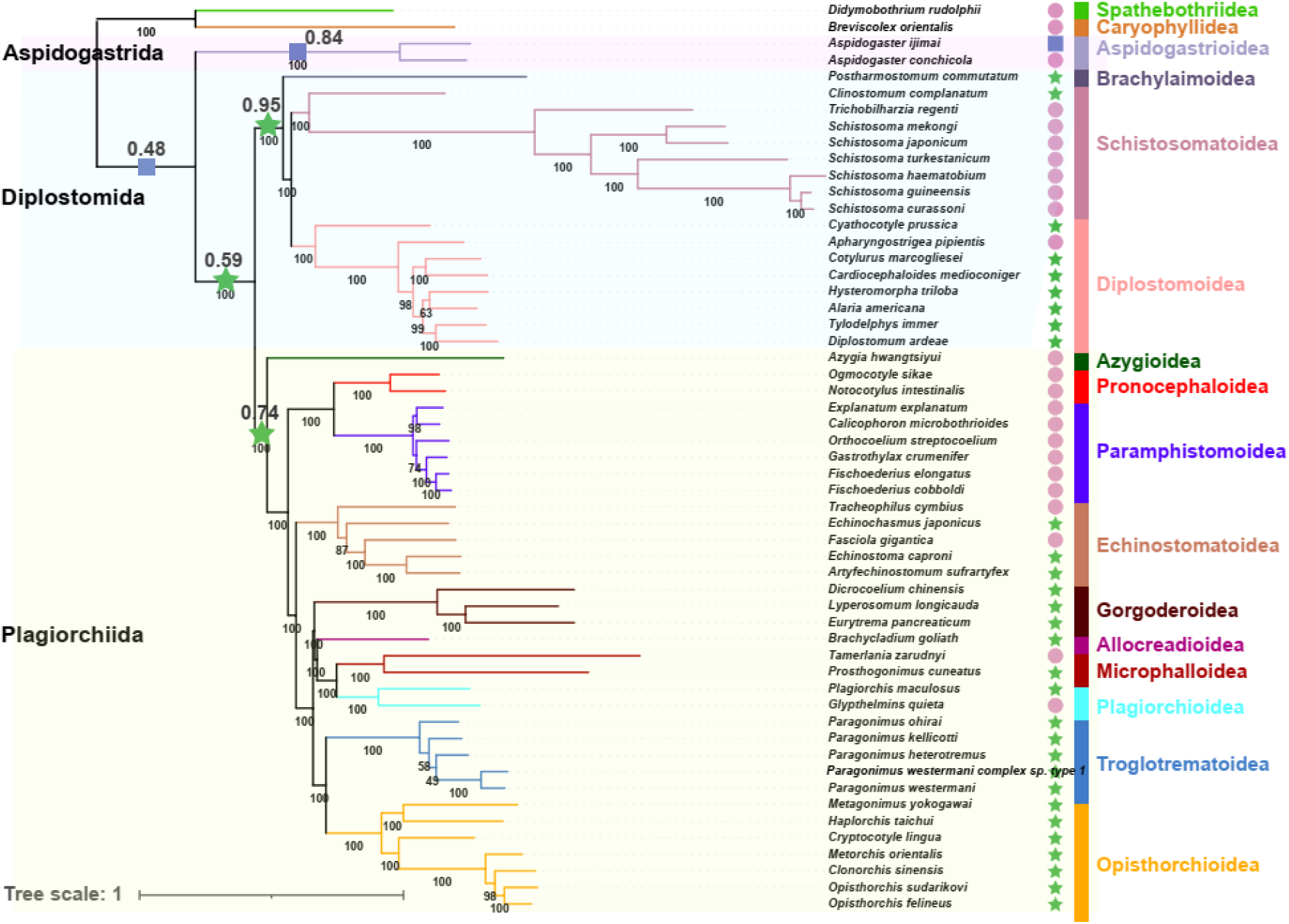
The number of ancestral intermediate hosts inferred using mitogenomic constrained trees with Cestoda as the outgroup and the nucleic sequence dataset (NUCc-cons). The blue square represents the absence of an intermediate host; the purple circles represent one intermediate host; and the green stars represent two intermediate hosts. Bayesian posterior probability values are shown under the branches. The values next to these symbols represent the probability of the most likely ancestral state.

**Table 3.**
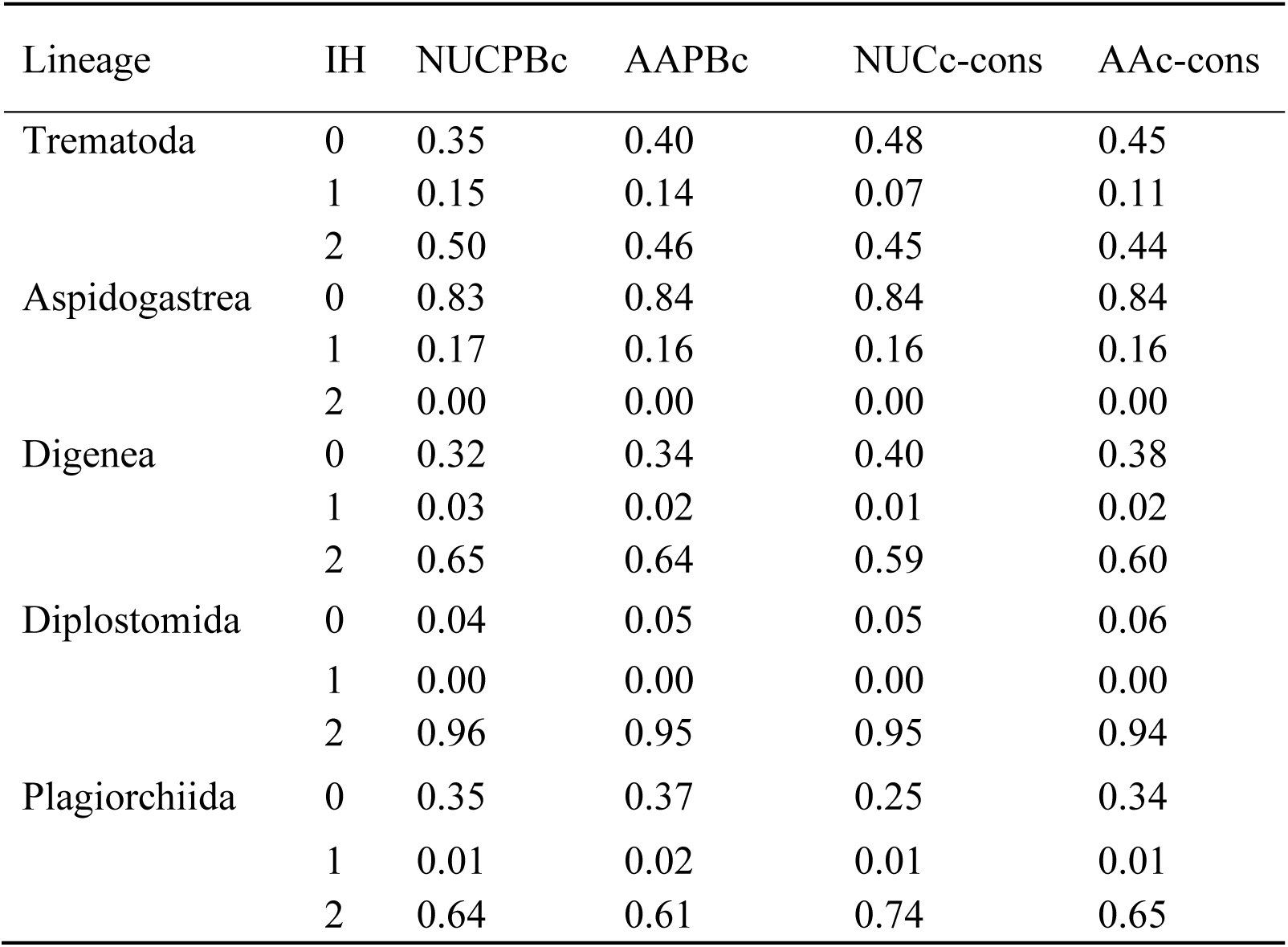
The probabilities of different ancestral states of the intermediate host number for major lineages of Trematoda across different topologies. IH represents the number of intermediate hosts, and tree names are explained in Table 1.

Following this, we tested the performance of the two topologies of Neodermata inferred by Brabec, et al. (2023) using nuclear genomic data (AAIQ-IQn and AAPB-IQn; Figure 5 and Figure S39). When the number of intermediate hosts of the “ancestral aspidogastrean” was 0, the number of intermediate hosts of proto-trematodes was 0; and when the number of intermediate hosts of the “ancestral aspidogastrean” was 1, the number of intermediate hosts of proto-trematodes was 1 (both with 1.0 probability). Therefore, the ancestral state of proto-trematodes was always in agreement with the ancestral state of aspidogastreans. In both cases, the ancestral state was 0 for Neodermata and 1 for Cestoda with a high probability (> 0.96) (“ancestor_Neodermata_Trematoda” in Dataset S2).

**Figure 5.**
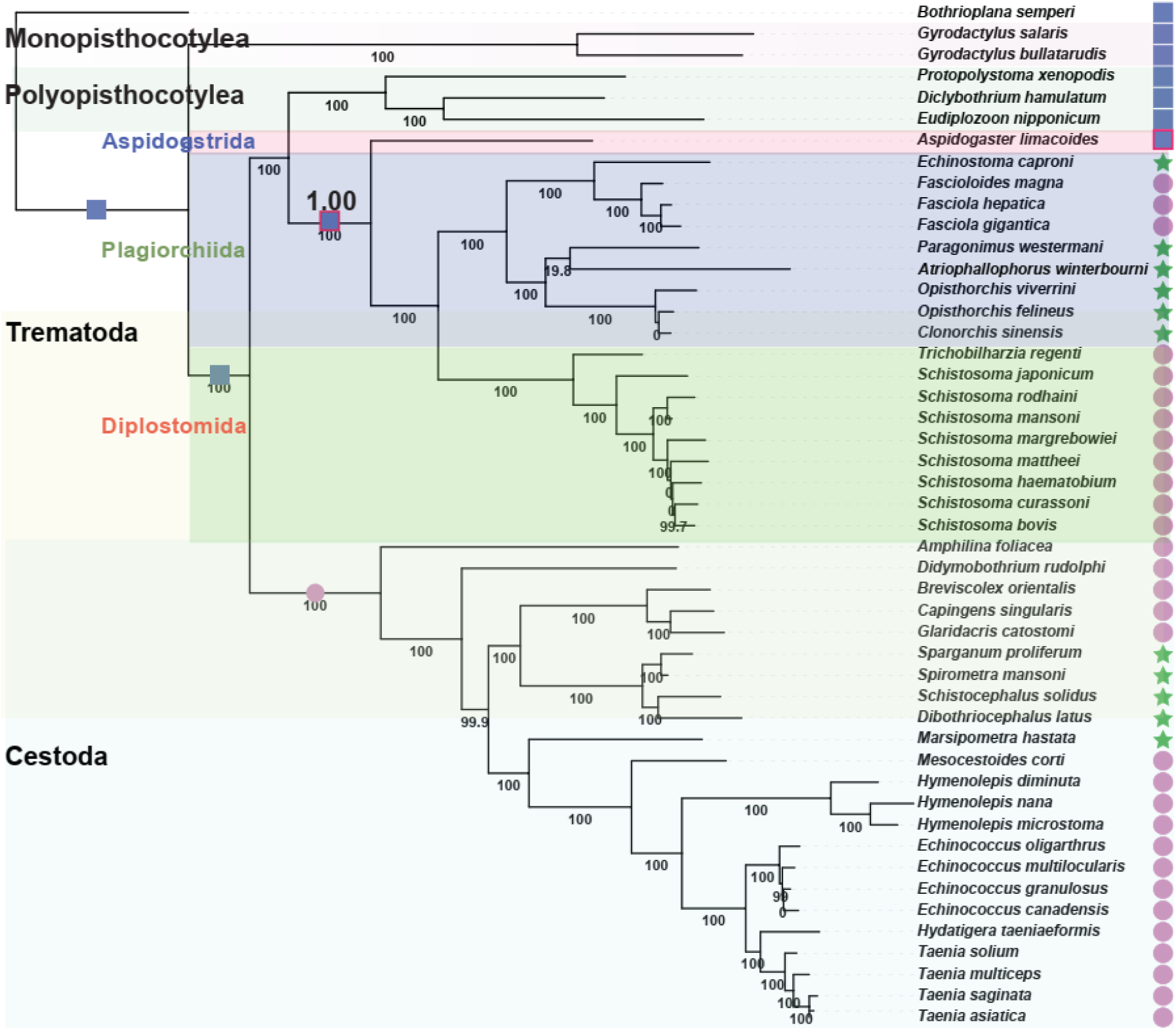
The number of ancestral intermediate hosts inferred on the basis of nuclear genomic (nDNA) topologies constrained according to Brabec, et al. (2023) and reconstructed by IQ-TREE (AAIQ-IQn), with the number of intermediate hosts of the “ancestral aspidogastrean” set to 0. The blue squares represent an absence of an intermediate host; the purple circles represent one intermediate host; and the green stars represent two intermediate hosts. Red borders around symbols highlight the lineages where the ancestral state is identical in proto-trematodes and proto-aspidogastreans. Bayesian posterior probability values are shown under the branches. The values next to these symbols represent the probability of the most likely ancestral state.

## 4. Discussion

### 4.1. Indications of long-branch attraction artefacts in mitogenome-based Trematoda phylogeny with Monopisthocotylea as outgroup support the “basal Aspidogastrea” hypothesis

A portion of topologies inferred with Monopisthocotylea (non-partitioned NUC dataset and all AA dataset analyses with homogeneous models) and Polyopisthocotylea (both with homogeneous and heterogeneous models) as the outgroup resolved Aspidogastrea nested within the Digenea, rendering it paraphyletic, which we explained as a typical long branch attraction phenomenon. However, mitochondrial topologies with Cestoda as the outgroup and all nuclear datasets (the total dataset and three groups of genes classified by function) all supported the “basal Aspidogastrea” hypothesis. To further explore this contradiction, we compared these topologies to constrained topologies, reconstructed on the basis of superfamily relationships reported in previous studies (for details, see “Materials and Methods”), and found that topologies supporting the “basal Aspidogastrea” were consistently more similar to the constraint trees (Dataset S2: sheet “ete3”). In addition, the sCF values also supported the basal position of Aspidogastrea.

Accordingly, the outgroup selection had a major impact on the phylogenetic reconstruction in Trematoda using mitogenomic data. We hypothesize that the use of Monopisthocotylea as the outgroup produced a long-branch attraction artefact (LBA) in phylogenetic tree reconstruction, whereas the use of Cestoda as the outgroup produced the “true” topology, with basal Aspidogastrea. In further support of this hypothesis, *G. salaris* (Monopisthocotylea) was identified as a spurious species and exhibited high long-branch scores in “monogenean” outgroup topologies, but no spurious species were identified when Cestoda were used as the outgroup. The inclusion of spurious species in phylogenetic reconstruction can easily destabilize topologies (Reitmeier, et al. 2021). Another lineage exhibiting high long-branch scores was Schistosomatoidea. As high long-branch scores increase the probability of LBA (Mai and Mirarab 2018), this indicates that mitogenomic topologies in which Schistosomatoidea was resolved as the basal lineage are likely to be a result of long-branch attraction between Schistosomatoidea and Monopisthocotylea.

### 4.2. Resolving topological instability: data heterogeneity and the optimal evolutionary model

Schistosomatoidea also exhibited relatively high compositional heterogeneity - all species failed the homogeneity test in IQ-TREE. As several previous studies reported that heterogeneous models perform better than homogeneous models on datasets exhibiting strong compositional biases (Coghlan, et al. 2019; Uribe, et al. 2019), we hypothesized that compositional heterogeneity of the dataset was also a major factor contributing to the long-branch attraction when using homogeneous models and Monopisthocotylea as the outgroup. To further validate this hypothesis, we conducted symmetry tests to detect and remove partitions that violated the assumptions of stationarity and homogeneity (Naser-Khdour, et al. 2019). These pruned datasets all produced basal Aspidogastrea and monophyletic Diplostomida, regardless of the outgroup and partitioning strategies. However, Plagiorchiida was still paraphyletic in topologies reconstructed based on the pruned AAc-passed dataset. We speculate that the reason may be that the symmetry test removed the data carrying crucial phylogenetic signal in the AAc dataset.

MAST was used to evaluate the reliability of topologies by calculating their weight values (Minh, et al. 2020). For NUCm, NUCc and AAm datasets, topologies reconstructed by the heterogeneous model of PhyloBayes had the highest weight values. For the AAc dataset, the highest weight was exhibited by the topology inferred using the heterogeneous model implemented in IQ-TREE (mtInv+C50+F+R6). However, unlike the heterogeneous model in PhyloBayes (CAT-GTR), mtInv+C50+F+R6 produced paraphyletic Diplostomida (AAIQc-He topology). As MAST analysis is included in the IQ-TREE platform, we hypothesize that the underlying reason for this discrepancy might be that CAT-GTR was not supported by MAST, so mtInv+C50+F+R6 was used to test the topology produced using CAT-GTR in PhyloBayes (for details, see “Materials and Methods”). In summary, our results suggested that Trematoda mitogenomes evolve in a heterogeneous manner and that CAT-GTR (PhyloBayes) performed better than all other models tested in this study.

### 4.3. The evolutionary history of parasitic strategies in the class Trematoda and divergence times

Mitogenomic topologies (constrained and “optimal”) produced somewhat inconsistent results, indicating that 0 and 2 intermediate hosts are approximately equally likely as the ancestral state for Trematoda. Nuclear genomic topologies indicated that the ancestral state of aspidogastreans was always consistent with that of proto-trematodes (0 or 1; regardless of the topology), and mitogenomic data consistently resolved 0 as the most likely ancestral state for aspidogastreans. As no known aspidogastreans have two intermediate hosts, and 0 intermediate hosts was resolved as the most likely ancestral state for Neodermata, we hypothesise that 2 intermediate hosts in the ancestral trematodes was an artefact caused by the strong dominance of digeneans (the most likely ancestral state = 2) in the dataset. On this basis, we infer that the best-supported scenario is the absence of intermediate hosts in both proto-trematodes and proto-aspidogastreans. This supports the scenario wherein proto-trematodes first parasitized mollusks (probably mainly bivalves), and then gradually evolved the ability to parasitize (arthropods and) vertebrates, whereas mollusks gradually became intermediate hosts. The latter scenario also partially supports the speculation of Gibson and Bray (1994). However, our analyses indicate that the ancestral number of intermediate hosts in digeneans is two, which opens a logical gap between the 0 intermediate hosts in proto-trematodes and 2 in the ancestral digeneans. Either our ancestral state reconstruction was wrong, and the true ancestral state for digeneans is 1, or we must assume that the ancestral digenean lineage underwent a relatively long evolutionary period before radiating into the modern digeneans. This would have allowed a stepwise evolution to one intermediate host in early proto-digeneans, and eventually two intermediate hosts in late proto-digeneans. This scenario is also more parsimonious as it requires fewer independent host losses than the required independent host gains in the scenario where the ancestral state for Digenea was 1: 8 vs. 11 (in the mitogenomic phylogram).

During the Jurassic period (200 to 150 Ma), mollusks were abundant and their migration rates were high (Hallam 1976; Crame 1993), which created favorable conditions for their potential parasites and predators (Abrams and Ginzburg 2000; Canard, et al. 2014). In addition, roughly during this period (Triassic and Jurassic), various orogenic movements (such as the Cimmerian orogeny) created novel geographical barriers (Golonka, et al. 2003), which facilitated taxonomic radiations. Therefore, host abundance and geographic isolation may have created a favorable environment for the divergence of proto-trematodes, which possessed a direct life cycle, relying solely on a single host (bivalves) for reproduction through strict sexual means, devoid of asexual generations in our hypothetical evolutionary trajectory. During the Late Triassic and/or Early Cretaceous, the ancestor of the digenean lineage switched to gastropods as hosts, facilitating the bifurcation of proto-trematodes into proto-aspidogastreans and proto-digeneans. Both types of hosts (gastropods and bivalves) were preyed upon by the lower vertebrates (predominantly fish), so both lineages had ample opportunities to evolve the ability to infect fish, thus eventually resulting in a one-intermediate-host parasitic lifestyle. Whereas only some aspidogastreans evolved this ability, it became the obligatory parasitic strategy in early proto-digeneans. As fish were commonly preyed upon by higher vertebrates, this further provided ample opportunities to infect a new definitive host, which eventually allowed late proto-digeneans to develop the obligatory two-intermediate hosts parasitic strategy, mainly with gastropods as the first intermediate host, fish (lower vertebrates) as the second, and higher vertebrates (birds/mammals/reptiles) as the definitive host.

This raises the question: why did aspidogastreans (on average) fail to develop more complex parasitic strategies if they had similar opportunities as digeneans? A parameter that may explain why the evolutionary trajectories differed between digeneans and aspidogastreans might be the reproduction mode. Unlike digeneans, aspidogastreans do not have an asexual generation (Huehner and Etges 1977). Accordingly, we assume that sexual reproduction is the ancestral state, and alternating sexual and asexual generations is a derived state in Trematoda. The number of offspring per adult produced in asexual reproduction is larger than in sexual reproduction: a miracidium hatched from an egg can multiply into hundreds of cercariae during its growth period in the intermediate host (Bengtsson 2003). For parasites, host transmission via the food chain is associated with multiple challenges, such as the competition with other parasites within the new host (Thompson, et al. 2013), and the fact that the intermediate host might be killed and eaten by a predator that is not a suitable definitive host (Cirtwill, et al. 2017). As a result of these large costs, lineages that lacked the ability of asexual reproduction, such as proto-aspidogastreans, did not form the two-intermediate-hosts life history and generally have zero or only one intermediate hosts. This also implies that the development of asexual reproduction may have been a key step in the evolution of proto-digeneans, allowing them to fully switch to parasitic strategies with multiple intermediate hosts.

As the major Cretaceous-Paleogene extinction event (about 66 Ma) resulted in the prevalence of mammals and birds over reptiles in subsequent eras (Schulte, et al. 2010), digeneans are currently more prevalent in birds and mammals than in reptiles. This abundance of definitive hosts may also have facilitated the subsequent loss of one intermediate host: cercariae released from the first intermediate host could frequently come into direct contact with the definitive hosts through drinking water and herbivory (e.g. Paramphistomoidea) (Jones 2005) or through wounds and reproductive organs (e.g. *Schistosoma*) (Horák, et al. 2002). This may have allowed the loss of the second intermediate host in some digeneans. To summarize, the sequence of intermediate host strategies in the evolutionary history of Trematoda follows four patterns, two in Aspidogastrea: 0 – 0 – 0/1 (0 in proto-trematodes – 0 in proto-aspidogastreans – 0 or 1 in contemporary aspidogastreans); and two in Digenea: 0 – 1 – 2 – 2/1 (0 in proto-trematodes – 1 in early proto-digeneans – 2 in late proto-digeneans – 2 or 1 in contemporary digeneans).

### 4.4. Limitations

Our phylogenetic analyses resulted in pronounced topological instability. Various parameters affected the topology, including datasets, outgroups, partitioning strategies, evolutionary models, etc. While we found the strongest support for topologies that resolved all major Trematoda lineages as monophyletic, there exists a certain risk of circular reasoning in our arguments. Similarly, there is currently no perfect algorithm for the time tree inference and ancestral state reconstruction, and both can be strongly affected by the topology, branch lengths, and outgroups (Litsios and Salamin 2012; Meade and Pagel 2022). Indeed, confidence intervals were very wide in time tree inference, and results differed between mitogenomic and nuclear genomic topologies. In addition, due to the scarcity of fossil evidence for these lineages, calibration points used for molecular dating may lack precision. The discrepancy was also likely to be caused by the severely limited sampling of aspidogastreans. As the small size of aspidogastrean parasites presents a major challenge for nuclear genome sequencing, currently there is only one transcriptome available for this lineage. We found that the state of this species had a major impact on the ancestral state reconstruction. In addition, results were inconsistent across different topologies for proto-trematodes and proto-aspidogastreans. In the mitogenomic dataset, we had only two (three with *A. limacoides*) aspidogastrean mitogenomes at disposal, comprising two different life histories, which offers a stronger resolution. However, this is only a small proportion of the valid 64 aspidogastrean species (WoRMS 2024). The sampling of Digenea was somewhat better: the 51 mitogenomes in our dataset covered 22 out of 54 valid families, 13 out of 20 superfamilies, and both valid orders (Diplostomida and Plagiorchiida). Once data are available for more lineages, our results should be further confirmed.

## 5. Conclusions

Our study offers first insights into the evolution of intermediate host number in Trematoda. We based the analyses on both mitogenomic and nuclear genomic topologies. For this, we sequenced mitogenomes of first two aspidogastrean species and applied various strategies to tackle the topological instability hampering the phylogenetic reconstruction in trematodes. The constraint trees with Cestoda as the outgroup were identified as the optimal topologies for ancestral state reconstruction analyses. These revealed that a direct life cycle, devoid of intermediate hosts, was probably the ancestral state for both Trematoda and Aspidogastrea, whereas the ancestor of Digenea likely utilized two intermediate hosts. Additionally, we propose a timeline outlining these evolutionary shifts and the different evolutionary pathways of intermediate host number in Aspidogastrea and Digenea. Finally, our analyses indicate that host strategies are relatively plastic among trematodes, putatively comprising several independent host gains, and multiple host losses.

## Supporting information

Supplementary files for The Phylogeny and the Evolution of Parasitic Strategies in Trematoda

BI: Bayesian Inference method
ML: Maximum Likelihood method
aBSREL: adaptive Branch-Site Random Effects Likelihood
RCV: relative composition variability
ESS: effective sample size
LBA: long-branch attraction artefact
gCF: gene concordance factor
sCF: site concordance factor

## CRediT authorship contribution statement

Conceptualization: CYX, DZ, IJ, GTW. Methodology and Formal analysis: CYX, IJ, TY, and DZ.; Investigation: RS, HZ, WXL, and GTW; Resources: WXL, GTW; Validation: CYX, DZ, IJ. Writing – original draft: CYX; Writing – review and editing: all authors; Visualization: CYX; Supervision: DZ, IJ, WXL, GTW; Funding acquisition: DZ.

## Data availability

The two newly sequenced mitogenomes are deposited in GenBank under the accession numbers: PP903627 (*Aspidogaster ijimai*) and PP903628 (*Aspidogaster conchicola*). The GenBank accession numbers of other mitogenomes used in our analyses are available in Dataset S1. We also provide the GenBank files of the two *Aspidogaster* species (GenBank S1 for *Aspidogaster ijimai*; GenBank S2 for *Aspidogaster conchicola*)

## Acknowledgments

The authors thank Dr. Fu Peipei, Dr. Liu Xinhua and Dr. Xi Bingwen for their contributions to sample collection.

## Supplementary Material

Supplementary Text and Tables, Supplementary Figures S1-S52, GenBank S1-S2, and Supplementary Datasets S1-S3 can be accessed via prepublication Dryad link: https://doi.org/10.5061/dryad.qnk98sfrh.

## References

Abrams PA, Ginzburg LR. 2000. The nature of predation: prey dependent, ratio dependent or neither? Trends in Ecology Evolution 15:337–341.

Alves PV, Vieira FM, Santos CP, Scholz T, Luque JL. 2015. A checklist of the aspidogastrea (platyhelminthes: trematoda) of the world. Zootaxa 3918:339–396. doi: 10.11646/zootaxa.3918.3.2.

Atopkin DM, Shedko MB, Sokolov SG, Zhokhov AE. 2018. Phylogenetic relationships among European and Asian representatives of the genus Aspidogaster Baer, 1827 (Trematoda: Aspidogastrea) inferred from molecular data. J Helminthol 92:343–352. doi: 10.1017/S0022149X17000505.

Bengtsson BO. 2003. Genetic variation in organisms with sexual and asexual reproduction. Journal of evolutionary biology 16:189–199.

Brabec J, Salomaki ED, Kolísko M, Scholz T, Kuchta R. 2023. The evolution of endoparasitism and complex life cycles in parasitic platyhelminths. Curr Biol. doi: 10.1016/j.cub.2023.08.064. Brooks DR, O’Grady RT, Glen DR. 1985. Phylogenetic analysis of the Digenea (Platyhelminthes: Cercomeria) with comments on their adaptive radiation. 63:411–443. doi: 10.1139/z85-062.

Canard E, Mouquet N, Mouillot D, Stanko M, Miklisova D, Gravel D. 2014. Empirical evaluation of neutral interactions in host-parasite networks. The American Naturalist 183:468–479.

Carlson CJ, Dallas TA, Alexander LW, Phelan AL, Phillips AJ. 2020. What would it take to describe the global diversity of parasites? Proceedings of the Royal Society B 287:20201841.

Chen M-X, Zhang L-Q, Wen C, Sun J, Gao Q. 2010. Phylogenetic Relationship of Species in the Genus Aspidogaster (Aspidogastridae, Aspidogastrinae) in China as Inferred from Its rDNA Sequences. Acta Hydrobiologica Sinica 34:312–316. doi: 10.3724/SP.J.1035.2009.00312.

Cirtwill AR, Lagrue C, Poulin R, Stouffer DB. 2017. Host taxonomy constrains the properties of trophic transmission routes for parasites in lake food webs. Ecology 98:2401–2412.

Coghlan A, Tyagi R, Cotton JA, Holroyd N, Rosa BA, Tsai IJ, Laetsch DR, Beech RN, Day TA, Hallsworth-Pepin K, et al. 2019. Comparative genomics of the major parasitic worms. Nature Genetics 51:163–174. doi: 10.1038/s41588-018-0262-1.

Crame JA. 1993. Bipolar molluscs and their evolutionary implications. Journal of Biogeography:145–161.

Cribb T, Bray R, Littlewood D. 2001. The nature and evolution of the association among digeneans, molluscs and fishes. International Journal for Parasitology 31:997–1011.

dos Reis M, Yang Z. 2017. MCMCTree tutorials. In.

Ferguson M, Cribb T, Smales L. 1999. Life-cycle and biology of Sychnocotyle kholo ng, n. sp.(Trematoda: Aspidogastrea) in Emydura macquarii (Pleurodira: Chelidae) from southern Queensland, Australia. Systematic Parasitology 43:41–48.

Gibson DI, Bray RA. 1994. The evolutionary expansion and host-parasite relationships of the Digenea. Int J Parasitol 24:1213–1226. doi: 10.1016/0020-7519(94)90192-9.

Giese EG, Silva MVO, Videira MN, Furtado AP, Matos ER, Gonçalves EC, Melo FTV, Santos JN. 2015. Rohdella amazonica n. sp. (Aspidogastrea: Aspidogastridae) from the Amazoninan banded puffer fish Colomesus psittacus (Bloch & Schneider, 1801). J Helminthol 89:288–293. doi: 10.1017/S0022149X14000054.

Golonka J, Krobicki M, Oszczypko N, Ślączka A, Słomka T. 2003. Geodynamic evolution and palaeogeography of the Polish Carpathians and adjacent areas during Neo-Cimmerian and preceding events (latest Triassic-earliest Cretaceous). Geological Society, London, Special Publications 208:137–158.

Guindon S. 2020. Rates and rocks: strengths and weaknesses of molecular dating methods. Frontiers in Genetics 11:493652.

Hallam A. 1976. Stratigraphic distribution and ecology of European Jurassic bivalves. Lethaia 9:245–259.

Hanson-Smith V, Kolaczkowski B, Thornton JW. 2010. Robustness of ancestral sequence reconstruction to phylogenetic uncertainty. Molecular biology and evolution 27:1988–1999.

Helden AJ, Dixon A. 2002. Life-cycle variation in the aphid Sitobion avenae: costs and benefits of male production. Ecological Entomology 27:692–701.

Horák P, Kolářová L, Adema C. 2002. Biology of the schistosome genus Trichobilharzia.

Huehner MK, Etges FJ. 1977. The Life Cycle and Development of Aspidogaster conchicola in the Snails, Viviparus malleatus and Goniobasis livescens. The Journal of Parasitology 63:669–674. doi: 10.2307/3279567.

Huyse T, Buchmann K, Littlewood D. 2008. The mitochondrial genome of Gyrodactylus derjavinoides (Platyhelminthes: Monogenea)—a mitogenomic approach for Gyrodactylus species and strain identification. Gene 417:27–34.

Jones A. 2005. Superfamily Paramphistomoidea Fischoeder, 1901. In. Keys to the Trematoda: Volume 2: CABI Publishing Wallingford UK. p. 221–227.

Jourdane J, Mingyi X. 1987. The primary sporocyst stage in the life cycle of Schistosoma japonicum (Trematoda: Digenea). Transactions of the American Microscopical Society:364–372.

Justine J-L. 1998. Non-monophyly of the monogeneans? International Journal for Parasitology 28:1653–1657.

Kalyaanamoorthy S, Minh BQ, Wong TKF, von Haeseler A, Jermiin LS. 2017. ModelFinder: fast model selection for accurate phylogenetic estimates. Nature Methods 14:587–589. doi: 10.1038/nmeth.4285.

Kenny NJ, Noreña C, Damborenea C, Grande C. 2019. Probing recalcitrant problems in polyclad evolution and systematics with novel mitochondrial genome resources. Genomics 111:343–355. doi: 10.1016/j.ygeno.2018.02.009.

Kim D. 1984. Paragonimus westermani: life cycle, intermediate hosts, transmission to man and geographical distribution in Korea. Arzneimittel-forschung 34:1180–1183.

Kosakovsky Pond SL, Poon AFY, Velazquez R, Weaver S, Hepler NL, Murrell B, Shank SD, Magalis BR, Bouvier D, Nekrutenko A, et al. 2019. HyPhy 2.5—A Customizable Platform for Evolutionary Hypothesis Testing Using Phylogenies. Molecular biology and evolution 37:295–299. doi: 10.1093/molbev/msz197%J Molecular Biology and Evolution.

Lanfear R, Welch JJ, Bromham L. 2010. Watching the clock: studying variation in rates of molecular evolution between species. Trends in Ecology Evolution 25:495–503.

Li Y, Ma XX, Lv QB, Hu Y, Qiu HY, Chang QC, Wang CR. 2020. Characterization of the complete mitochondrial genome sequence of Tracheophilus cymbius (Digenea), the first representative from the family Cyclocoelidae. J Helminthol 94:e101. doi: 10.1017/S0022149X19000932.

Litsios G, Salamin N. 2012. Effects of Phylogenetic Signal on Ancestral State Reconstruction. Systematic Biology 61:533–538. doi: 10.1093/sysbio/syr124.

Lotfy WM, Brant SV, DeJong RJ, Le TH, Demiaszkiewicz A, Rajapakse RP, Perera VB, Laursen JR, Loker ES. 2008. Evolutionary origins, diversification, and biogeography of liver flukes (Digenea, Fasciolidae). Am J Trop Med Hyg 79:248–255.

Mai U, Mirarab S. 2018. TreeShrink: fast and accurate detection of outlier long branches in collections of phylogenetic trees. BMC genomics 19:23–40.

Meade A, Pagel M. 2022. Ancestral state reconstruction using BayesTraits. In. Environmental Microbial Evolution: Methods and Protocols: Springer. p. 255–266.

Minh BQ, Schmidt HA, Chernomor O, Schrempf D, Woodhams MD, von Haeseler A, Lanfear R. 2020. IQ-TREE 2: New Models and Efficient Methods for Phylogenetic Inference in the Genomic Era. Mol Biol Evol 37:1530–1534. doi: 10.1093/molbev/msaa015.

Naser-Khdour S, Minh BQ, Zhang W, Stone EA, Lanfear R. 2019. The prevalence and impact of model violations in phylogenetic analysis. Genome biology evolution 11:3341–3352.

Olson PD, Cribb TH, Tkach VV, Bray RA, Littlewood DTJ. 2003. Phylogeny and classification of the Digenea (Platyhelminthes: Trematoda)11Nucleotide sequence data reported in this paper are available in the GenBank™, EMBL and DDBJ databases under the accession numbers AY222082–AY222285. International Journal for Parasitology 33:733–755. doi: 10.1016/S0020-7519(03)00049-3.

Parfrey LW, Lahr DJ, Knoll AH, Katz LA. 2011. Estimating the timing of early eukaryotic diversification with multigene molecular clocks. Proceedings of the National Academy of Sciences 108:13624–13629.

Peterson KJ, Lyons JB, Nowak KS, Takacs CM, Wargo MJ, McPeek MA. 2004. Estimating metazoan divergence times with a molecular clock. Proceedings of the National Academy of Sciences 101:6536–6541.

Reed P, Francis-Floyd R, Klinger R, Petty D. 2009. Monogenean parasites of fish. Fisheries aquatic sciences. University of Florida UF, IFAS Extension. FA28, USA 4:1–4.

Reitmeier S, Hitch TCA, Treichel N, Fikas N, Hausmann B, Ramer-Tait AE, Neuhaus K, Berry D, Haller D, Lagkouvardos I, et al. 2021. Handling of spurious sequences affects the outcome of high-throughput 16S rRNA gene amplicon profiling. ISME Communications 1:31. doi: 10.1038/s43705-021-00033-z.

Rohde K. 1971. Phylogenetic origin of trematodes. Parasitologische Schriftenreihe 21:17–27.

Rubinoff D, Holland BS. 2005. Between Two Extremes: Mitochondrial DNA is neither the Panacea nor the Nemesis of Phylogenetic and Taxonomic Inference. Systematic Biology 54:952–961. doi: 10.1080/10635150500234674.

Schulte P, Alegret L, Arenillas I, Arz JA, Barton PJ, Bown PR, Bralower TJ, Christeson GL, Claeys P, Cockell CS. 2010. The Chicxulub asteroid impact and mass extinction at the Cretaceous-Paleogene boundary. Science 327:1214–1218.

Smyth JD, McManus DP. 1989. The physiology and biochemistry of cestodes: Cambridge university press.

Suleman, Ma J, Khan MS, Tkach VV, Muhammad N, Zhang D, Zhu X-Q. 2019. Characterization of the complete mitochondrial genome of Plagiorchis maculosus (Digenea, Plagiorchiidae), Representative of a taxonomically complex digenean family. Parasitology International 71:99–105. doi: 10.1016/j.parint.2019.04.001.

Suleman, Muhammad N, Khan MS, Tkach VV, Ullah H, Ehsan M, Ma J, Zhu X-Q. 2021. Mitochondrial genomes of two eucotylids as the first representatives from the superfamily Microphalloidea (Trematoda) and phylogenetic implications. Parasites & Vectors 14:48. doi: 10.1186/s13071-020-04547-8.

Tang Chongti TZ. 2015. Chinese Trematology: Science Press.

Thompson RM, Poulin R, Mouritsen KN, Thieltges DW. 2013. Resource tracking in marine parasites: going with the flow? Oikos 122:1187–1194.

Tice AK, Žihala D, Pánek T, Jones RE, Salomaki ED, Nenarokov S, Burki F, Eliáš M, Eme L, Roger AJ, et al. 2021. PhyloFisher: A phylogenomic package for resolving eukaryotic relationships. PLoS Biol 19:e3001365. doi: 10.1371/journal.pbio.3001365.

Uribe JE, Irisarri I, Templado J, Zardoya R. 2019. New patellogastropod mitogenomes help counteracting long-branch attraction in the deep phylogeny of gastropod mollusks. Molecular phylogenetics evolution 133:12–23.

Vainutis KS, Voronova AN, Duscher GG, Shchelkanov EM, Shchelkanov MY. 2022. Origins, phylogenetic relationships and host-parasite interactions of Troglotrematoidea since the cretaceous. Infection, Genetics Evolutionary Bioinformatics 101:105274.

Wong TK, Cherryh C, Rodrigo AG, Hahn MW, Minh BQ, Lanfear RJb. 2022. MAST: Phylogenetic Inference with Mixtures Across Sites and Trees.

Aspidogastrea [Internet]. 2024. cited Accessed at: https://www.marinespecies.org/aphia.php?p=taxdetails&id=103973 on 2024-07-15].

Wright C. 1967. The schistosome life-cycle. In. Bilharziasis: International Academy of Pathology· Special Monograph: Springer. p. 3–7.

Yuan ML, Zhang LJ, Zhang QL, Zhang L, Li M, Wang XT, Feng RQ, Tang PA. 2020. Mitogenome evolution in ladybirds: Potential association with dietary adaptation. Ecology Evolution 10:1042–1053.

Zhang D, Jakovlić I, Zou H, Liu F, Xiang CY, Gusang Q, Tso S, Xue S, Zhu WJ, Li Z, et al. 2024. Strong mitonuclear discordance in the phylogeny of Neodermata and evolutionary rates of Polyopisthocotylea. Int J Parasitol. doi: 10.1016/j.ijpara.2024.01.001.

Zhang D, Zou H, Hua C-J, Li W-X, Mahboob S, Al-Ghanim KA, Al-Misned F, Jakovlić I, Wang G-T. 2019. Mitochondrial Architecture Rearrangements Produce Asymmetrical Nonadaptive Mutational Pressures That Subvert the Phylogenetic Reconstruction in Isopoda. Genome Biology and Evolution 11:1797–1812. doi: 10.1093/gbe/evz121.

Zhang J. 1999. Fish Parasites and Parasitic Diseases: Science Press.

