## Supplementary files for The Phylogeny and the Evolution of Parasitic Strategies in Trematoda for "The Phylogeny and the Evolution of Parasitic Strategies in Trematoda": Text S1.docx

### Supplementary Materials and Methods

#### DNA extraction, Amplification and Sequencing

The study used two types of DNA for amplification and sequencing of small parasite specimens - mixture DNA (pooled multiple specimens) and individual DNA (a single specimen). Degenerate primer pairs were designed to amplify and sequence selected mitochondrial gene regions, followed by the use of specific primers, design to amplify and sequence the whole mitogenome in several PCR steps. The PCR products were sequenced bi-directionally using the Sanger method and analyzed for potential sequence variations using BLAST (Altschul, Gish et al. 1990). Intraspecific sequence variation was addressed using individual DNA and long-range PCR.

#### Sequence Annotation and Analyses

The mitogenomic sequences of *A. ijimai* and *A. conchicola* were manually assembled and annotated using various programs, including MITOS (Bernt, Donath et al. 2013), ARWEN (Laslett and Canbäck 2008). Codon usage and relative synonymous codon usage (RSCU) were calculated for twelve protein-encoding genes (PCGs) of the two studied aspidogastrids. Non-synonymous (dN)/ synonymous (dS) mutation rate ratios were computed using KaKs_Calculator (Zhang, Li et al. 2006), and nucleotide divergence Pi was estimated between the two mitogenomes using the sliding window analysis with DnaSP v5 (Librado and Rozas 2009). Tandem Repeats Finder (Benson 1999) was used to identify tandem repeats in non-coding regions, and their secondary structures were predicted by Mfold (Zuker 2003). Genetic distances among mitogenomic sequences were calculated using the “DistanceCalculator” function in Biopython (Cock, Antao et al. 2009).

#### Phylogenetic Analyses

Phylogenetic analyses were conducted using the two newly sequenced Aspidogastrea mitogenomes and 51 Trematoda mitogenomes available in the NCBI’s GenBank database (last accessed on 2022.6.7.). *Benedenia seriolae* and *Gyrodactylus salaris* (“monogenea” or Monopisthocotylea) and *Didymobothrium rudolphii* and *Breviscolex orientalis* (Cestoda) were tested as outgroups, thus adding up to 55 mitogenomes in each group. Besides, as Polyopisthocotylea were the sister clade to Trematoda in a recent nuclear genome-based phylogenetic reconstruction (Brabec, Salomaki et al. 2023), we also tested the performance of two Polyopisthocotylea species as outgroups for phylogenetic reconstruction (for details, see “Phylogenetic analyses” and “Phylogenetic analysis with Polyopisthocotylea as the outgroup” in Supplementary Text). The genes were extracted from GenBank files using PhyloSuite (Zhang, Gao et al. 2020, Xiang, Gao et al. 2023) as described before (Zhang, Zou et al. 2018, Naser-Khdour, Minh et al. 2019).

In PhyloSuite, nucleotide sequences and amino acid sequences were aligned in batches using MAFFT (Katoh and Standley 2013, Katoh, Standley et al. 2013), and Gblocks 0.91b (Castresana 2000) was used to remove the ambiguously aligned fragments with “relaxed” parameters: minimum number of sequences for a conserved/flank position, maximum number of contiguous non-conserved positions, minimum length of a block, allowed gap positions (with half). The optimized alignments were concatenated into two datasets by PhyloSuite, where AA refers to the supermatrix of amino acid sequences of protein-coding sequences (PCGs), and NUC denotes the supermatrix of nucleotide sequences of PCGs and RNA genes. The most appropriate evolutionary models for the above datasets were inferred using ModelFinder (Kalyaanamoorthy, Minh et al. 2017). Phylogenetic analyses with homogeneous models were conducted using three programs: RAxML (Stamatakis 2014) and two plugins of PhyloSuite: IQ-TREE 1.65 (Nguyen, Shuval et al. 2015) and MrBayes 3.2.6 (Ronquist, Teslenko et al. 2012). IQ-TREE was run with default settings, and the standard bootstrap with 1000 repetitions for three topologies (AAIQm, NUCIQm and NUCIQc), and the ultrafast bootstrap with 5000 repetitions for all other topologies (see reasons for the different settings in the Supplementary Text). MrBayes was also run with default settings and 4×10^6^ metropolis-coupled MCMC generations. Stationarity was considered to be reached when the average standard deviation of split frequencies was below 0.01, PSRF (potential scale reduction factor) approached 1, and the ESS values were higher than 100. For the “heterogeneous” CAT-GTR model analysis, implemented in PhyloBayes-MPI 1.7a (PB) (Lartillot, Rodrigue et al. 2013), two MCMC chains were run after the removal of invariable sites from the alignment. The analysis was stopped when the conditions considered to indicate a good run (PhyloBayes manual) were reached: maxdiff ≈0.09 (< 0.1) and minimum effective size >300. Phylogenetic trees were visualized and annotated by iTOL (Letunic and Bork 2024) with the help of several dataset files generated by PhyloSuite.

IQ-TREE was run with default settings, and the standard bootstrap with 1000 repetitions for three topologies (AAIQm, NUCIQm and NUCIQc) and ultrafast bootstrap with 5000 repetitions for all other topologies. We changed the parameters because the first three topologies were reconstructed normally, but the standard bootstrap mode failed when Cestoda were used as the outgroup due to a bug in IQ-TREE, whereas the ultrafast mode was successfully run.

We also divided all topologies into four testing groups based on outgroups (monogeneans or cestodes) and datasets (NUC or AA) for basic tree comparison. The basic tree comparison analysis was based on the ete-compare, where higher “%ref_br” (frequency of edges in the reference tree found in target) and “%src_br” (frequency of edges in target tree found in the reference) values and lower “RF” values (Robinson-Foulds symmetric distance) indicated that the tested tree was comparatively consistent with the reference tree (Huerta-Cepas, Serra et al. 2016). As Brabec, Salomaki et al. (2023) resolved Polyopisthocotylea as the sister clade to Trematoda, to further explore the impact of outgroup selection and heterogeneity models on the topology, we used concatenated nucleotide sequences of two Polyopisthocotylea species (*Sphyranura euryceae* and *Eudiplozoon nipponicum*) as the outgroup and reconstructed topologies using homogeneous (IQ-TREE, NUCIQmp) and heterogeneous (PhyloBayes, NUCPBmp) models.

#### References for Topologies and Hosts

References for NUCc-cons and AAc-cons topologies: (Olson, Cribb et al. 2003, Morand 2015, Alba 2018, Li, Ma et al. 2019, Pérez-Ponce de León and Hernández-Mena 2019, Chan, Saralamba et al. 2022). References for the number of intermediate hosts and definitive host species: (Kaw , Srivastava 1944, Maldonado 1945, Leigh 1946, Clarke 1954, Hutchison 1959, Meyer 1960, Sindermann and Farrin 1962, Macy 1965, Yokogawa 1965, Seo and Kwak 1972, Bakker and Davids 1973, Dubinina 1974, Dubinina 1974, Thompson, Sue et al. 1982, Harinasuta and Harinasuta 1984, Ogbe 1985, Ogbe 1985, Mariano, Borja et al. 1986, Roberts, Lawson et al. 1986, Théron and Touassem 1989, Théron and Touassem 1989, Wu, Sun et al. 1991, Van Rensburg, Heitmann et al. 1997, Van Wyk, Van Rensburg et al. 1997, Zhang 1999, Henttonen, Fuglei et al. 2001, Madsen, Bloch et al. 2001, Dias, Eiras et al. 2003, Andreassen, Ito et al. 2004, Kumchoo, Wongsawad et al. 2005, Webster, Southgate et al. 2006, Dorny and Praet 2007, Li, Zhang et al. 2008, Möhl, Große et al. 2009, Collins III, King et al. 2011, De Liberato, Scaramozzino et al. 2011, Han and Xue 2011, Arai 2012, Świderski, Poddubnaya et al. 2012, Doanh, Horii et al. 2013, Del Brutto, García et al. 2014, Huang, Huang et al. 2014, Jiezhu, Xinyan et al. 2014, Malcicka 2015, Oksanen and Lavikainen 2015, Tang Chongti 2015, Thompson 2015, Devkota, Brant et al. 2016, Farrow 2016, Hong, Feng et al. 2016, Kvach, Ondracková et al. 2016, Léger, Garba et al. 2016, Moazeni and Ahmadi 2016, Arrabal, Avila et al. 2017, Blasco-Costa, Poulin et al. 2017, de la Torre-Escudero, Pérez-Sánchez et al. 2017, Lee, Park et al. 2017, Mohanta, Rana et al. 2017, Hildebrand, Pyrka et al. 2019, Prasad, Dahal et al. 2019, Sereno-Uribe, López-Jimenez et al. 2019, Eom, Rim et al. 2020, Kikuchi and Maruyama 2020, Kremnev, Gonchar et al. 2020, Radačovská, Bazsalovicsová et al. 2020, Wu, Gao et al. 2020, Sasaki, Kobayashi et al. 2021, Zajac, Zoller et al. 2021, Pyrka, Kanarek et al. 2022, Varcasia, Tamponi et al. 2022, de Buron, Hill-Spanik et al. 2023, Furusawa, Ikezawa et al. 2023).

#### Categorization and Filtering of Homologous Nuclear Genome Proteins Into Three Groups by PhyloFisher

The method refers to a recently published article: (Zhang, Jakovlić et al. 2024). The nuclear genome sequences provided by Brabec, Salomaki et al. (2023) were incorporated into the protein matrix containing the single-protein homologs from the initial matrix using the "working_dataset_constructor.py" script of PhyloFisher (Tice, Žihala et al. 2021). The resulting fasta files were subjected to filtering to remove non-homologous sites and sequencing errors using PREQUAL (Whelan, Irisarri et al. 2018), followed by the alignment using MAFFT's G-INS-I algorithm. To address the alignment uncertainty and errors, we applied DIVVIER 56 (Ali, Bogusz et al. 2019) with the options "-partial -mincol 4 -divvygap" to filter out problematic regions. The filtered alignments were further processed by trimAl (using the parameter "-gt 0.1") to remove sites that comprised more than 90% gaps (Capella-Gutiérrez, Silla-Martínez et al. 2009). Phylogenetic trees were constructed for each resulting alignment using IQ-TREE under the LG + C20 + F + G4 profile mixture model, with 5000 ultrafast bootstraps (selected as a trade-off between speed and reliability). The resulting trees were manually examined through the graphical user interface of PhyloFisher – ParaSorter to identify orthologues, paralogues, and potential contaminants for each taxon. Among the identified orthologues, those with taxon occupancy less than 60% were excluded, resulting in a final matrix containing 220 genes. To match the availability of mtDNA, PhyloFisher's "select_taxa.py" script was utilized to create the final nuclear DNA (nDNA) matrix consisting of 22 taxa (19 Trematoda + three outgroups) and 220 genes.

### Supplementary Results

#### Genome Organization and Base Composition

The length of the complete mitogenome of *Aspidogaster ijimai* was 13599 bp, and the length of *Aspidogaster conchicola* was 15532 bp (Fig. S40). Both mitogenomes contained the standard 36 neodermatan mitochondrial genes, including 12 protein-coding genes (PCGs, *atp8* not found), 22 tRNA genes and 2 rRNA genes (Table S1). The anti-codon of tRNA-Ile of *A. ijimai* mutated from GAT to GGT. The A+T content of the PCGs of *A. conchicola* (62.4%) was slightly higher than that of *A. ijimai* (58.7%), but their A+T skews were similar (Table S2). The A+T skew at the first codon site was lower than at the second and third codon sites.

Most of the start codons (ATG/GTG) of PCGs in *A. ijimai* and *A. conchicola* were standard for the genetic code 9 (echinoderm and ﬂatworm mitochondrion). A non-standard start codon, TTG, was identified in *nad4L* of *A. ijimai*, and *nad2* and *nad6* of *A. conchicola*. Recent studies indicate that TTG might be relatively common in flatworms (Ye, King et al. 2014, Zhang, Zou et al. 2018). Most stop codons of the PCGs of *A. ijimai* and *A. conchicola* were also standard TAG and TAA codons, but *cox3* and *nad4* of both aspidogastreans and *nad5* of *A. ijimai* used an abbreviated T-codon. Finally, the similarity of mitochondrial PCGs of these two aspidogastreans ranged from 66.15% (*nad5*) to 71.67% (*cox3*).

#### Gene Orders

The GOs of the two aspidogastreans were identical (similarity score = 1254). The similarity scores between the Schistosomatoidea species (except for *C. complanatum*) and the ancestral GO (inferred by Zhang, Li et al. (2019)) were lower than 520. However, the scores between the two aspidogastreans and the ancestral GO were both 1186 (Dataset S3), as they exhibited rearrangements of only *trnE* and *trnG* in comparison to the ancestral GO.

Table S1 Comparison of the architecture of the mitochondrial genomes of *Aspidogaster ijimai* and *Aspidogaster conchicola.*

| Gene | Position | | Size | Intergenic nucleotides | Codon | | Identity |
| --- | --- | --- | --- | --- | --- | --- | --- |
|  | From | To |  |  | Start | Stop |  |
| *Aspidogaster conchicola/Aspidogaster ijimai* | | | | | | | |
| *cox3* | 1/1 | 646/646 | 646/646 |  | ATG/ATG | T/T | 71.67 |
| *trnH* | 647/647 | 717/716 | 71/70 |  |  |  | 73.24 |
| *cytb* | 721/720 | 1836/1835 | 1116/1116 | 3/3 | ATG/ATG | TAG/TAG | 78.41 |
| *nad4L* | 1844/1836 | 2104/2096 | 261/261 | 7/- | ATG/TTG | TAG/TAG | 76.63 |
| *nad4* | 2065/2057 | 3340/3326 | 1276/1270 | -40/-40 | GTG/GTG | T/T | 71.16 |
| *trnQ* | 3342/3328 | 3402/3388 | 61/61 | 1/1 |  |  | 83.61 |
| *trnF* | 3403/3389 | 3465/3450 | 63/62 |  |  |  | 82.54 |
| *trnM* | 3466/3451 | 3525/3512 | 60/62 |  |  |  | 82.26 |
| *atp6* | 3532/3516 | 4044/4028 | 513/513 | 6/3 | GTG/ATG | TAA/TAG | 73.49 |
| *nad2* | 4047/4032 | 4904/4892 | 858/861 | 2/3 | TTG/GTG | TAA/TAG | 69.69 |
| *trnV* | 4913/4897 | 4975/4957 | 63/61 | 8/4 |  |  | 85.71 |
| *trnA* | 4976/4959 | 5040/5020 | 65/62 | -/1 |  |  | 70.77 |
| *trnD* | 5043/5022 | 5103/5084 | 61/63 | 2/1 |  |  | 76.19 |
| *nad1* | 5107/5072 | 6006/5986 | 900/915 | 3/-13 | ATG/GTG | TAG/TAG | 73.97 |
| *trnN* | 6007/5986 | 6071/6047 | 65/62 | -/-1 |  |  | 80 |
| *trnP* | 6081/6052 | 6145/6115 | 65/64 | 9/4 |  |  | 66.15 |
| *trnI* | 6153/6122 | 6218/6190 | 66/69 | 7/6 |  |  | 84.06 |
| *trnK* | 6233/6199 | 6293/6261 | 61/63 | 14/8 |  |  | 69.84 |
| *nad3* | 6294/6262 | 6647/6615 | 354/354 |  | ATG/ATG | TAG/TAG | 74.29 |
| *trnS1* | 6648/6617 | 6703/6673 | 56/57 | -/1 |  |  | 70.18 |
| *trnW* | 6707/6677 | 6775/6740 | 69/64 | 3/3 |  |  | 63.77 |
| *cox1* | 6779/6744 | 8323/8288 | 1545/1545 | 3/3 | GTG/ATG | TAG/TAG | 78.77 |
| *trnT* | 8335/8297 | 8397/8358 | 63/62 | 11/8 |  |  | 68.25 |
| *rrnL* | 8398/8359 | 9354/9314 | 957/956 |  |  |  | 80.81 |
| *trnC* | 9355/9315 | 9414/9376 | 60/62 |  |  |  | 70.97 |
| *rrnS* | 9415/9377 | 10135/10098 | 721/722 |  |  |  | 78.21 |
| *cox2* | 10136/10099 | 10705/10674 | 570/576 |  | ATG/ATG | TAG/TAG | 66.32 |
| *nad6* | 10706/10684 | 11155/11130 | 450/447 | -/9 | TTG/GTG | TAA/TAG | 66.67 |
| *trnY* | 11157/11132 | 11220/11195 | 64/64 | 1/1 |  |  | 75.38 |
| *trnL1* | 11229/11206 | 11292/11269 | 64/64 | 8/10 |  |  | 65.62 |
| *trnS2* | 11290/11267 | 11354/11331 | 65/65 | -3/-3 |  |  | 83.08 |
| *trnL2* | 11359/11336 | 11421/11400 | 63/65 | 4/4 |  |  | 73.85 |
| *trnR* | 11424/11401 | 11481/11462 | 58/62 | 2/- |  |  | 67.74 |
| *nad5* | 11460/11464 | 13058/13027 | 1599/1564 | -22/1 | ATG/GTG | TAA/T | 66.23 |
| *trnG* | 13284/13028 | 13348/13094 | 65/67 | 225/- |  |  | 82.09 |
| *trnE* | 13728/13246 | 13791/13311 | 64/66 | 379/151 |  |  | 71.21 |

Codon usage, RSCU, and the ratio of codon families (based on amino acid usage) analyses revealed that Val (12.31% and 11.05%), Phe (12.04% and 11.84%), and Leu2 (10.76% and 11.9%) codon families were the most commonly used amino acids in *A. ijimai* and *A. conchicola* respectively (Fig. S41).

Table S2 A comparison of nucleotide composition and skewness of different elements of mitochondrial genomes of *Aspidogaster ijimai* and *Aspidogaster conchicola.*

| Region | Size (bp) | T(U) | C | A | G | AT(%) | GC(%) | GT(%) | AT skew | GC skew |
| --- | --- | --- | --- | --- | --- | --- | --- | --- | --- | --- |
| *Aspidogaster conchicola/Aspidogaster ijimai* | | | | | | | | | | |
| PCGs | 10086/10065 | 46.2/43 | 11.5/13.6 | 16.2/15.7 | 26.1/27.7 | 62.4/58.7 | 37.6/41.3 | 72.3/70.7 | -0.48/-0.466 | 0.389/0.34 |
| 1st codon | 3362/3355 | 39.8/38.6 | 12.3/13.4 | 18.6/18 | 29.2/30.1 | 58.4/56.6 | 41.5/43.5 | 69/68.7 | -0.362/-0.364 | 0.408/0.385 |
| 2nd codon | 3362/3355 | 48.6/48.6 | 15.2/15.5 | 14.3/14.6 | 21.8/21.4 | 62.9/63.2 | 37/36.9 | 70.4/70 | -0.545/-0.538 | 0.179/0.16 |
| 3rd codon | 3362/3355 | 50.1/41.9 | 7/12.1 | 15.7/14.4 | 27.2/31.6 | 65.8/56.3 | 34.2/43.7 | 77.3/73.5 | -0.522/-0.488 | 0.592/0.446 |
| *atp6* | 513/513 | 51.1/46.2 | 11.7/12.7 | 11.5/14 | 25.7/27.1 | 62.6/60.2 | 37.4/39.8 | 76.8/73.3 | -0.632/-0.534 | 0.375/0.363 |
| *cox1* | 1545/1545 | 44.2/40.3 | 12.6/15.7 | 17.5/16.7 | 25.6/27.2 | 61.7/57 | 38.2/42.9 | 69.8/67.5 | -0.432/-0.414 | 0.34/0.268 |
| *cox2* | 570/576 | 36.3/34.9 | 12.5/15.1 | 22.3/19.6 | 28.9/30.4 | 58.6/54.5 | 41.4/45.5 | 65.2/65.3 | -0.24/-0.28 | 0.398/0.336 |
| *cox3* | 646/646 | 48.1/43.7 | 10.1/11.6 | 15.9/16.3 | 25.9/28.5 | 64/60 | 36/40.1 | 74/72.2 | -0.502/-0.457 | 0.44/0.421 |
| *cytb* | 1116/1116 | 44.8/41.9 | 12.5/13.7 | 18.5/18.6 | 24.1/25.7 | 63.3/60.5 | 36.6/39.4 | 68.9/67.6 | -0.414/-0.385 | 0.315/0.305 |
| *nad1* | 900/915 | 46.1/42.1 | 9.7/11.9 | 16.4/16.3 | 27.8/29.7 | 62.5/58.4 | 37.5/41.6 | 73.9/71.8 | -0.474/-0.442 | 0.484/0.428 |
| *nad2* | 858/861 | 48.3/47 | 12.4/13.2 | 14.8/15.1 | 24.6/24.6 | 63.1/62.1 | 37/37.8 | 72.9/71.6 | -0.53/-0.514 | 0.331/0.301 |
| *nad3* | 354/354 | 47.7/48.3 | 12.7/11.6 | 13.8/15.3 | 25.7/24.9 | 61.5/63.6 | 38.4/36.5 | 73.4/73.2 | -0.55/-0.52 | 0.338/0.364 |
| *nad4* | 1276/1270 | 47.1/43.8 | 12.9/15.5 | 14.3/13.1 | 25.6/27.6 | 61.4/56.9 | 38.5/43.1 | 72.7/71.4 | -0.533/-0.54 | 0.329/0.281 |
| *nad4L* | 261/261 | 51/47.1 | 6.9/7.7 | 16.5/17.2 | 25.7/28 | 67.5/64.3 | 32.6/35.7 | 76.7/75.1 | -0.511/-0.464 | 0.576/0.57 |
| *nad5* | 1599/1564 | 47/43.2 | 10.4/13.2 | 15.4/13.6 | 27.2/30.1 | 62.4/56.8 | 37.6/43.3 | 74.2/73.3 | -0.507/-0.522 | 0.448/0.388 |
| *nad6* | 450/447 | 47.6/46.3 | 8.9/13.9 | 16.7/14.5 | 26.9/25.3 | 64.3/60.8 | 35.8/39.2 | 74.5/71.6 | -0.481/-0.522 | 0.503/0.291 |
| *rrnL* | 957/956 | 37.8/36.2 | 12.7/13.4 | 23.5/23.7 | 25.9/26.7 | 61.3/59.9 | 38.6/40.1 | 63.7/62.9 | -0.233/-0.208 | 0.341/0.332 |
| *rrnS* | 721/722 | 35.4/32.1 | 15/15.8 | 23.4/24 | 26.2/28.1 | 58.8/56.1 | 41.2/43.9 | 61.6/60.2 | -0.203/-0.146 | 0.273/0.281 |
| rRNAs | 1678/1678 | 36.8/34.4 | 13.7/14.4 | 23.5/23.8 | 26/27.3 | 60.3/58.2 | 39.7/41.7 | 62.8/61.7 | -0.221/-0.182 | 0.31/0.309 |
| tRNAs | 1392/1397 | 34.8/31.1 | 14.7/17.5 | 22.6/23.1 | 27.9/28.3 | 57.4/54.2 | 42.6/45.8 | 62.7/59.4 | -0.213/-0.147 | 0.31/0.237 |
| Full genome | 15532/13599 | 42.8/40.5 | 11.8/14 | 19.5/18 | 25.8/27.5 | 62.3/58.5 | 37.6/41.5 | 68.6/68 | -0.373/-0.384 | 0.374/0.324 |

#### Transfer RNA Genes

Nineteen tRNAs of *A. ijimai* and twenty-one tRNAs of *A. conchicola* could be folded into the standard cloverleaf structure, with the exception of *tRNA^Cys(GCA)^*, *tRNA^Arg(TCG)^* and *tRNA^Ser(GCT)^* in the former, and the *tRNA^Ser(GCT)^* in the latter, which lacked DHU arms in both species. The concatenated size of tRNAs was nearly identical: 1397bp in *A. ijimai* and 1392bp in *A. conchicola*.

#### Non-coding Regions

There were two large non-coding regions (NCR, 100bp was selected as threshold for “large” NCR) in the mitogenome of *A. ijimai* (151 bp and 288 bp), and three (225 bp, 379 bp and 1741 bp) in *A. conchicola* (Fig. S40). Both mitogenomes exhibited the largest NCR between *trnE* and *cox3* (Fig. S40), and both of these comprised a highly repetitive region (HRR). The major NCR in *A. conchicola* was strongly enlarged, with 1741 bp, and it was the major underlying reason for the different mitogenome sizes of two aspidogastreans. The HRR of the 1741bp NCR comprised 14 tandem repeats (TRs). Among these, the sequences of repeat units 3, 4, 7, 8, 10 and 11 were identical, while others exhibited nucleotide mutations and deletions. The HRR of the 288 bp NCR of *A. ijimai* contained four TRs with a consensus size of 56 bp. Three of these exhibited identical sequences, whereas unit 4 exhibited a deletion of some downstream nucleotides. Both consensus repeat patterns of the HRRs in *A. conchicola* and *A. ijimai* were capable of forming stem-loop structures (Fig. S42). Corresponding findings were reported in other major parasitic flatworm radiations: “monogeneans” (Park, Kim et al. 2007, Zhang, Zou et al. 2018) and cestodes (von Nickisch-Rosenegk, Brown et al. 2001, Kyu-Heon, Hyeong-Kyu et al. 2007). It had been reported that the presence of tandem repeats forming stable secondary structures was associated with the replication origin in mitogenomes (Fumagalli, Taberlet et al. 1996, Park, Kim et al. 2007), so these repeat regions might be embedded within the control region.

#### Nucleotide Diversity and Evolutionary Rate Analysis

The nucleotide diversity in the concatenated alignment of 12 PCGs, 22 tRNAs and 2 rRNAs for the two aspidogastreans is shown in Fig. S43a. The Pi values based on the 200 bp sliding window ranged from 0.179 (*rrnL*) to 0.323 (*nad5*). High nucleotide diversity was exhibited by *cox2* (0.332), *nad5* (0.323), and *nad6* (0.308) genes, while *cox1* (0.213), *cytb* (0.213), *rrnL* (0.179), *rrnS* (0.210), and tRNAs (0.229) exhibited low diversity (low Pi values). Combined with the non-synonymous/synonymous (dN/dS) ratio analysis (Fig. S43b), the similarity analysis of genes (Table S1 shows that genes with higher dN/dS value (> 0.1) and lower identity (all < 72%) always had higher nucleotide diversity (all > 0.28), indicating a comparatively faster rate of evolution. Examples are *cox2*, *cox3* and *nad4*. These results were also consistent with previous research: *cox1*, which is commonly used as an universal barcode for species identification (Hebert, Cywinska et al. 2003), and population genetics in Trematoda (Wicht, Ruggeri-Bernardi et al. 2010, Králová-Hromadová, Čisovská Bazsalovicsová et al. 2011), was the slowest evolving and least variable gene (with the lowest dN/dS value).

#### Phylogenetic Analysis Based on Commonly-used Single-locus Molecular Markers

The commonly-used single-locus molecular markers often produce variable phylogenetic relationships among studies and datasets (Pérez-Ponce de León and Hernández-Mena 2019, Xu, Zhu et al. 2021). For example, using a partial *28S* rDNA, Heronimata and Hemiurata suborders were sister groups in the maximum-likelihood (ML) analysis, but not in the Bayesian Inference (BI) analysis (Pérez-Ponce de León and Hernández-Mena 2019). At the family level, *18S* and *28S* rDNA genes in combination with the (ML) method resolved Schistosomatidae as sister-clade to Diplostomidae and Brachylaimidae respectively; whereas an *ITS-1* dataset resolved Troglotrematidae and Notocotylidae as sister lineages in the ML-based topology, but not in the BI-based topology (Xu, Zhu et al. 2021).

#### Phylogenetic Analysis with Monopisthocotylea (“monogenea”) as the Outgroup

Regarding the superfamily-level phylogeny, homogeneous models resolved all superfamilies as monophyletic, apart from Schistosomatoidea, due to *Clinostomum complanatum* (Schistosomatoidea) forming a sister clade with *Postharmostomum commutatum* (Brachylaimoidea) (Figs. S20-25). As for the results of PhyloBayes, Schistosomatoidea was resolved as monophyletic in the NUCPBm topology, but in the AAPBm topology, *C. complanatum* formed a sister clade with the genus *Schistosoma*, which rendered Schistosomatoidea paraphyletic.

#### Phylogenetic Analysis with Polyopisthocotylea (”monogenea”) as the Outgroup

Regarding the order-level phylogeny, the topology reconstructed using the standard (“homogeneous”) model of IQ-TREE (NUCIQmp, Fig. S13), produced the two Aspidogastrea species nested within the Diplostomida clade. Accordingly, it resolved Diplostomida as paraphyletic and Plagiorchiida as monophyletic. However, the topology reconstructed with CAT-GTR in PhyloBayes (NUCPBmp, Fig. S14) resolved Trematoda as a polytomy. Regarding the superfamily-level phylogeny, NUCIQmp also resolved all superfamilies as monophyletic, apart from Schistosomatoidea. As for NUCPBmp, all superfamilies including Schistosomatoidea were resolved as monophyletic.

#### Phylogenetic Analysis with Cestoda as the Outgroup

Regarding the superfamily-level phylogeny, homogeneous models produced monophyletic superfamilies. The exception was Schistosomatoidea, which was mainly resolved as paraphyletic due to the unstable position of *C. complanatum*. Among the results of heterogeneous models, all superfamilies were monophyletic in the NUCPBc topology (Fig. 1d), but Schistosomatoidea was rendered paraphyletic by *P. commutatum* nested within it in AAPBc and AAIQc-He topologies (Fig. S32 and Fig. S10). Details of the interrelationships of the 14 superfamilies are displayed in Table 2 in main text.

#### Partitioning

Partitioning affected the topologies reconstructed using both AA and NUC datasets. With monogeneans as the outgroup, the topologies reconstructed using the partitioned AAm dataset and homogeneous models (AAIQPm and AAMBPm, Fig. S44-S45) produced paraphyletic Diplostomida due to Aspidogastrida nested within it. Conversely, the topologies reconstructed using the NUCm dataset (NUCIQPm and NUCMBPm) resolved Aspidogastrida as the sister group to digeneans, and all three digenean orders (Aspidogastrida, Diplostomida and Plagiorchiida) as monophyletic (Fig. S46-S47). With cestodes as the outgroup, the topologies reconstructed using both NUCc and AAc datasets (both partitioned; AAIQPc, AAMBPc, NUCIQPc and NUCMBPc, Fig. S48-S51) all resolved Aspidogastrida as the sister group to digeneans. Interestingly, the AAc dataset produced paraphyletic Diplostomida, but the NUCc dataset resolved it as monophyletic. Notably, the Azygioidea superfamily was placed at the root of the Plagiorchiida order in NUCIQPc and NUCMBPc topologies (the order was monophyletic), but it was resolved at the root of the subclass Digenea in NUCIQc and NUCMBc topologies, producing paraphyletic Plagiorchiida.

#### Basic Tree Comparison

Regarding the basic tree comparison results (Huerta-Cepas, Serra et al. 2016) (for details, see Dataset S2: “ete3” sheet), in all four testing groups (two different outgroups and two different datasets), the “%ref_br” (frequency of edges in the reference tree found in the target tree) and the “%src_br” (frequency of edges in the target tree found in the reference tree) values of topologies were all higher than 75%. Besides, the RF (Robinson-Foulds symmetric distance) values were all lower than 25. Interestingly, among the topologies reconstructed with monogeneans as the outgroup, the topologies which supported the basal position of Aspidogastrea, NUCPBm, NUCIQPm, AAIQm-He, AAIQPm, AASYMm and AASYM-Ho, had the highest %ref_br and %src_br values (both 0.85), and the lowest “RF” values (0.15).

#### Ancestral State Reconstruction

In order to further verify the impact of branch length on the ancestral state reconstruction results, we set all branch lengths of the NUCc-AL-cons topology to 1, and conducted multiple ancestral state reconstruction analyses. All analyses produced similar probabilities for all tested states: 1. when there were three possible ancestral states, 0, 1, and 2, each had a probability of 0.33; 2. when there were two possible ancestral states, 0 and 1, both had a probability of approximately 0.5 (for details, see Dataset S2 “ancestor”). As branch length affecting the ancestral state reconstruction results is a recognized phenomenon (Andrew Meade 2021), we can conclude that branch length probably played a crucial role in producing the contradictory result.

#### Evolutionary Scenarios for Trematoda Inferred Using the Nuclear Genome Data

In light of the evidence of a strong mitonuclear discordance in the topologies of Neodermata produced by the two genomic compartments, as well as the fact that nuclear genes associated with mitogenomic metabolism also produced a different topology than the remaining nuclear genes (Zhang, Jakovlić et al. 2024), we used the nuclear genomic dataset assembled by Brabec, Salomaki et al. (2023) and categorized the genes into three groups: nuclear genes not associated with mitochondria (non-MANE; AAn-nomito-T), mitochondria-associated nuclear-encoded genes (MANE; AAn-mito-T topology), and a subset of MANE comprising only the nuclear-encoded OXPHOS genes (OXPHOS; AAn-OXPHOS-T topology). The three gene groups were used to reconstruct phylogenies: the interrelationships of Trematoda were identical between the AAn-mito-T and AAn-OXPHOS-T datasets (consistent with AAn-T, except for *Atriophallophorus winterbourni* and *Paragonimus westermani* forming a sister clade in the AAn-T), but different from the AAn-nomito-T (Fig. S52).

### Supplementary Discussion

#### Partitioning vs. No Partitioning

Partitioning also influenced the topology. Regarding the phylogenetic trees reconstructed using the NUC dataset (NUCm and NUCc datasets) and homogeneous models, partitioning consistently resulted in the basal phylogenetic position of Aspidogastrida and monophyletic three orders (Aspidogastrida, Diplostomida and Plagiorchiida). The nonpartitioned dataset produced Aspidogastrida nested within the (paraphyletic) Diplostomida (the NUCm dataset), and a paraphyletic Plagiorchiida order (NUCm and NUCc datasets). This indicates that partitioning appears to produce more reliable results. However, regarding the phylogenetic trees reconstructed with the AA dataset and homogeneous models, partitioned and nonpartitioned datasets all produced Aspidogastrida nested within the Diplostomida (based on the AAm dataset) and paraphyletic Diplostomida order (based on the AAm and AAc datasets). Interestingly, the use of heterogeneous models in combination with the AA dataset resolved these problems, which is in agreement with previous reports that heterogeneous models performed better than homogeneous models on highly heterogeneous datasets regardless of partitioning (Uribe, Irisarri et al. 2019, Wang, Susko et al. 2019). In short, partitioning improved the performance of the NUC datasets, but not the performance of the AA datasets.

#### Phylogenetic Positions of Certain Trematoda Lineages Require Further Investigation

When reconstructing the topologies with different outgroups, datasets and models, the phylogenetic positions of several families (Brachylaimidae and Clinostomidae) and superfamilies (Allocreadioidea, Troglotrematoidea, Opisthorchioidea, Gorgoderoidea, Microphalloidea, and Plagiorchioidea) were highly variable. Some topologies supported Brachylaimidae and Clinostomidae as a sister group to the Diplostomoidea superfamily, but others did not (for details, see Table 2 in main text). Several previous studies also observed that the positions of Brachylaimidae and Clinostomidae were unstable (Olson, Cribb et al. 2003, Pérez-Ponce de León and Hernández-Mena 2019, Chan, Saralamba et al. 2022). Regarding the superfamilies, Allocreadioidea was resolved as the sister clade to Opisthorchioidea in some topologies, and to Troglotrematoidea in others. Gorgoderoidea and Microphalloidea formed a sister clade to Plagiorchioidea in some topologies, but in some other topologies, Gorgoderoidea formed a sister clade with Allocreadioidea, Troglotrematoidea, and Opisthorchioidea superfamilies, while Microphalloidea was sister clade to Plagiorchioidea. Interestingly, the CAT-GTR model also failed to resolve these contradictory phylogenetic relationships; for example, Allocreadioidea and Troglotrematoidea were sister groups in the NUCPBm topology, but Allocreadioidea formed a sister clade with Opisthorchioidea in the AAPBm topology; and Plagiorchiida was resolved as paraphyletic in the NUCPBmp topology.

In short, although the CAT-GTR model managed to resolve some of the LBA artefacts, it failed to resolve all of the topological instability. These results indicate that additional datasets (such as genomic and transcriptomic data) and appropriate evolutionary models for phylogenetic reconstruction are required to resolve the phylogeny of Trematoda.

### References

Alba, A. (2018). Comparative biology of susceptible and naturally- resistant Pseudosuccinea columella snails to Fasciola hepatica (Trematoda) infection in Cuba : ecological, molecular and phenotypical aspects.

Ali, R. H., M. Bogusz and S. Whelan (2019). "Identifying clusters of high confidence homologies in multiple sequence alignments." Molecular biology evolution **36**(10): 2340-2351.

Altschul, S. F., W. Gish, W. Miller, E. W. Myers and D. J. Lipman (1990). "Basic local alignment search tool." Journal of Molecular Biology **215**(3): 403-410.

Andreassen, J., A. Ito, M. Ito, M. Nakao and K. Nakaya (2004). "Hymenolepis microstoma: direct life cycle in immunodeficient mice." J Journal of helminthology **78**(1): 1-5.

Andrew Meade, M. P. (2021). "BayesTraitsV3.0.5-Manual."

Arai, H. (2012). Biology of the Tapeworm Hymenolepis diminuta, Elsevier.

Arrabal, J. P., H. G. Avila, M. R. Rivero, F. Camicia, M. M. Salas, S. A. Costa, C. G. Nocera, M. C. Rosenzvit and L. Kamenetzky (2017). "Echinococcus oligarthrus in the subtropical region of Argentina: first integration of morphological and molecular analyses determines two distinct populations." J Veterinary Parasitology **240**: 60-67.

Bakker, K. and C. J. J. o. H. Davids (1973). "Notes on the life history of Aspidogaster conchicola Baer, 1826 (Trematoda; Aspidogastridae)." **47**(3): 269-276.

Benson, G. (1999). "Tandem repeats finder: a program to analyze DNA sequences." Nucleic Acids Research **27**(2): 573-580.

Bernt, M., A. Donath, F. Jühling, F. Externbrink, C. Florentz, G. Fritzsch, J. Pütz, M. Middendorf and P. F. Stadler (2013). "MITOS: Improved de novo metazoan mitochondrial genome annotation." Molecular Phylogenetics and Evolution **69**(2): 313-319.

Blasco-Costa, I., R. Poulin and B. Presswell (2017). "Morphological description and molecular analyses of Tylodelphys sp.(Trematoda: Diplostomidae) newly recorded from the freshwater fish Gobiomorphus cotidianus (common bully) in New Zealand." Journal of Helminthology **91**(3): 332-345.

Brabec, J., E. D. Salomaki, M. Kolísko, T. Scholz and R. Kuchta (2023). "The evolution of endoparasitism and complex life cycles in parasitic platyhelminths." Curr Biol.

Capella-Gutiérrez, S., J. M. Silla-Martínez and T. Gabaldón (2009). "trimAl: a tool for automated alignment trimming in large-scale phylogenetic analyses." Bioinformatics **25**(15): 1972-1973.

Castresana, J. (2000). "Selection of conserved blocks from multiple alignments for their use in phylogenetic analysis." Molecular biology evolution **17**(4): 540-552.

Chan, A. H. E., N. Saralamba, S. Saralamba, J. Ruangsittichai and U. Thaenkham (2022). "The potential use of mitochondrial ribosomal genes (12S and 16S) in DNA barcoding and phylogenetic analysis of trematodes." BMC genomics **23**(1): 104.

Clarke, A. (1954). Studies on the life cycle of the pseudophyllidean cestode Schistocephalus solidus. Proceedings of the Zoological Society of London, Wiley Online Library.

Cock, P. J., T. Antao, J. T. Chang, B. A. Chapman, C. J. Cox, A. Dalke, I. Friedberg, T. Hamelryck, F. Kauff, B. Wilczynski and M. J. de Hoon (2009). "Biopython: freely available Python tools for computational molecular biology and bioinformatics." Bioinformatics **25**(11): 1422-1423.

Collins III, J. J., R. S. King, A. Cogswell, D. L. Williams and P. A. Newmark (2011). "An atlas for Schistosoma mansoni organs and life-cycle stages using cell type-specific markers and confocal microscopy." J PLoS neglected tropical diseases **5**(3): e1009.

de Buron, I., K. M. Hill-Spanik, T. Baker, G. Fignar and J. Broach (2023). "Infection of Atlantic tripletail Lobotes surinamensis (Teleostei: Lobotidae) by brain metacercariae Cardiocephaloides medioconiger (Digenea: Strigeidae)." PeerJ **11**: e15365.

de la Torre-Escudero, E., R. Pérez-Sánchez, R. Manzano-Román and A. Oleaga (2017). "Schistosoma bovis-host interplay: Proteomics for knowing and acting." Molecular Biochemical Parasitology **215**: 30-39.

De Liberato, C., P. Scaramozzino, A. Brozzi, R. Lorenzetti, D. Di Cave, E. Martini, C. Lucangeli, E. Pozio, F. Berrilli and T. Bossù (2011). "Investigation on Opisthorchis felineus occurrence and life cycle in Italy." Veterinary parasitology **177**(1-2): 67-71.

Del Brutto, O. H., H. H. García, O. H. Del Brutto and H. H. García (2014). "Taenia solium: biological characteristics and life cycle." Cysticercosis of the Human Nervous System: 11-21.

Devkota, R., S. V. Brant and E. S. Loker (2016). "A genetically distinct Schistosoma from Radix luteola from Nepal related to Schistosoma turkestanicum: A phylogenetic study of schistosome and snail host." Acta tropica **164**: 45-53.

Dias, M., J. Eiras, M. Machado, G. Souza and G. Pavanelli (2003). "The life cycle of Clinostomum complanatum Rudolphi, 1814 (Digenea, Clinostomidae) on the floodplain of the high Paraná river, Brazil." Parasitology Research **89**: 506-508.

Doanh, P. N., Y. Horii and Y. Nawa (2013). "Paragonimus and paragonimiasis in Vietnam: an update." The Korean Journal of Parasitology **51**(6): 621.

Dorny, P. and N. Praet (2007). "Taenia saginata in Europe." J Veterinary Parasitology **149**(1-2): 22-24.

Dubinina, M. (1974). "The development of Amphilina foliacea (Rud.) at all stages of its life-cycle and the position of Amphilinidea in the Platyhelminthes." J Parazitologicheskii Sbornik, Leningrad(26): 9-38.

Dubinina, M. N. (1974). The development of Amphilina foliacea (Rud.) at all stages of its life-cycle and the position of Amphilinidea in the Platyhelminthes.

Eom, K. S., H.-J. Rim and H.-K. Jeon (2020). "Taenia asiatica: historical overview of taeniasis and cysticercosis with molecular characterization." J Advances in parasitology **108**: 133-173.

Farrow, A. (2016). Aspects of the Life Cycle of Apharyngostrigea Pipientis in Central Florida Wetlands, Florida Southern College.

Fumagalli, L., P. Taberlet, L. Favre and J. Hausser (1996). "Origin and evolution of homologous repeated sequences in the mitochondrial DNA control region of shrews." Molecular biology evolution **13 1**: 31-46.

Furusawa, H., H. Ikezawa, S. G. Tsujimoto, M. Ichikawa-Seki and T. Waki (2023). "Introducing the land snail Bradybaena pellucida increased infection risk of the avian parasite Postharmostomum commutatum in the Kanto region of Japan." Parasitology Research: 1-10.

Han, H. and D. Xue (2011). Zoology of China, Ke xue chu ban she.

Harinasuta, C. and T. Harinasuta (1984). "Opisthorchis viverrini: life cycle, intermediate hosts, transmission to man and geographical distribution in Thailand." J Arzneimittel-forschung **34**(9B): 1164-1167.

Hebert, P. D., A. Cywinska, S. L. Ball and J. R. deWaard (2003). "Biological identifications through DNA barcodes." Proc Biol Sci **270**(1512): 313-321.

Henttonen, H., E. Fuglei, C. Gower, V. Haukisalmi, R. A. Ims, J. Niemimaa and N. G. Yoccoz (2001). "Echinococcus multilocularis on Svalbard: introduction of an intermediate host has enabled the local life-cycle." J Parasitology **123**(6): 547-552.

Hildebrand, J., E. Pyrka, J. Sitko, W. Jeżewski, G. Zaleśny, V. V. Tkach and Z. Laskowski (2019). "Molecular phylogeny provides new insights on the taxonomy and composition of Lyperosomum Looss, 1899 (Digenea, Dicrocoeliidae) and related genera." International Journal for Parasitology: Parasites Wildlife **9**: 90-99.

Hong, Q., J. Feng, H. Liu, X. Li, L. Gong, Z. Yang, W. Yang, X. Liang, R. Zheng and Z. Cui (2016). "Prevalence of Spirometra mansoni in dogs, cats, and frogs and its medical relevance in Guangzhou, China." J International Journal of Infectious Diseases **53**: 41-45.

Huang, J., X. Huang, z. Liu and L. Li (2014). "Prevention and Control of Duck Schistosomiasis." Jiangxi Animal Husbandry and Veterinary Journal(6): 51-52.

Huerta-Cepas, J., F. Serra and P. Bork (2016). "ETE 3: Reconstruction, Analysis, and Visualization of Phylogenomic Data." Molecular Biology and Evolution **33**(6): 1635-1638.

Hutchison, W. (1959). "Studies on Hydatigera (Taenia) taeniaeformis. II. Growth of the adult phase." J Experimental Parasitology **8**(6): 557-567.

Jiezhu, H., H. Xinyan, L. Zhou and L. Lanming (2014). "Prevention and Control of Duck Naviculariasis." Jiangxi Animal Husbandry and Veterinary Journal(6): 51-52.

Kalyaanamoorthy, S., B. Q. Minh, T. K. F. Wong, A. von Haeseler and L. S. Jermiin (2017). "ModelFinder: fast model selection for accurate phylogenetic estimates." Nature Methods **14**(6): 587-589.

Katoh, K. and D. M. Standley (2013). "MAFFT multiple sequence alignment software version 7: improvements in performance and usability." Mol Biol Evol **30**(4): 772-780.

Katoh, K., D. M. J. M. b. Standley and evolution (2013). "MAFFT multiple sequence alignment software version 7: improvements in performance and usability." Molecular biology evolutionary **30**(4): 772-780.

Kaw, M. U. "THE LIFE CYCLE OF OR THOCOELIUM SCOLJOCOELIUM (FISCHOEDER, 1904) YAMAGUTI, 1971 (PARAMPHISTOMIDAE: ORTHOCOELIINAE) IN TH. E PIDLIPPINES."

Kikuchi, T. and H. Maruyama (2020). "Human proliferative sparganosis update." J Parasitology international **75**: 102036.

Králová-Hromadová, I., E. Čisovská Bazsalovicsová, J. Štefka, M. Špakulová, S. Vavrova, T. Szemes, V. Tkach, A. Trudgett and M. Pybus (2011). "Multiple origins of European populations of the giant liver fluke Fascioloides magna (Trematoda: Fasciolidae), a liver parasite of ruminants." International journal for parasitology **41**: 373-383.

Kremnev, G., A. Gonchar, V. Krapivin, O. Knyazeva and D. Krupenko (2020). "First elucidation of the life cycle in the family Brachycladiidae (Digenea), parasites of marine mammals." International Journal for Parasitology **50**(12): 997-1009.

Kumchoo, K., C. Wongsawad, J.-Y. Chai, P. Vanittanakom and A. Rojanapaibul (2005). "High prevalence of Haplorchis taichui metacercariae in cyprinoid fish from Chiang Mai Province, Thailand." Southeast Asian J Trop Med Public Health **36**(2): 451-455.

Kvach, Y., M. Ondracková and P. Jurajda (2016). "First report of metacercariae of Cyathocotyle prussica parasitising a fish host in the Czech Republic, Central Europe." Helminthologia **53**(3): 257.

Kyu-Heon, K., J. Hyeong-Kyu, K. Seokha, S. Tahera, K. Gil Jung, S. E. Keeseon and P. Joong-Ki (2007). "Characterization of the Complete Mitochondrial Genome of Diphyllobothrium nihonkaiense (Diphyllobothriidae: Cestoda), and Development of Molecular Markers for Differentiating Fish Tapeworms." Molecules and Cells **23**(3): 379-390.

Lartillot, N., N. Rodrigue, D. Stubbs and J. Richer (2013). "PhyloBayes MPI: Phylogenetic Reconstruction with Infinite Mixtures of Profiles in a Parallel Environment." Systematic Biology **62**(4): 611-615.

Laslett, D. and B. Canbäck (2008). "ARWEN: a program to detect tRNA genes in metazoan mitochondrial nucleotide sequences." Bioinformatics methods protocols **24**(2): 172-175.

Lee, D., H. Park, S. Choe, Y. Kang, H.-K. Jeon and K. S. J. T. K. J. o. P. Eom (2017). "New Record of Aspidogaster ijimai Kawamura, 1913 (Trematoda: Aspidogastridae) from Cyprinus carpio in Korea." **55**(5): 575.

Léger, E., A. Garba, A. A. Hamidou, B. L. Webster, T. Pennance, D. Rollinson and J. P. Webster (2016). "Introgressed animal schistosomes Schistosoma curassoni and S. bovis naturally infecting humans." Emerging infectious diseases **22**(12): 2212.

Leigh, W. H. (1946). "Experimental studies on the life cycle of Glypthelmins quieta (Stafford, 1900), a trematode of frogs." The American Midland Naturalist **35**(2): 460-483.

Letunic, I. and P. Bork (2024). "Interactive Tree of Life (iTOL) v6: recent updates to the phylogenetic tree display and annotation tool." Nucleic Acids Research: gkae268.

Li, W., L. Zhang, Q. Gao and P. Nie (2008). "Species composition and community characteristics of parasitic helminths in fishes in the Lhasa River, Tibet." Journal of Zoology **43**(2): 1-8.

Li, Y., X. X. Ma, Q. B. Lv, Y. Hu, H. Y. Qiu, Q. C. Chang and C. R. Wang (2019). "Characterization of the complete mitochondrial genome sequence of Tracheophilus cymbius (Digenea), the first representative from the family Cyclocoelidae." J Helminthol **94**: e101.

Librado, P. and J. Rozas (2009). "DnaSP v5: a software for comprehensive analysis of DNA polymorphism data." Bioinformatics **25**(11): 1451-1452.

Macy, R. W. (1965). "On the life cycle of the trematode Prosthogonimus cuneatus (Rudolphi, 1809)(Plagiorchiidae) in Egypt." Transactions of the American Microscopical Society **84**(4): 577-580.

Madsen, H., P. Bloch, H. Phiri, T. Kristensen and P. Furu (2001). "Bulinus nyassanus is an intermediate host for Schistosoma haematobium in Lake Malawi." Annals of Tropical Medicine Parasitology **95**(4): 353-360.

Malcicka, M. (2015). "Life history and biology of Fascioloides magna (Trematoda) and its native and exotic hosts." J Ecology evolution **5**(7): 1381-1397.

Maldonado, J. F. (1945). "The life cycle of Tamerlania bragai, Santos 1934,(Eucotylidae), a kidney fluke of domestic pigeons." The Journal of Parasitology **31**(5): 306-314.

Mariano, E. G., S. R. Borja and M. Vruno (1986). "A human infection with Paragonimus kellicotti (lung fluke) in the United States." American journal of clinical pathology **86**(5): 685-687.

Meyer, F. (1960). "Life history of Marsipometra hastata and the biology of its host." J Polyodon spathula.

Moazeni, M. and A. Ahmadi (2016). "Controversial aspects of the life cycle of Fasciola hepatica." J Experimental parasitology **169**: 81-89.

Mohanta, U., H. Rana, B. Devkota and T. Itagaki (2017). "Molecular and phylogenetic analyses of the liver amphistome Explanatum explanatum (Creplin, 1847) Fukui, 1929 in ruminants from Bangladesh and Nepal based on nuclear ribosomal ITS2 and mitochondrial nad1 sequences." Journal of helminthology **91**(4): 497-503.

Möhl, K., K. Große, A. Hamedy, T. Wüste, P. Kabelitz and E. Lücker (2009). "Biology of Alaria spp. and human exposition risk to Alaria mesocercariae—a review." Parasitology research **105**: 1-15.

Morand, S., Krasnov, B., & Littlewood, D. (2015). Parasite Diversity and Diversification: Evolutionary Ecology Meets Phylogenetics. Cambridge, Cambridge University Press.

Naser-Khdour, S., B. Q. Minh, W. Zhang, E. A. Stone and R. Lanfear (2019). "The prevalence and impact of model violations in phylogenetic analysis." Genome biology evolution **11**(12): 3341-3352.

Nguyen, B. T., K. Shuval and A. L. Yaroch (2015). "Nguyen et al. respond." American journal of public health **105**(10): e2.

Ogbe, M. G. (1985). "Aspects of the life cycle of Schistosoma margrebowiei infection in laboratory mammals." Int J Parasitol **15**(2): 141-145.

Ogbe, M. G. (1985). "Aspects of the life cycle of Schistosoma margrebowiei infection in laboratory mammals." J International journal for parasitology **15**(2): 141-145.

Oksanen, A. and A. Lavikainen (2015). "Echinococcus canadensis transmission in the North." J Veterinary Parasitology **213**(3-4): 182-186.

Olson, P. D., T. H. Cribb, V. V. Tkach, R. A. Bray and D. T. J. Littlewood (2003). "Phylogeny and classification of the Digenea (Platyhelminthes: Trematoda)11Nucleotide sequence data reported in this paper are available in the GenBank™, EMBL and DDBJ databases under the accession numbers AY222082–AY222285." International Journal for Parasitology **33**(7): 733-755.

Park, J.-K., K.-H. Kim, S. Kang, W. Kim, K. S. Eom and D. Littlewood (2007). "A common origin of complex life cycles in parasitic flatworms: evidence from the complete mitochondrial genome of Microcotyle sebastis (Monogenea: Platyhelminthes)." BMC Evolutionary Biology **7**: 1-13.

Pérez-Ponce de León, G. and D. I. Hernández-Mena (2019). "Testing the higher-level phylogenetic classification of Digenea (Platyhelminthes, Trematoda) based on nuclear rDNA sequences before entering the age of the 'next-generation' Tree of Life." J Helminthol **93**(3): 260-276.

Prasad, Y. K., S. Dahal, B. Saikia, B. Bordoloi, V. Tandon and S. Ghatani (2019). "Artyfechinostomum sufrartyfex trematode infections in children, Bihar, India." Emerging Infectious Diseases **25**(8): 1571.

Pyrka, E., G. Kanarek, J. Gabrysiak, W. Jeżewski, A. Cichy, A. Stanicka, E. Żbikowska, G. Zaleśny and J. Hildebrand (2022). "Life history strategies of Cotylurus spp. Szidat, 1928 (Trematoda, Strigeidae) in the molecular era–Evolutionary consequences and implications for taxonomy." International Journal for Parasitology: Parasites Wildlife **18**: 201-211.

Radačovská, A., E. Bazsalovicsová, I. B. Costa, M. Orosová, A. Gustinelli and I. Králová-Hromadová (2020). "Occurrence of Dibothriocephalus latus in European perch from Alpine lakes, an important focus of diphyllobothriosis in Europe." J Revue suisse de Zoologie **126**(2): 219-225.

Roberts, M., J. Lawson and M. Gemmell (1986). "Population dynamics in echinococcosis and cysticercosis: mathematical model of the life-cycle of Echinococcus granulosus." J Parasitology **92**(3): 621-641.

Ronquist, F., M. Teslenko, P. Van Der Mark, D. L. Ayres, A. Darling, S. Höhna, B. Larget, L. Liu, M. A. Suchard and J. P. Huelsenbeck (2012). "MrBayes 3.2: efficient Bayesian phylogenetic inference and model choice across a large model space." Systematic biology **61**(3): 539-542.

Sasaki, M., M. Kobayashi, T. Yoshino, M. Asakawa and M. Nakao (2021). "Notocotylus ikutai n. sp.(Digenea: Notocotylidae) from lymnaeid snails and anatid birds in Hokkaido, Japan." Parasitology International **83**: 102318.

Seo, B.-S. and J.-W. Kwak (1972). "Studies on the lung fluke, Paragonimus iloktsuenensis. II. On the metacercaria, the second intermediate hosts and development in mice." Seoul Journal of Medicine **13**(4).

Sereno-Uribe, A., A. López-Jimenez, L. Andrade-Gómez and M. García-Varela (2019). "A morphological and molecular study of adults and metacercariae of Hysteromorpha triloba (Rudolpi, 1819), Lutz 1931 (Diplostomidae) from the Neotropical region." Journal of Helminthology **93**(1): 91-99.

Sindermann, C. J. and A. E. Farrin (1962). "Ecological studies of Cryptocotyle lingua (Trematoda: Heterophyidae) whose larvae cause" pigment spots" of marine fish." Ecology: 69-75.

Srivastava, H. (1944). "A study of the life-history of Gastrothylax crumenifer of Indian ruminants." (Part III).

Stamatakis, A. (2014). "RAxML version 8: a tool for phylogenetic analysis and post-analysis of large phylogenies." Bioinformatics **30**(9): 1312-1313.

Świderski, Z., L. G. Poddubnaya, D. I. Gibson and D. Młocicki (2012). "Advanced stages of embryonic development and cotylocidial morphogenesis in the intrauterine eggs of Aspidogaster limacoides Diesing, 1835 (Aspidogastrea), with comments on their phylogenetic implications." J Acta Parasitologica **57**: 131-148.

Tang Chongti, T. Z. (2015). Chinese Trematology, Science Press.

Théron, A. and R. Touassem (1989). "Schistosoma rodhaini: intramolluscan larval development, migration and replication processes of daughter sporocysts." J Acta tropica **46**(1): 39-45.

Théron, A. and R. Touassem (1989). "Schistosoma rodhaini: intramolluscan larval development, migration and replication processes of daughter sporocysts." Acta Trop **46**(1): 39-45.

Thompson, R. (2015). "Neglected zoonotic helminths: Hymenolepis nana, Echinococcus canadensis and Ancylostoma ceylanicum." J Clinical Microbiology Infection **21**(5): 426-432.

Thompson, R., L. J. Sue and S. Buckley (1982). "In vitro development of the strobilar stage of Mesocestoides corti." J International Journal for Parasitology **12**(4): 303-314.

Tice, A. K., D. Žihala, T. Pánek, R. E. Jones, E. D. Salomaki, S. Nenarokov, F. Burki, M. Eliáš, L. Eme, A. J. Roger, A. Rokas, X. X. Shen, J. F. H. Strassert, M. Kolísko and M. W. Brown (2021). "PhyloFisher: A phylogenomic package for resolving eukaryotic relationships." PLoS Biol **19**(8): e3001365.

Uribe, J. E., I. Irisarri, J. Templado and R. Zardoya (2019). "New patellogastropod mitogenomes help counteracting long-branch attraction in the deep phylogeny of gastropod mollusks." Molecular phylogenetics evolution **133**: 12-23.

Van Rensburg, L., L. Heitmann and J. A. Van Wyk (1997). "Schistosoma mattheei infection in cattle: The course of the intestinal syndrome, and an estimate of the lethal dose of cercariae."

Van Wyk, J. A., L. J. Van Rensburg and L. P. Heitmann (1997). "Schistosoma mattheei infection in cattle: the course of the intestinal syndrome, and an estimate of the lethal dose of cercariae." Onderstepoort J Vet Res **64**(1): 65-75.

Varcasia, A., C. Tamponi, F. Ahmed, M. G. Cappai, F. Porcu, N. Mehmood, G. Dessì and A. Scala (2022). "Taenia multiceps coenurosis: A review." J Parasites Vectors **15**(1): 1-18.

von Nickisch-Rosenegk, M., W. M. Brown and J. L. Boore (2001). "Complete sequence of the mitochondrial genome of the tapeworm Hymenolepis diminuta: gene arrangements indicate that Platyhelminths are Eutrochozoans." Mol Biol Evol **18**(5): 721-730.

Wang, H.-C., E. Susko and A. J. Roger (2019). "The Relative Importance of Modeling Site Pattern Heterogeneity Versus Partition-Wise Heterotachy in Phylogenomic Inference." Systematic Biology **68**(6): 1003-1019.

Webster, B. L., V. R. Southgate and D. T. Littlewood (2006). "A revision of the interrelationships of Schistosoma including the recently described Schistosoma guineensis." International Journal for Parasitology: Parasites Wildlife **36**(8): 947-955.

Whelan, S., I. Irisarri and F. Burki (2018). "PREQUAL: detecting non-homologous characters in sets of unaligned homologous sequences." Bioinformatics **34**(22): 3929-3930.

Wicht, B., N. Ruggeri-Bernardi, T. Yanagida, M. Nakao, R. Peduzzi and A. J. P. i. Ito (2010). "Inter-and intra-specific characterization of tapeworms of the genus Diphyllobothrium (Cestoda: Diphyllobothriidea) from Switzerland, using nuclear and mitochondrial DNA targets." **59**(1): 35-39.

Wu, B., X. Sun and C. Song (1991). Zhejiang Fauna (Trematodes), 杭.

Wu, Y.-A., J.-W. Gao, X.-F. Cheng, M. Xie, X.-P. Yuan, D. Liu and R. Song (2020). "Characterization and comparative analysis of the complete mitochondrial genome of Azygia hwangtsiyui Tsin, 1933 (Digenea), the first for a member of the family Azygiidae." ZooKeys **945**: 1.

Xiang, C.-Y., F. Gao, I. Jakovlić, H.-P. Lei, Y. Hu, H. Zhang, H. Zou, G.-T. Wang and D. Zhang (2023). "Using PhyloSuite for molecular phylogeny and tree-based analyses." **2**(1): e87.

Xu, G., P. Zhu, W. Zhu, B. Ma, X. Li and W. Li (2021). "Characterization of the complete mitochondrial genome of Notocotylus sp. (Trematoda, Notocotylidae) and its phylogenetic implications." Parasitol Res **120**(4): 1291-1301.

Ye, F., S. D. King, D. K. Cone and P. You (2014). "The mitochondrial genome of Paragyrodactylus variegatus (Platyhelminthes: Monogenea): differences in major non-coding region and gene order compared to Gyrodactylus." Parasit Vectors **7**: 377.

Yokogawa, M. (1965). "Paragonimus and paragonimiasis." Advances in parasitology **3**: 99-158.

Zajac, N., S. Zoller, K. Seppälä, D. Moi, C. Dessimoz, J. Jokela, H. Hartikainen and N. Glover (2021). "Gene duplication and gain in the trematode Atriophallophorus winterbourni contributes to adaptation to parasitism." J Genome biology evolution **13**(3): evab010.

Zhang, D., F. Gao, I. Jakovlić, H. Zou, J. Zhang, W. X. Li and G. T. Wang (2020). "PhyloSuite: An integrated and scalable desktop platform for streamlined molecular sequence data management and evolutionary phylogenetics studies." **20**(1): 348-355.

Zhang, D., I. Jakovlić, H. Zou, F. Liu, C. Y. Xiang, Q. Gusang, S. Tso, S. Xue, W. J. Zhu, Z. Li, J. Wu and G. T. Wang (2024). "Strong mitonuclear discordance in the phylogeny of Neodermata and evolutionary rates of Polyopisthocotylea." Int J Parasitol.

Zhang, D., W. X. Li, H. Zou, S. G. Wu, M. Li, I. Jakovlić, J. Zhang, R. Chen and G. Wang (2019). "Homoplasy or plesiomorphy? Reconstruction of the evolutionary history of mitochondrial gene order rearrangements in the subphylum Neodermata." International Journal for Parasitology **49**(10): 819-829.

Zhang, D., H. Zou, S. Wu, M. Li, I. Jakovlić, J. Zhang, R. Chen, G. Wang and W. Li (2018). "Sequencing, characterization and phylogenomics of the complete mitochondrial genome of Dactylogyrus lamellatus (Monogenea: Dactylogyridae)." Journal of helminthology **92**(4): 455-466.

Zhang, J. (1999). Fish Parasites and Parasitic Diseases, Science Press.

Zhang, Z., J. Li, X. Q. Zhao, J. Wang, G. K. Wong and J. Yu (2006). "KaKs_Calculator: calculating Ka and Ks through model selection and model averaging." Genomics Proteomics Bioinformatics **4**(4): 259-263.

Zuker, M. (2003). "Mfold web server for nucleic acid folding and hybridization prediction." Nucleic Acids Res **31**(13): 3406-3415.
