## Supplementary figures and images for "The Phylogeny and the Evolution of Parasitic Strategies in Trematoda"

### S1-NUCSYMm.pdf

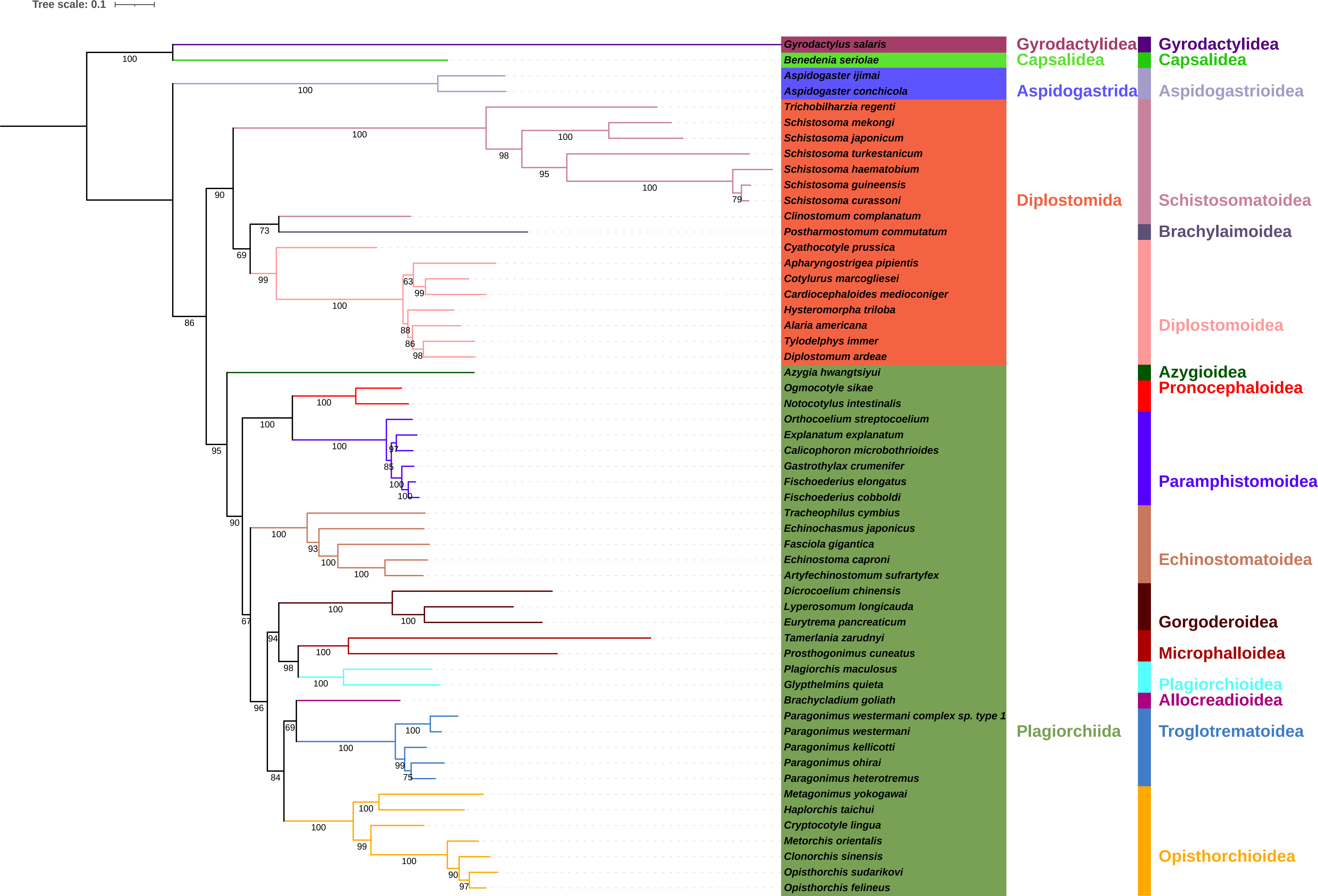

### S2-AASYMm.pdf

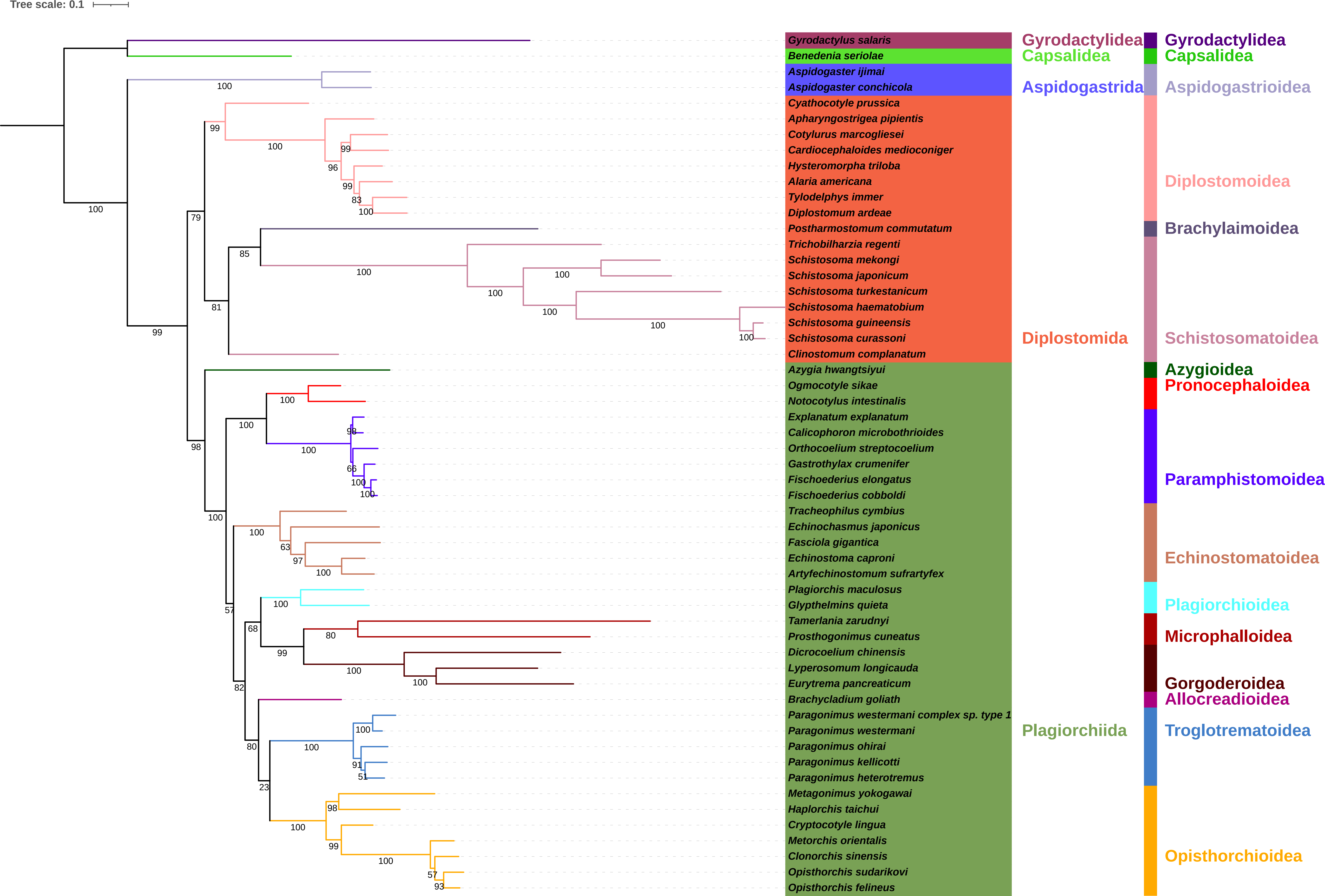

### S3-NUCSYMc.pdf

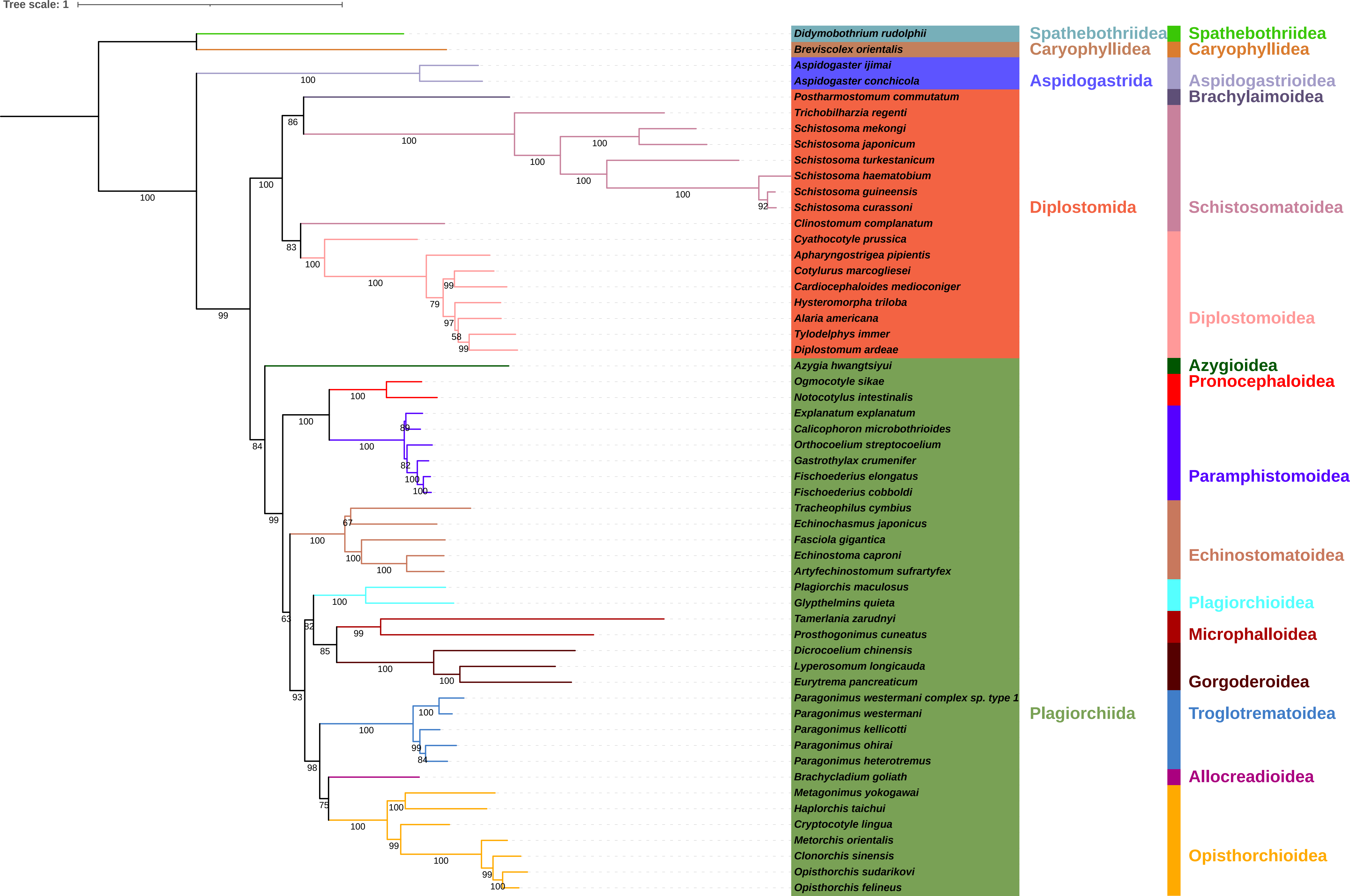

### S4-AASYMc.pdf

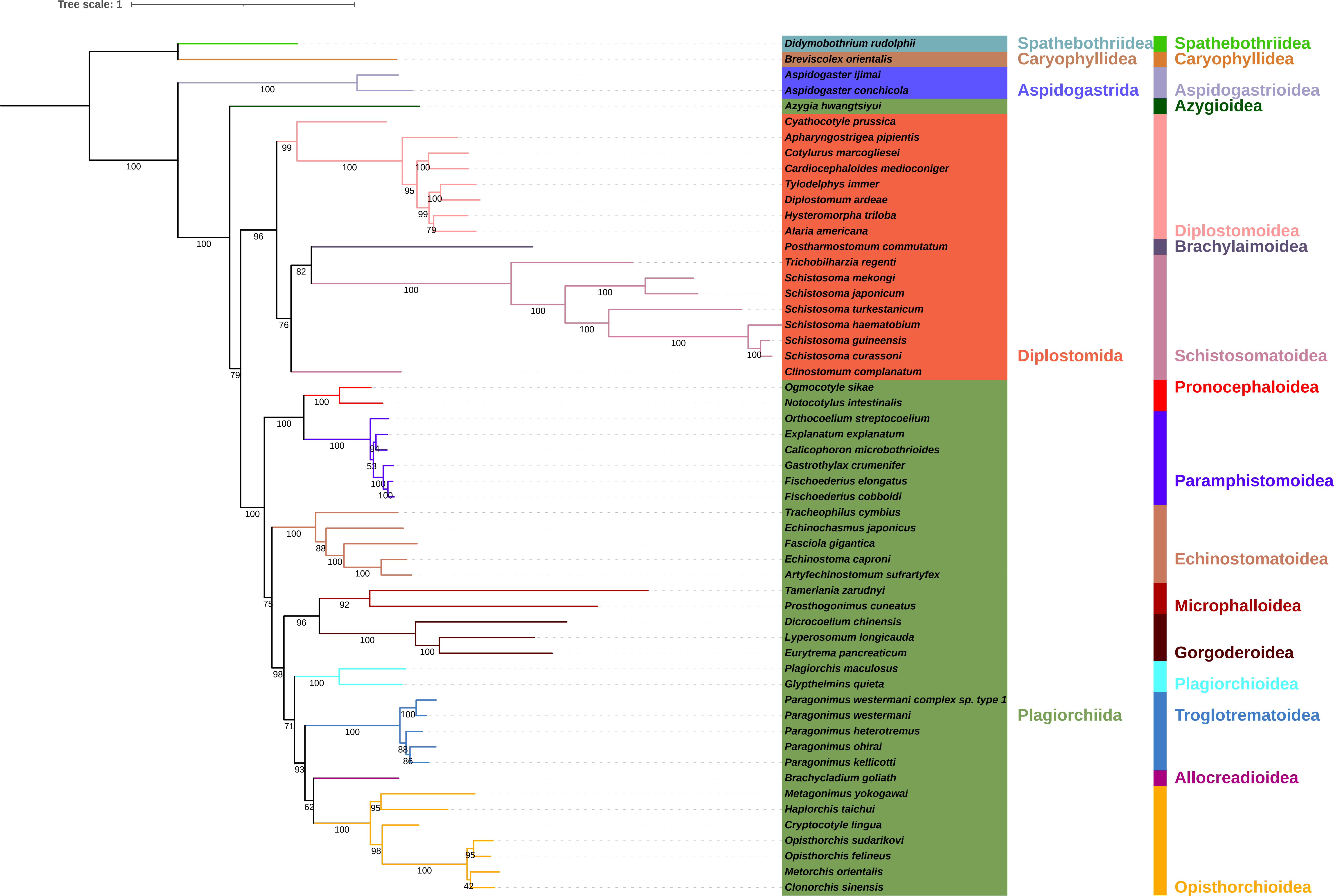

### S5-AASYMm-Ho.pdf

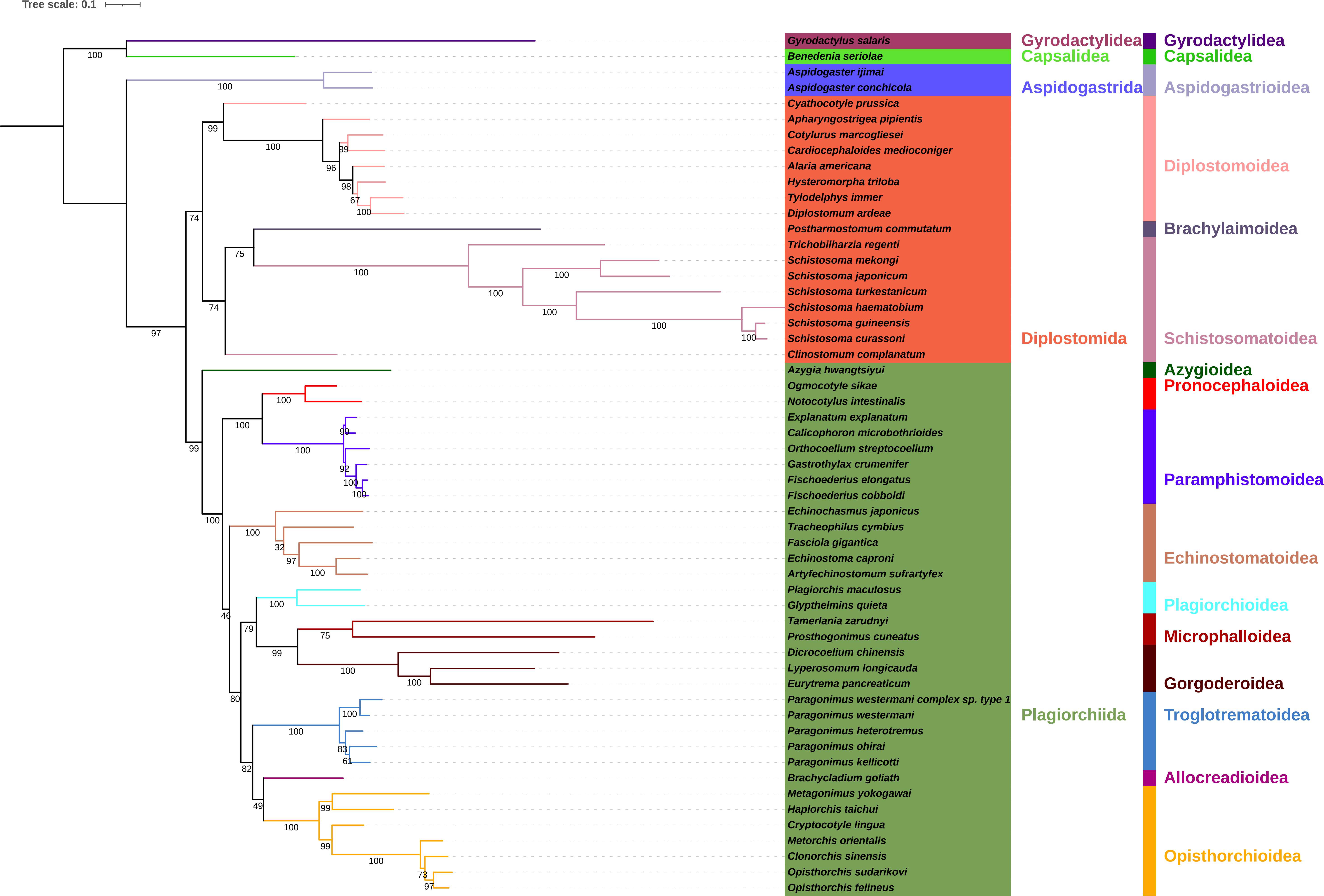

### S6-NUCSYMm-Ho.pdf

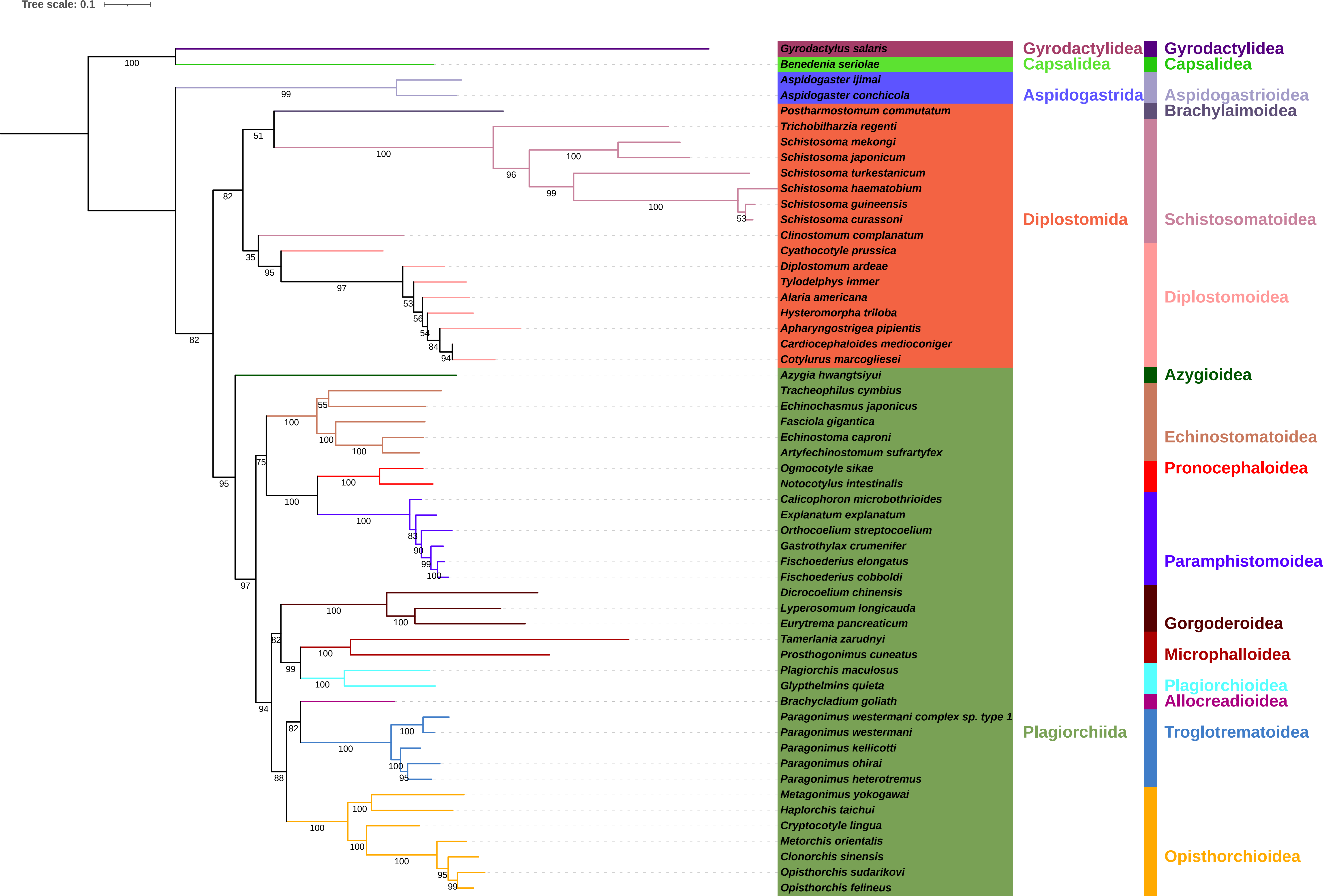

### S7-NUCSYMc-Ho.pdf

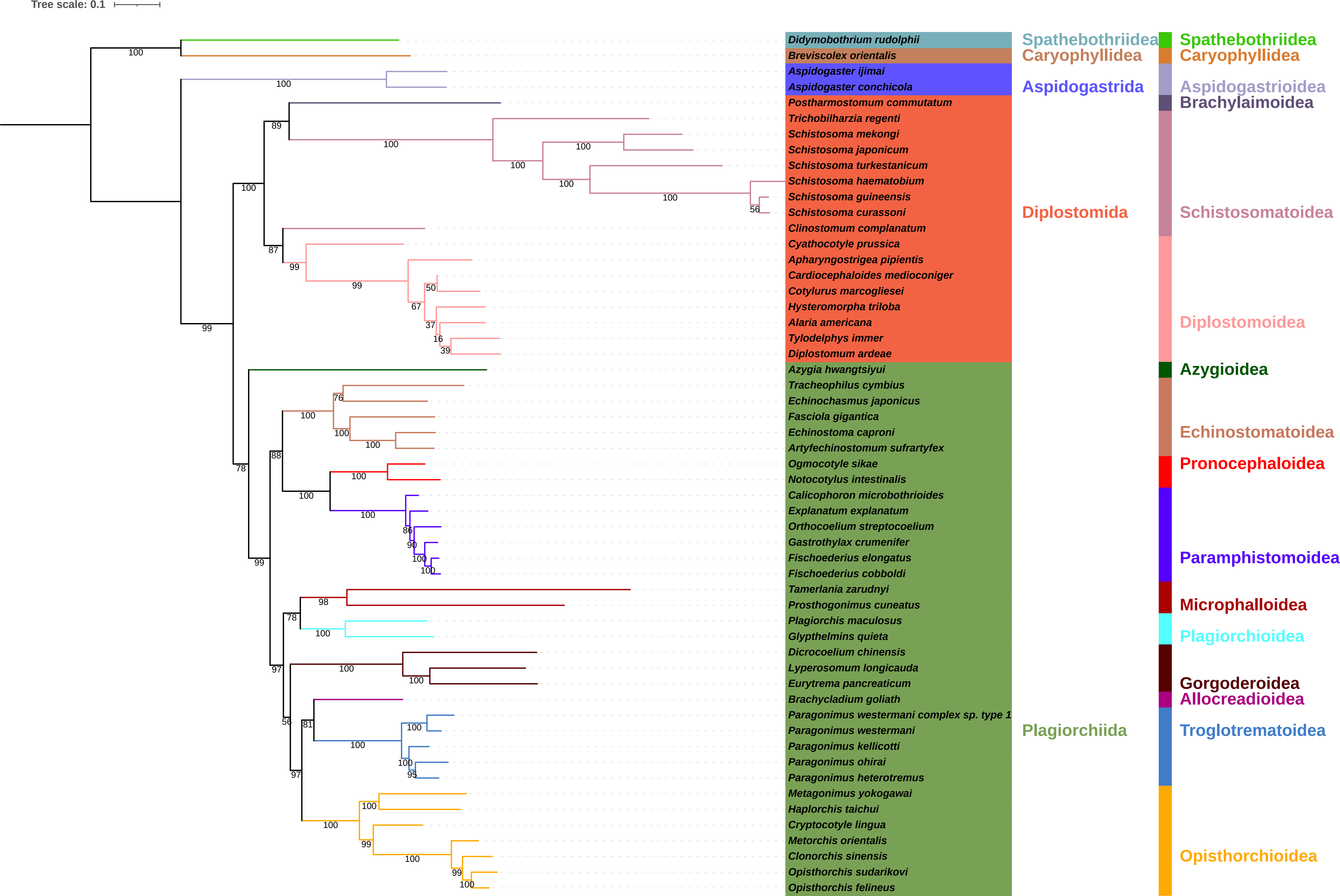

### S8-AASYMc-Ho.pdf

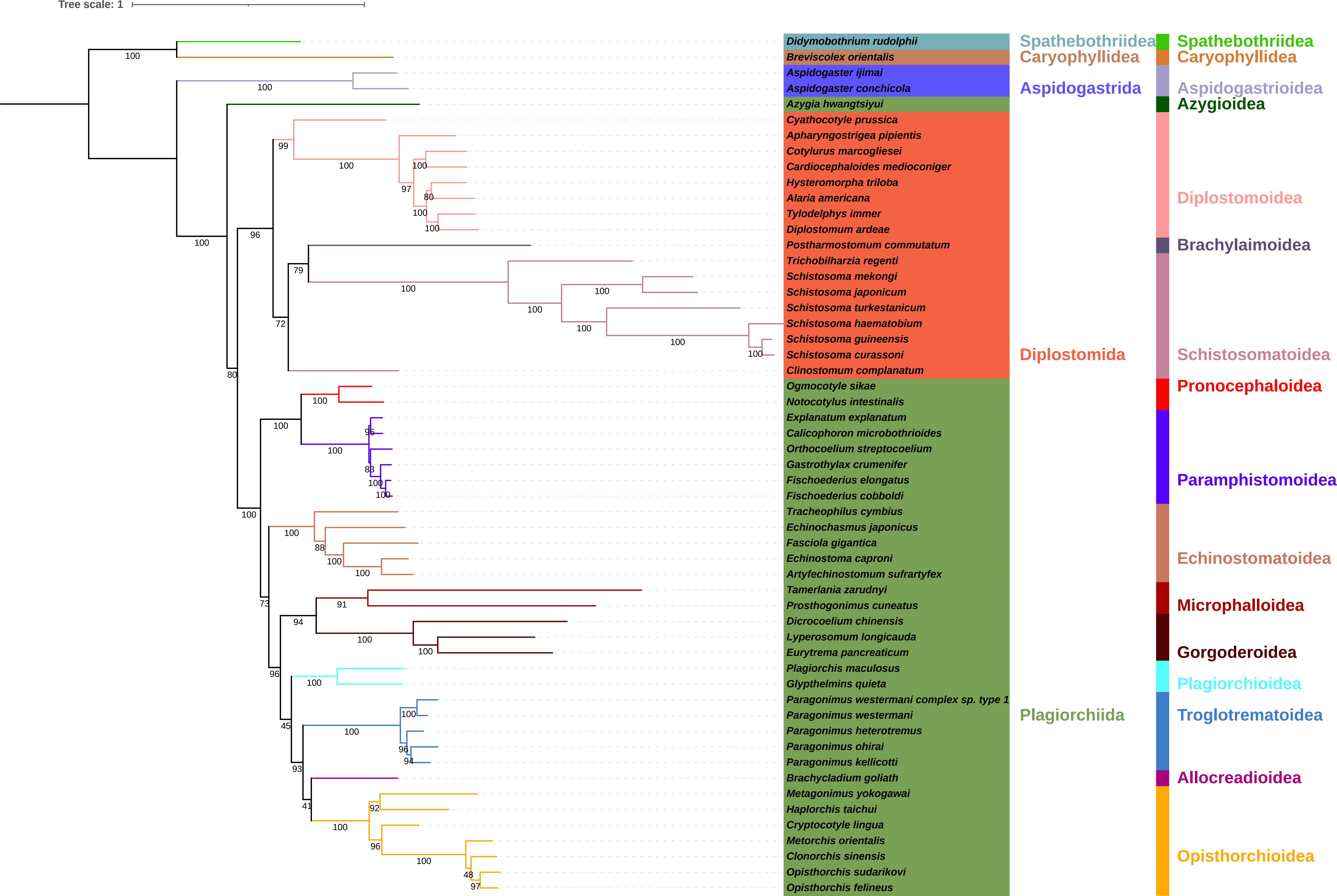

### S9-AASYMm-He.pdf

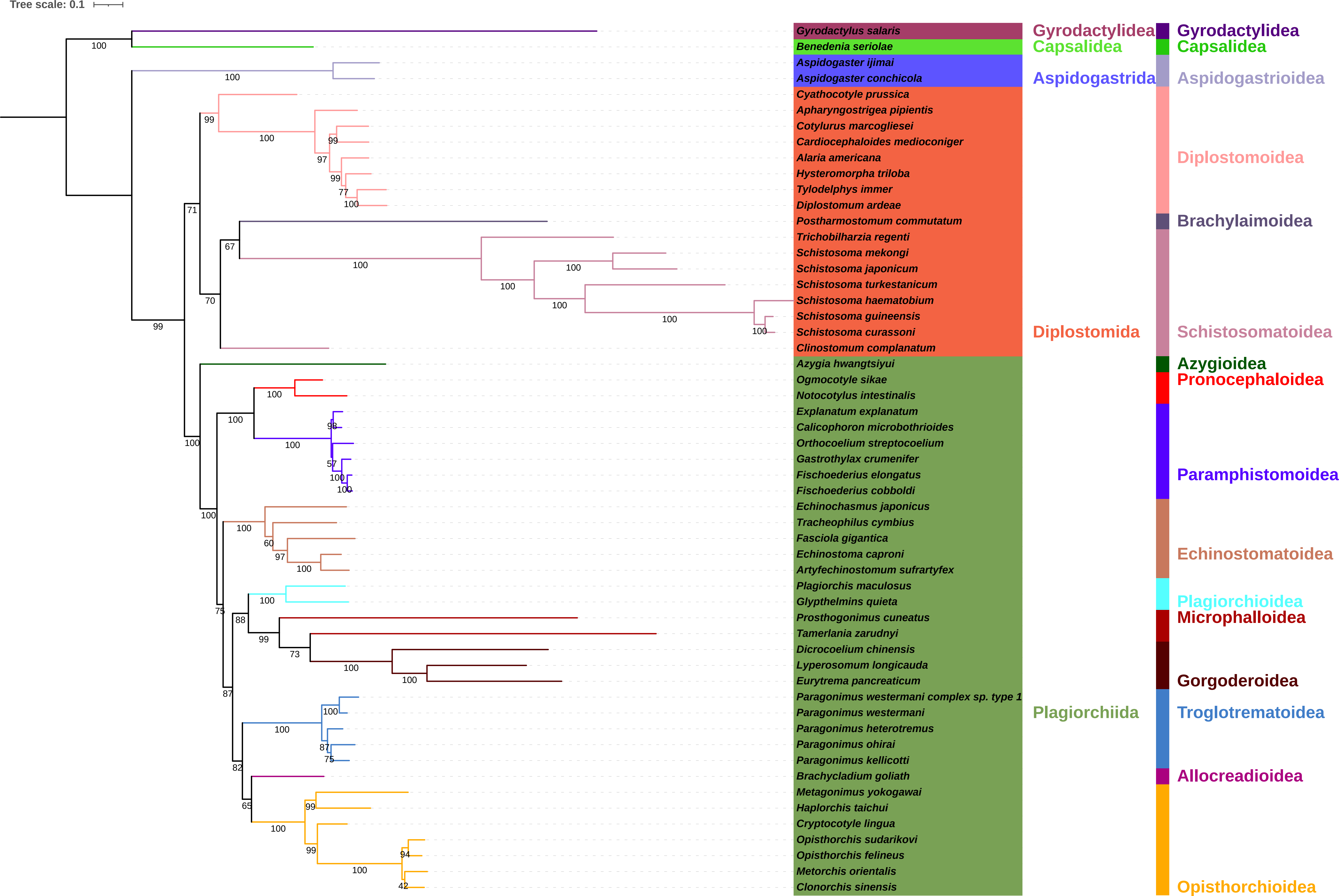

### S10-AAIQc-He.pdf

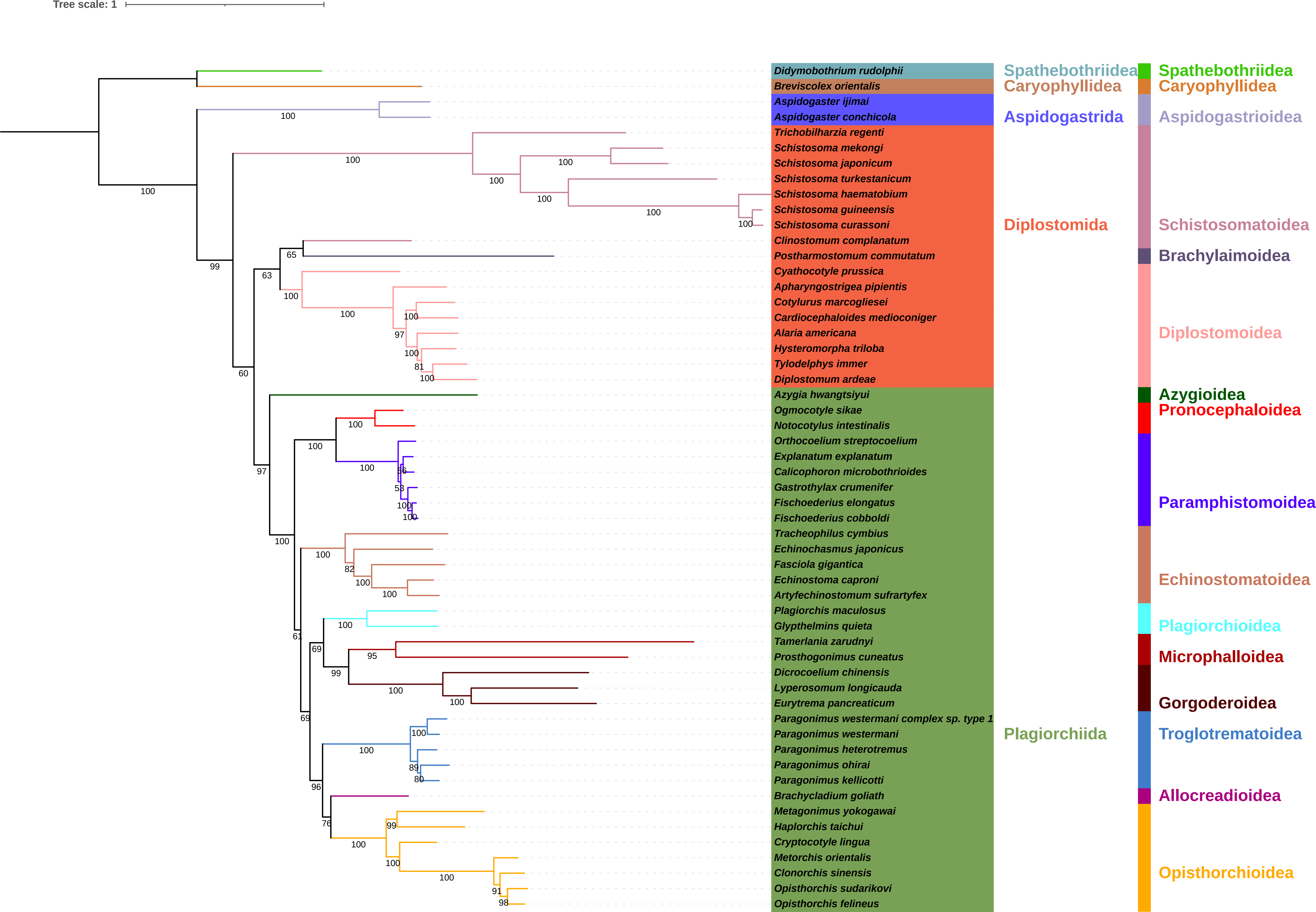

### S11-NUCm-cons.pdf

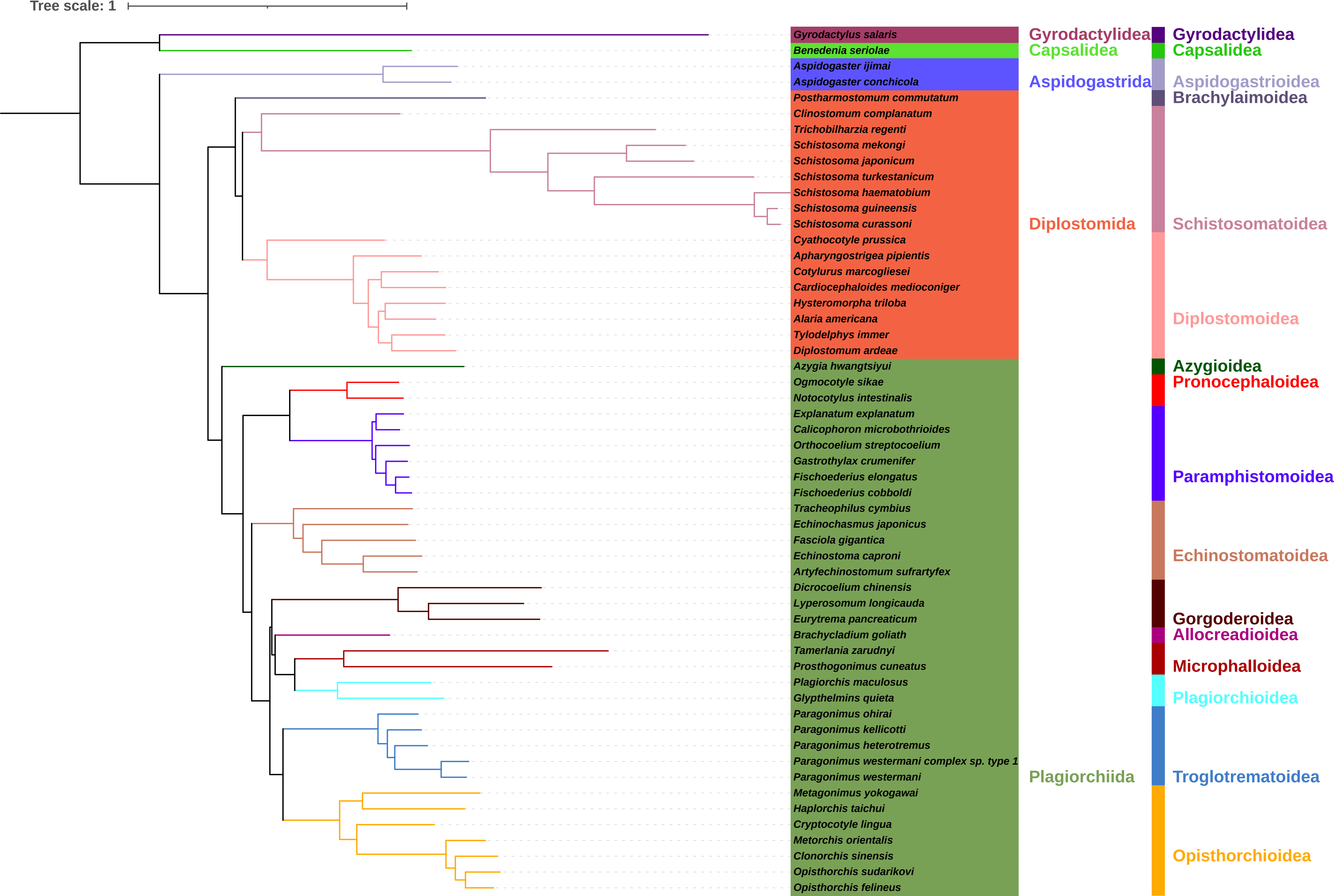

### S12-AAm-cons.pdf

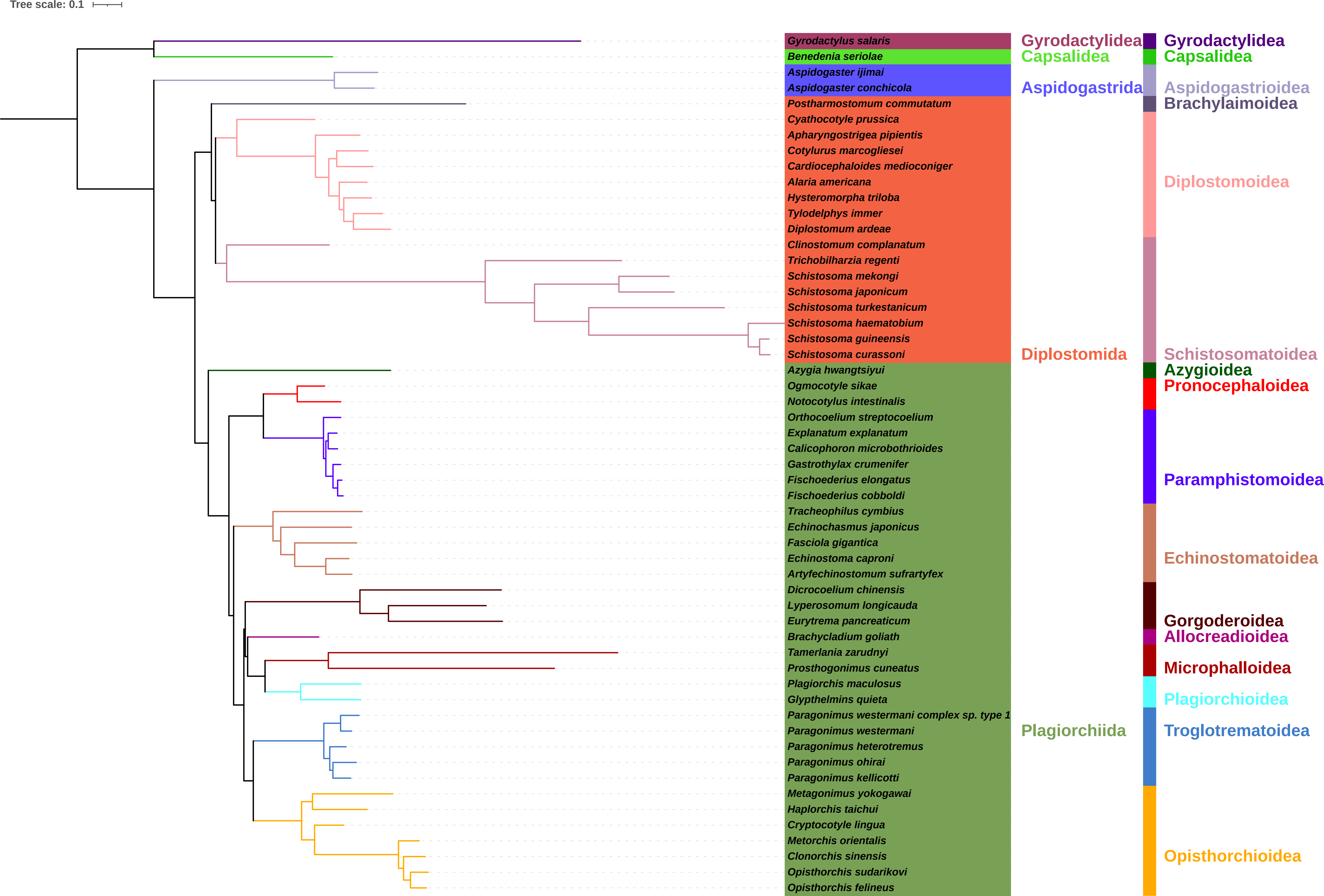

### S13-NUCIQmp.pdf

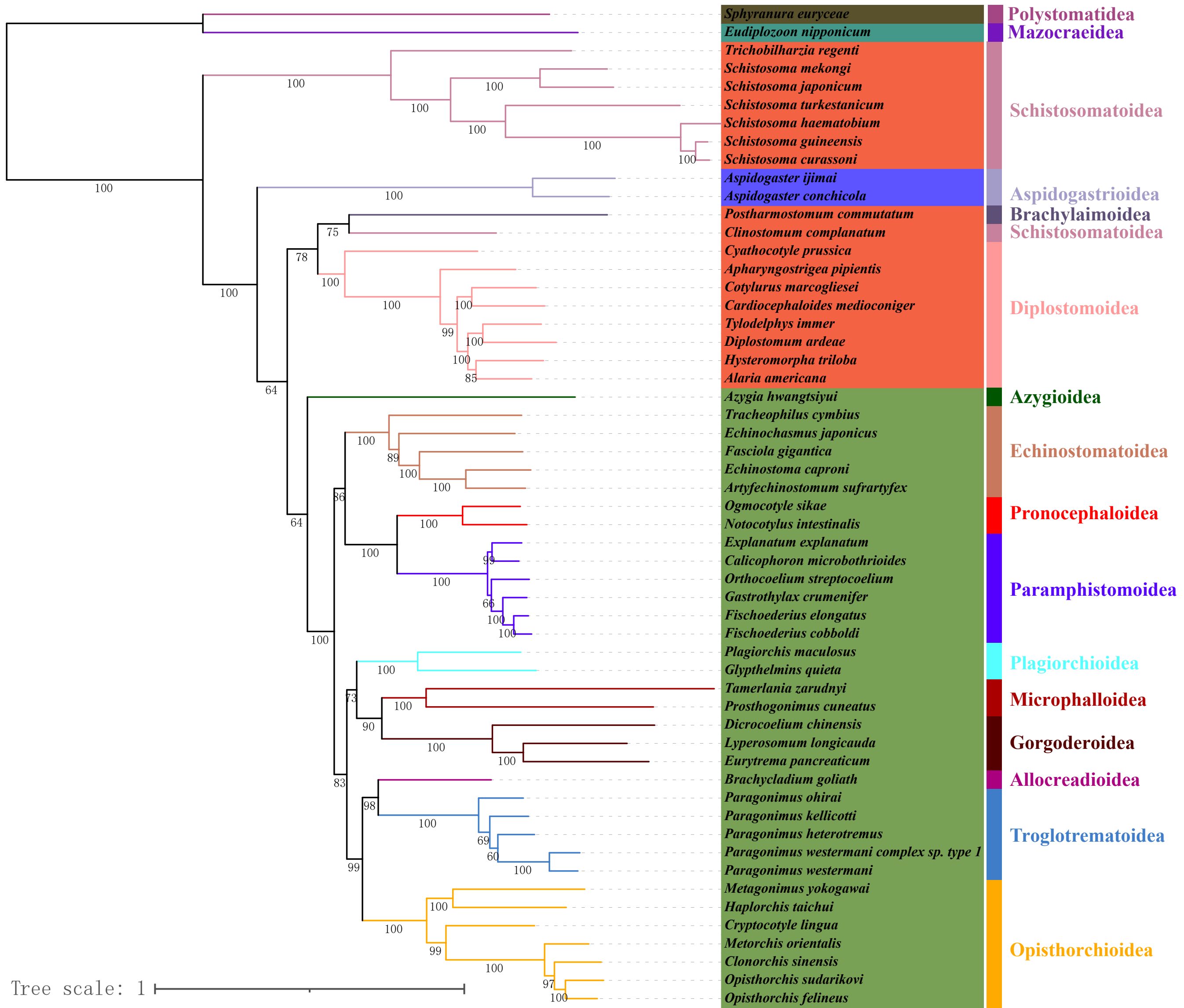

### S14-NUCPBmp.pdf

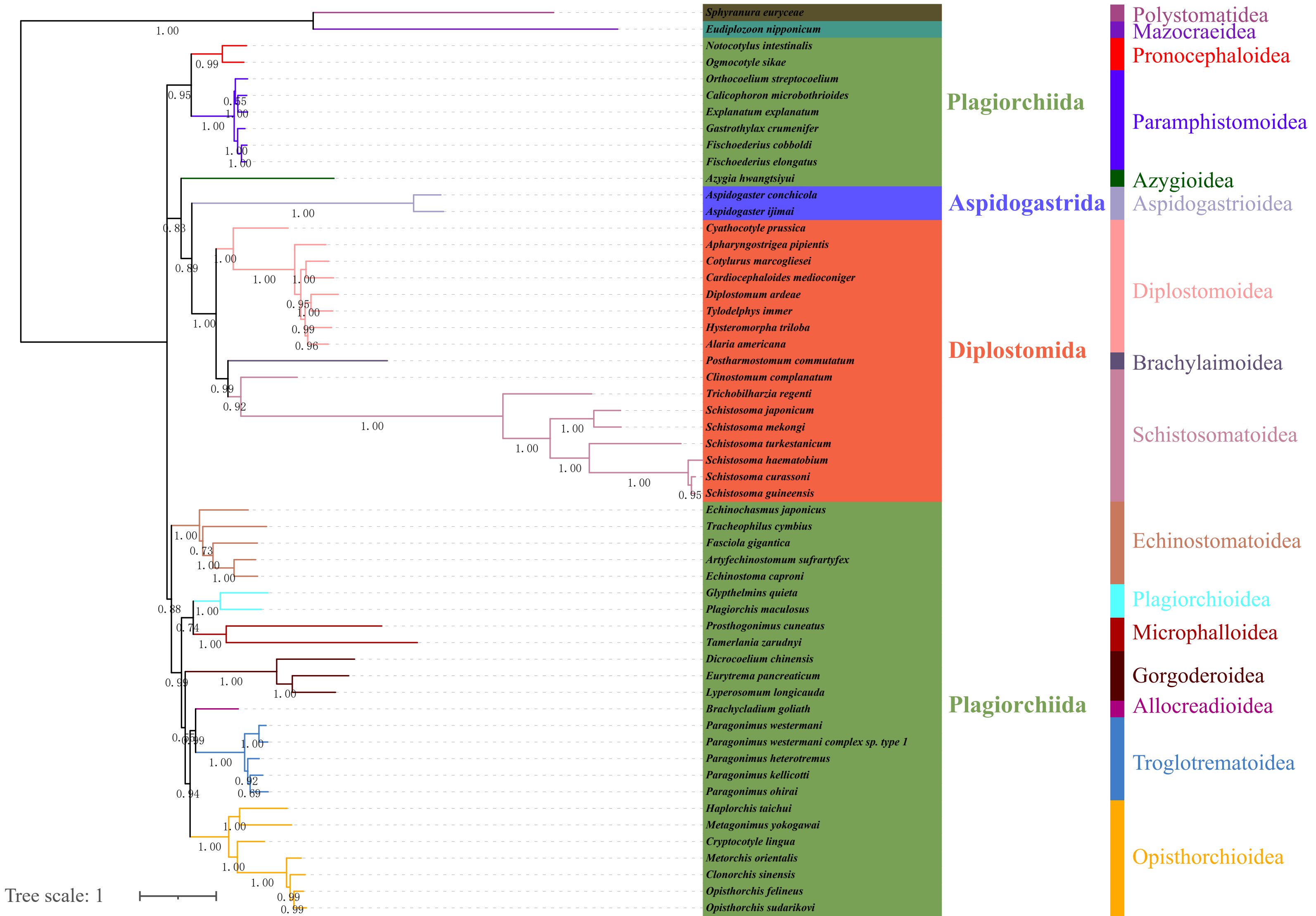

### S15-NUCPBm-sCF.pdf

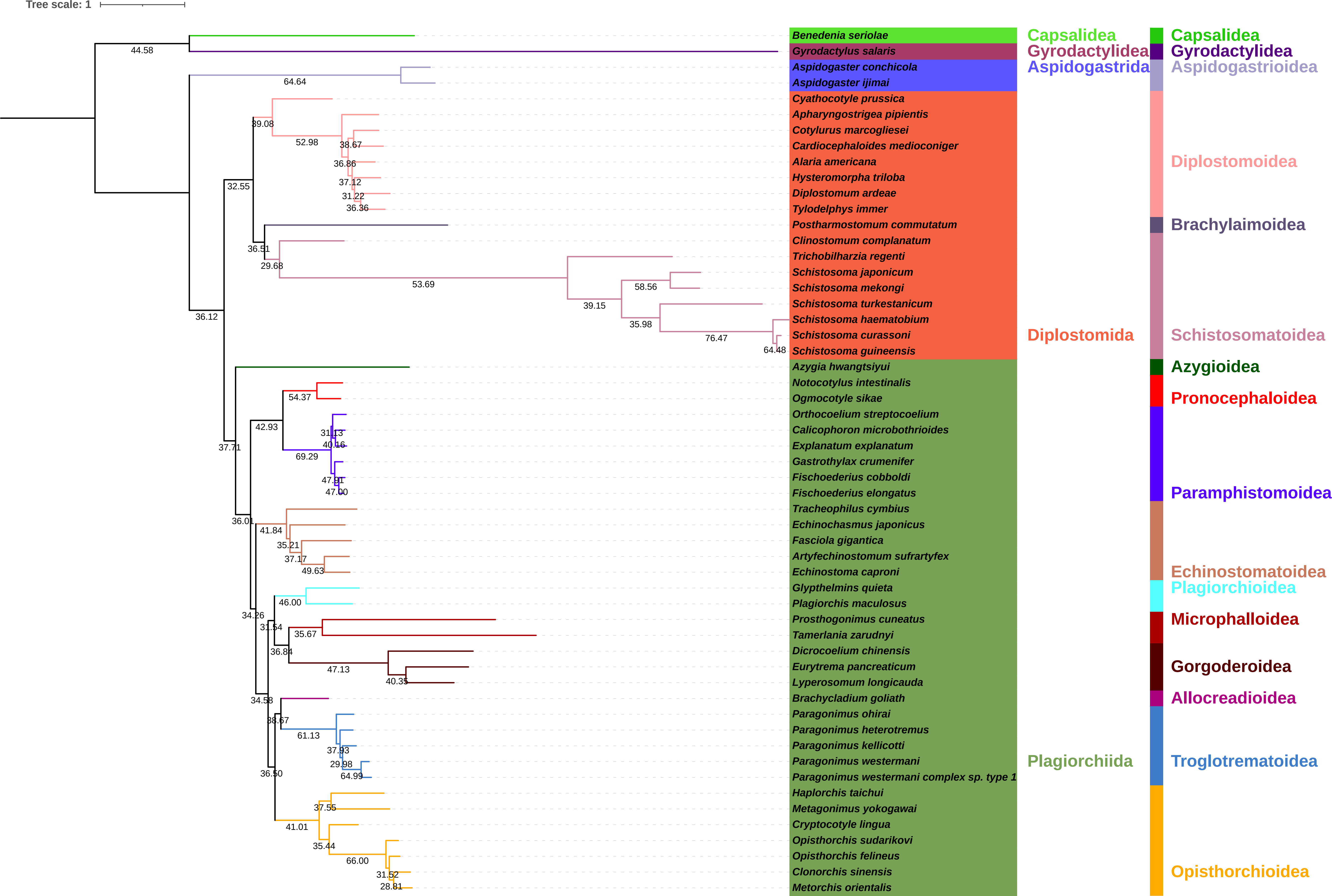

### S16-NUCPBm-gCF.pdf

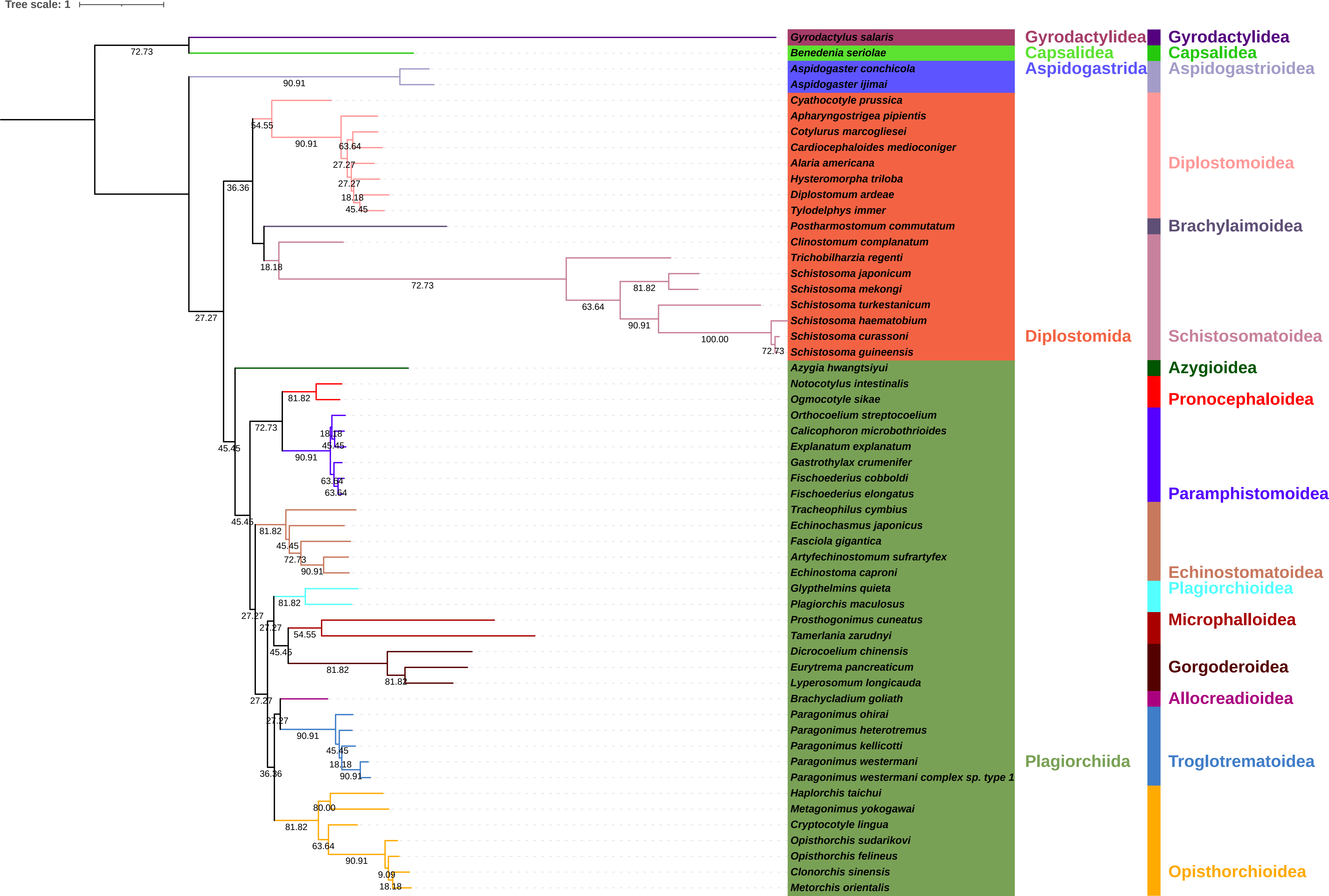

### S17-NUCPBc-sCF.pdf

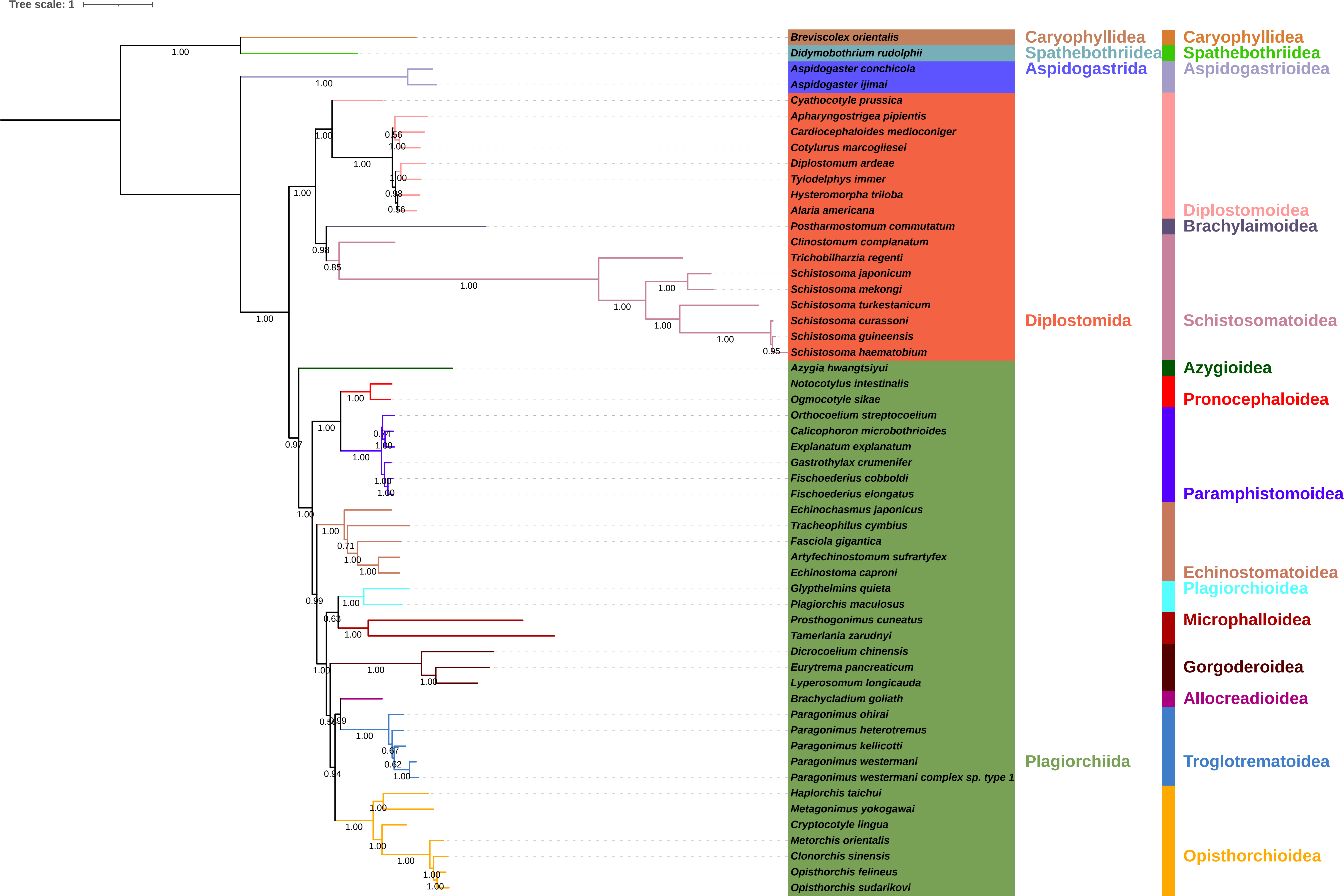

### S18-NUCPBc-gCF.pdf

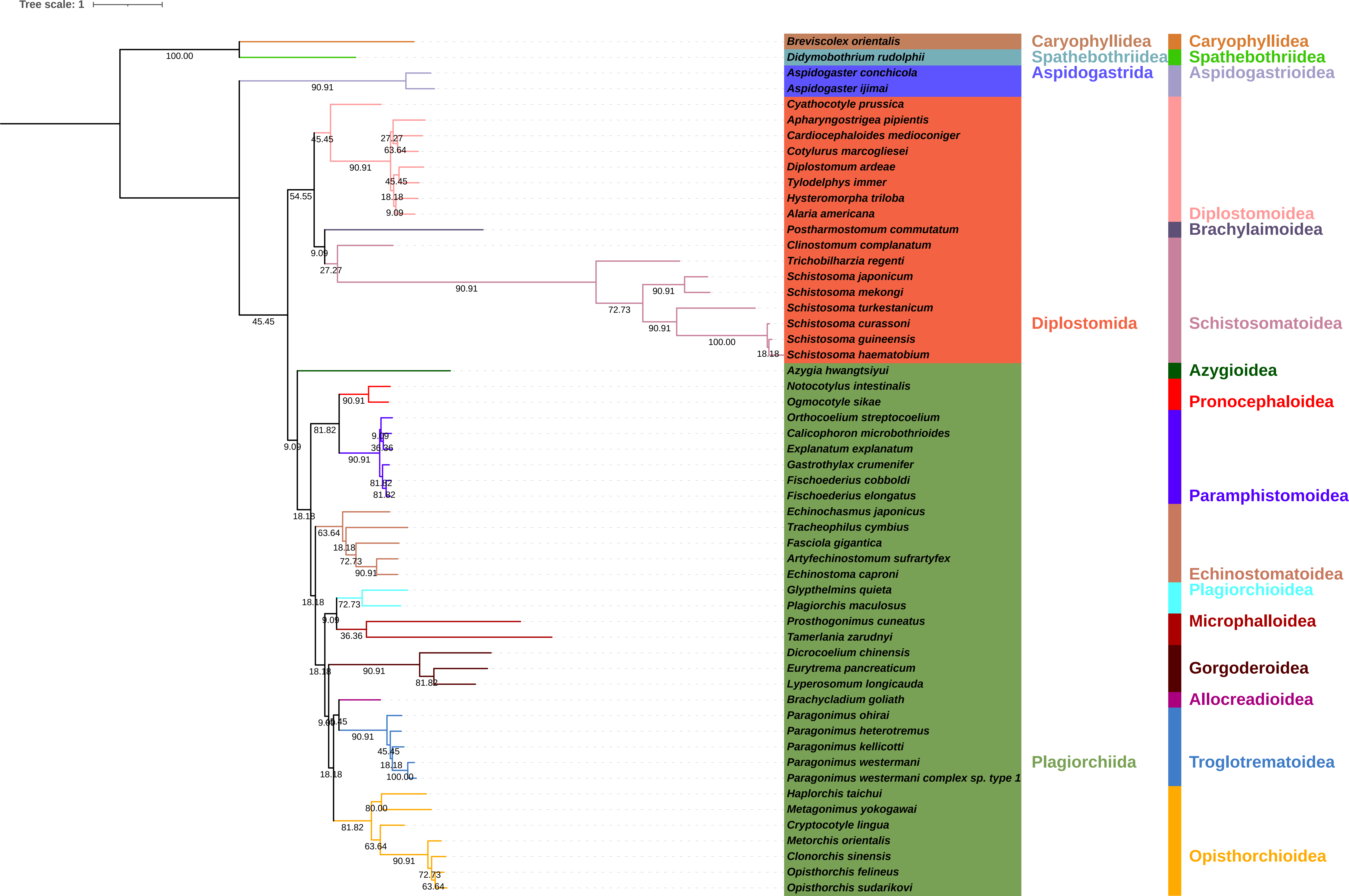

### S19-NUCc-AL-cons.pdf

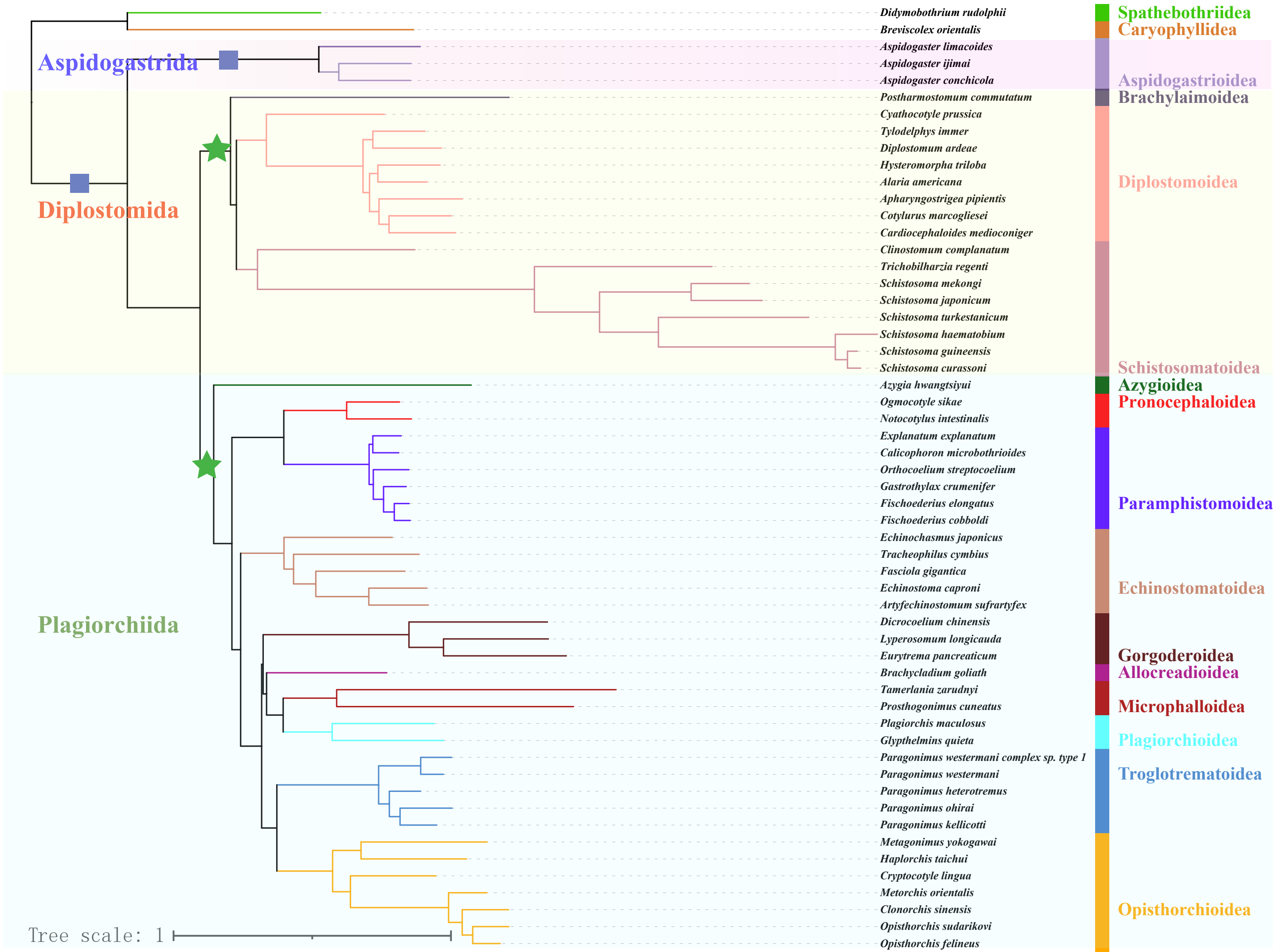

### S20-AAIQm.pdf

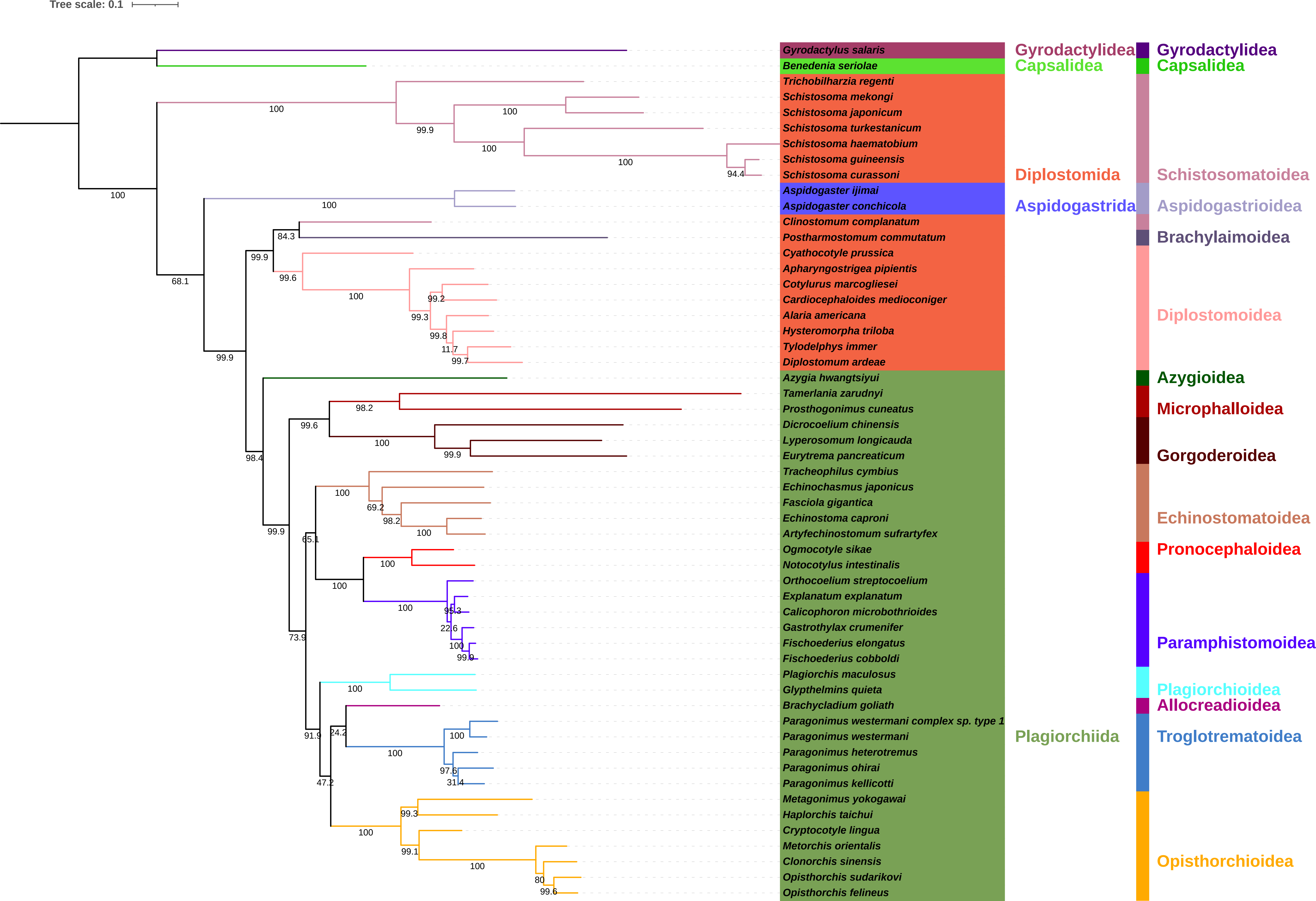

### S21-NUCIQm.pdf

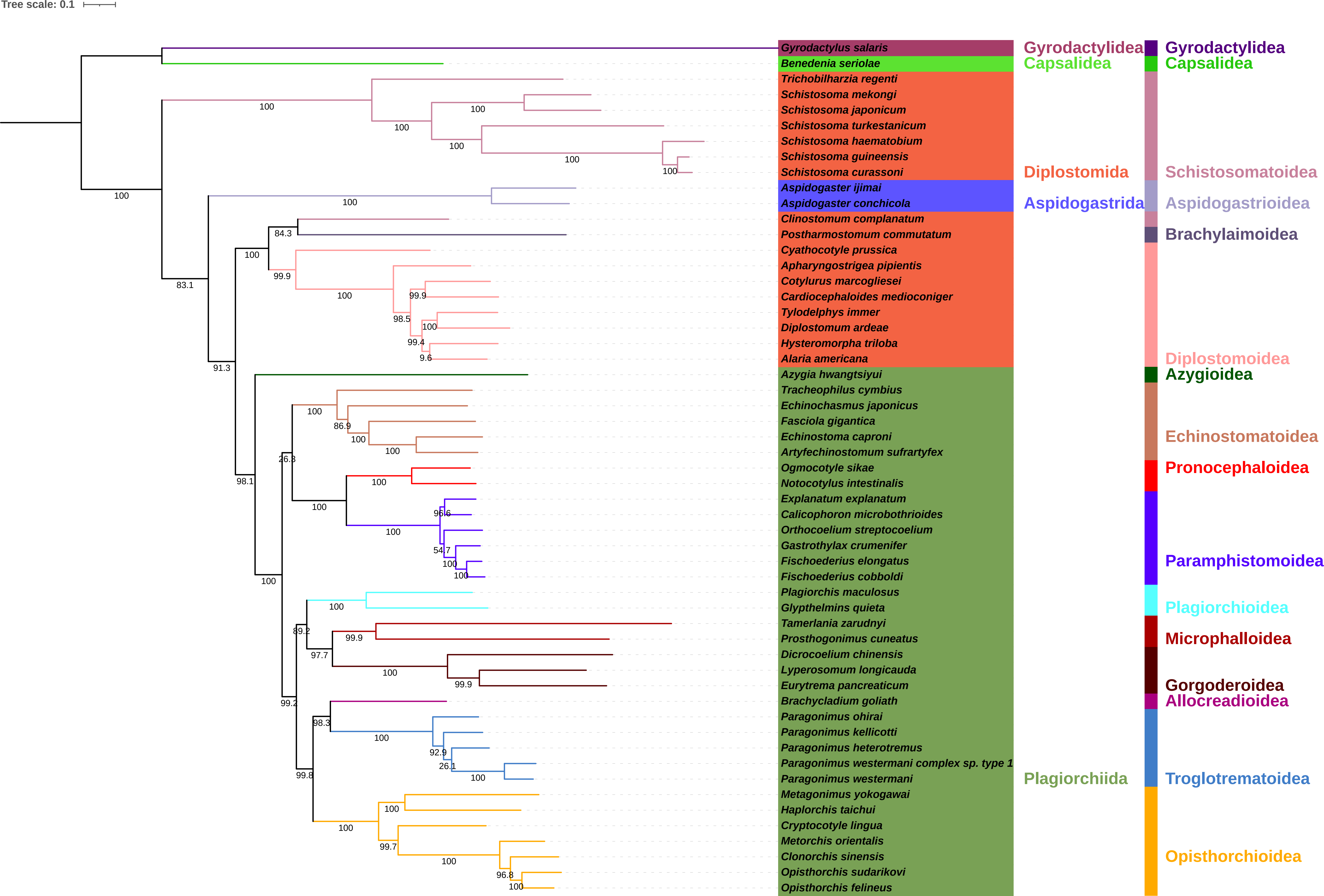

### S22-AARAm.pdf

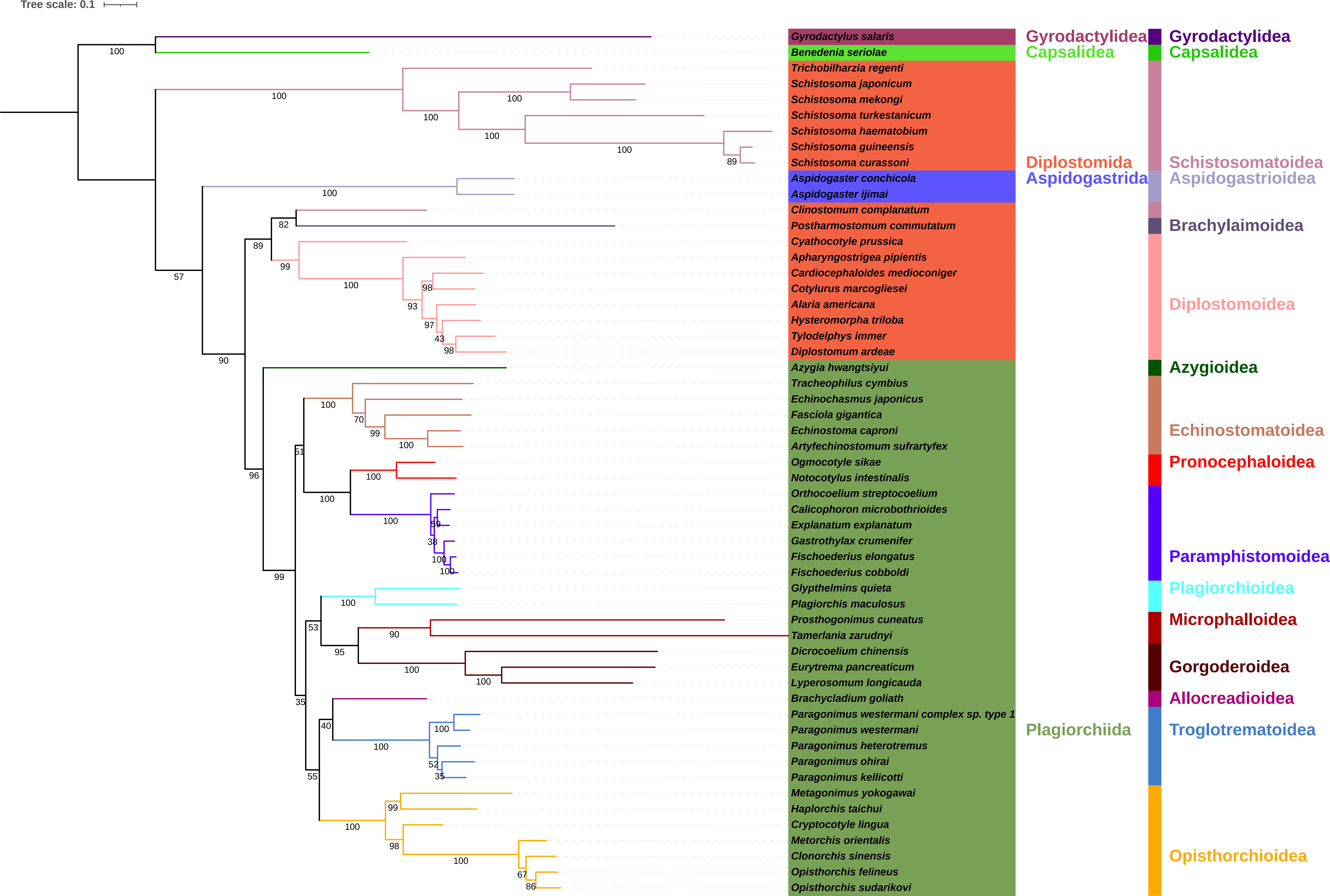

### S23-NUCRAm.pdf

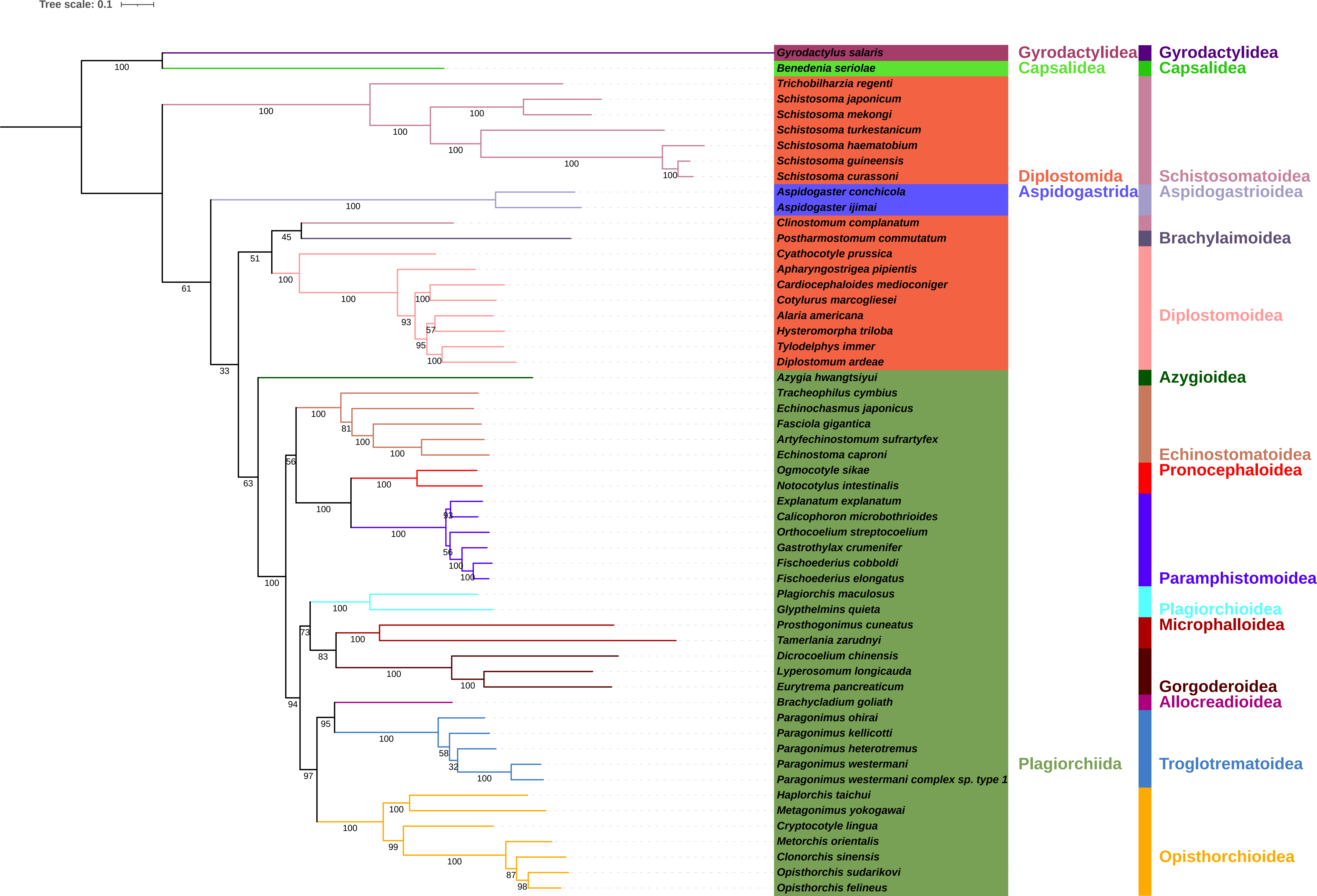

### S24-AAMBm.pdf

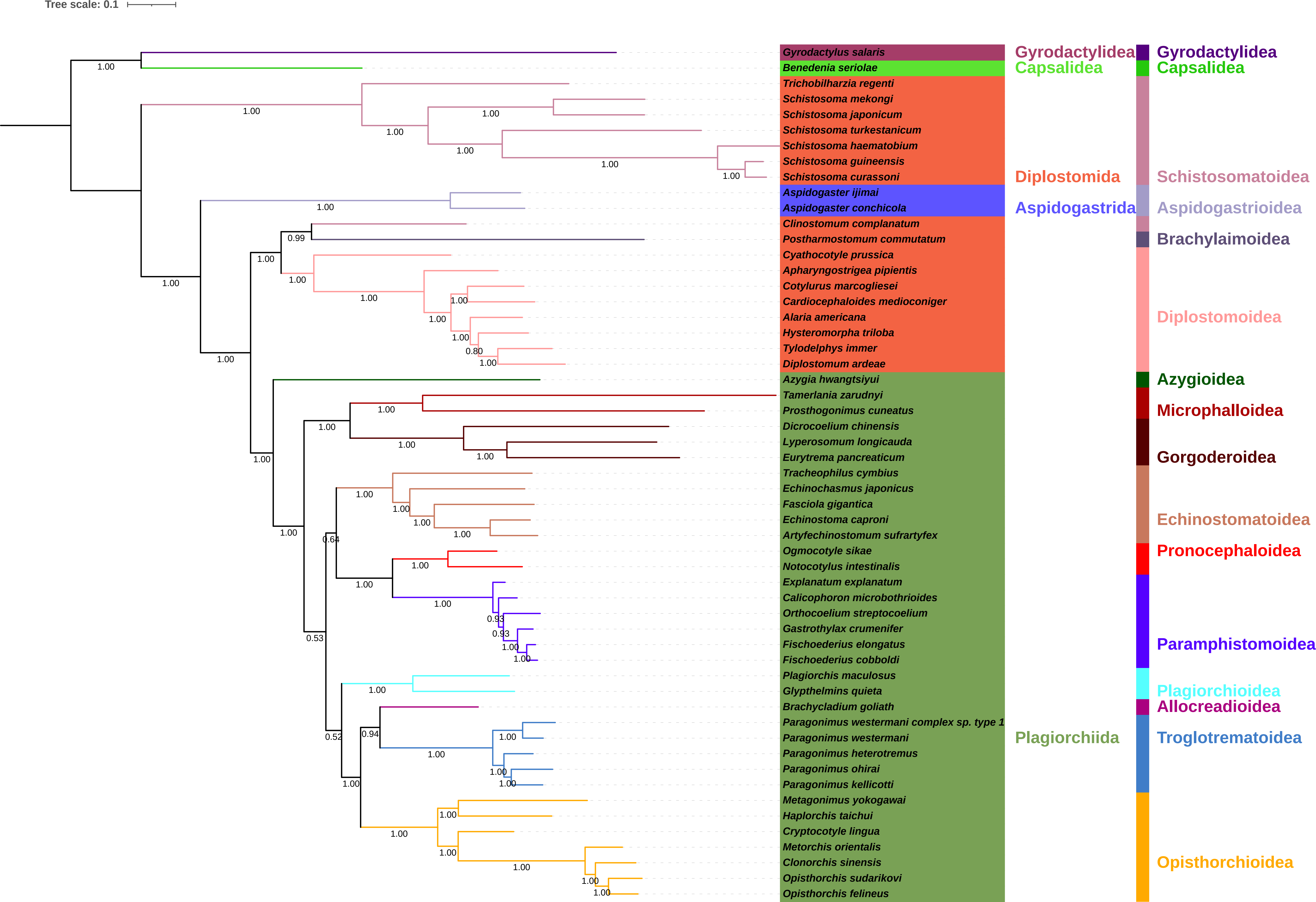

### S25-AAPBm.pdf

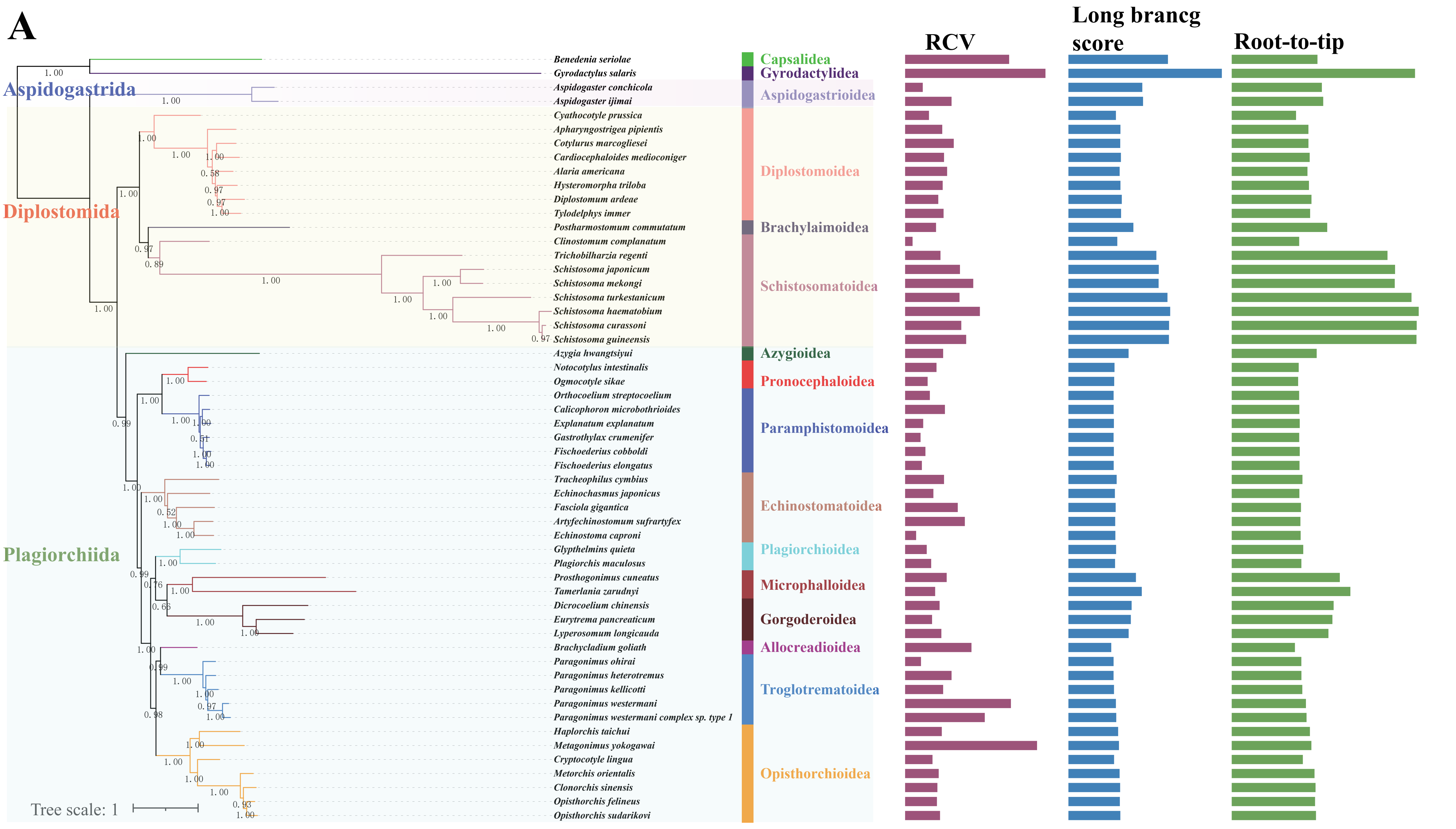

### S26-AAIQm-He.pdf

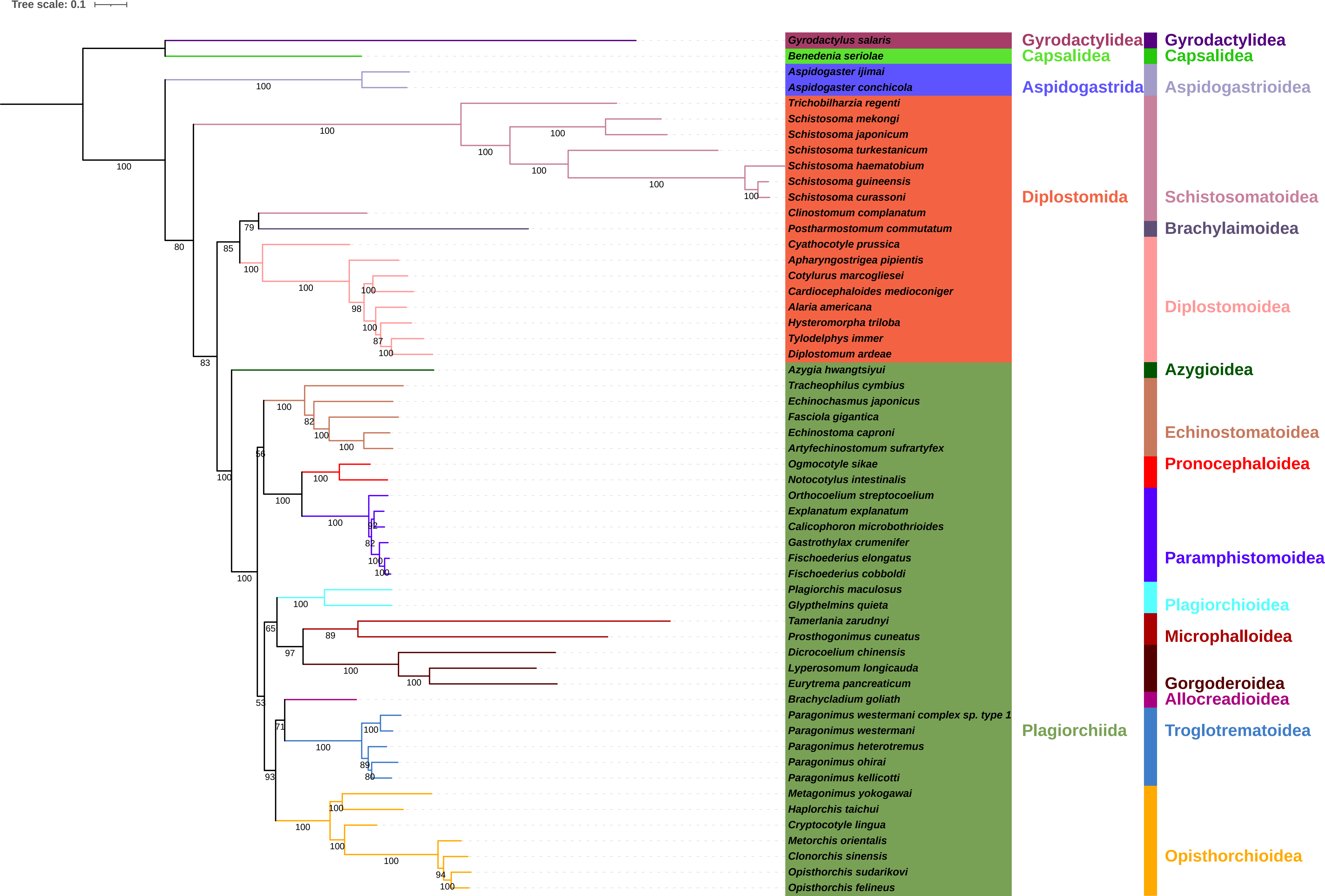

### S27-AAIQc.pdf

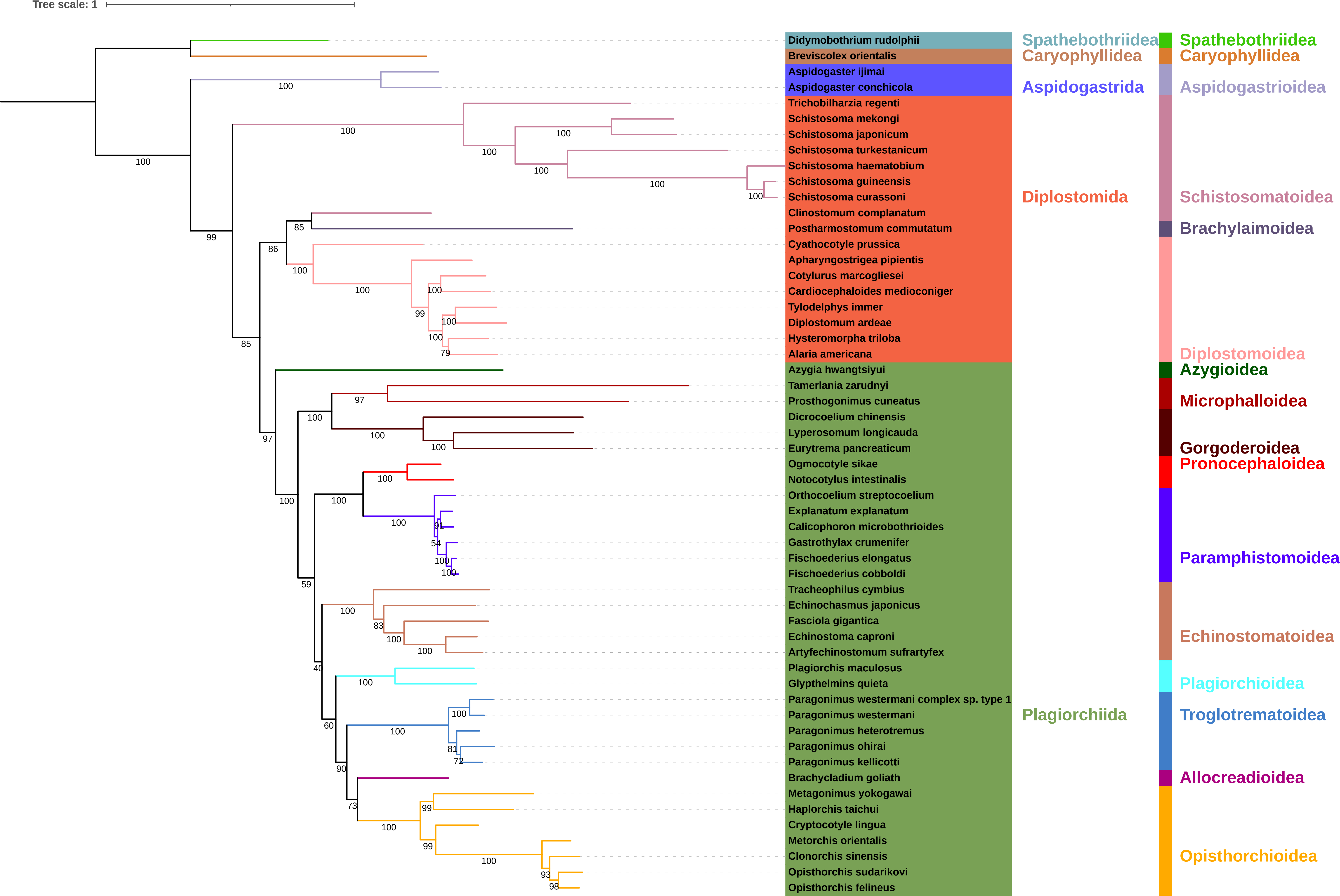

### S28-NUCIQc.pdf

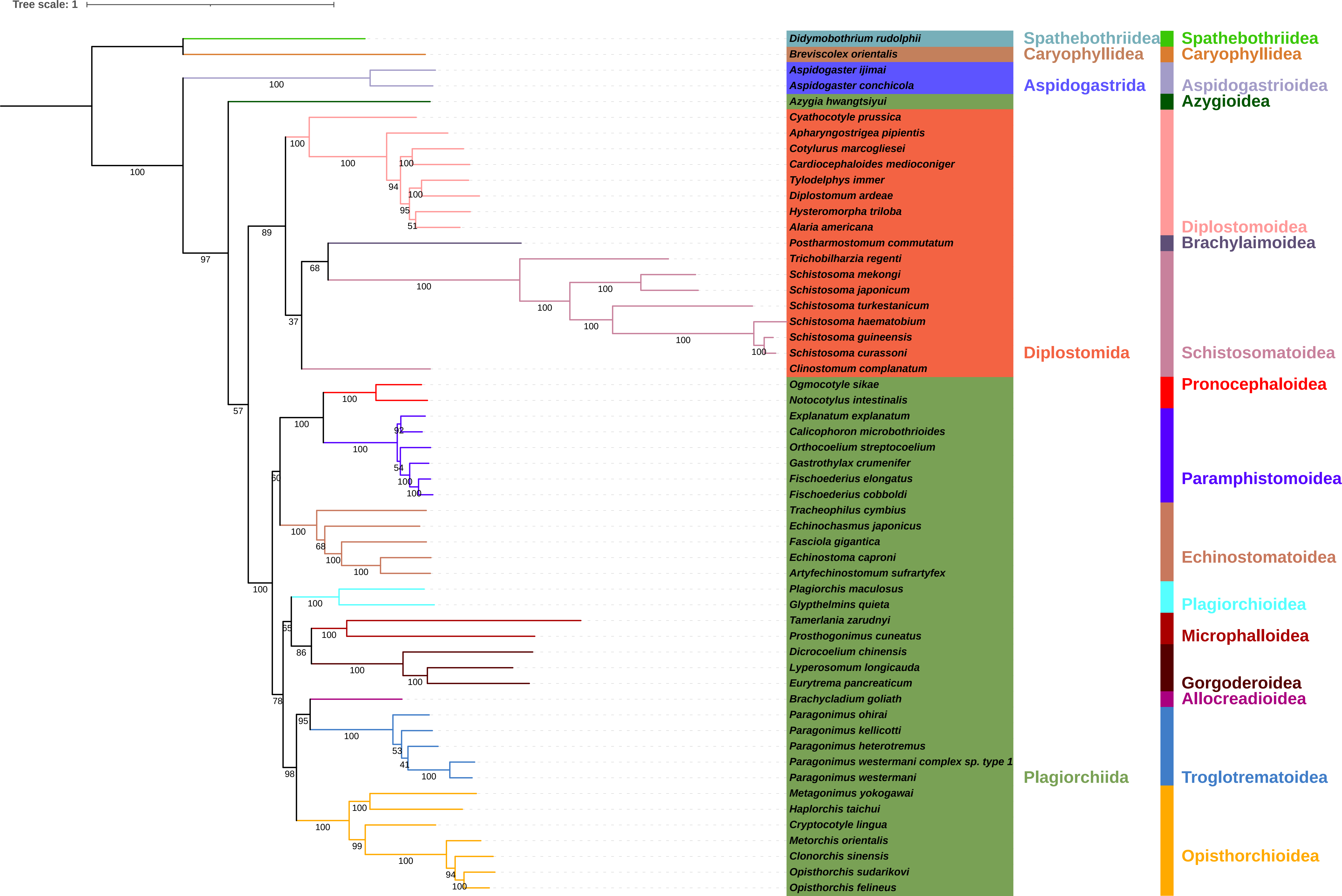

### S29-AARAc.pdf

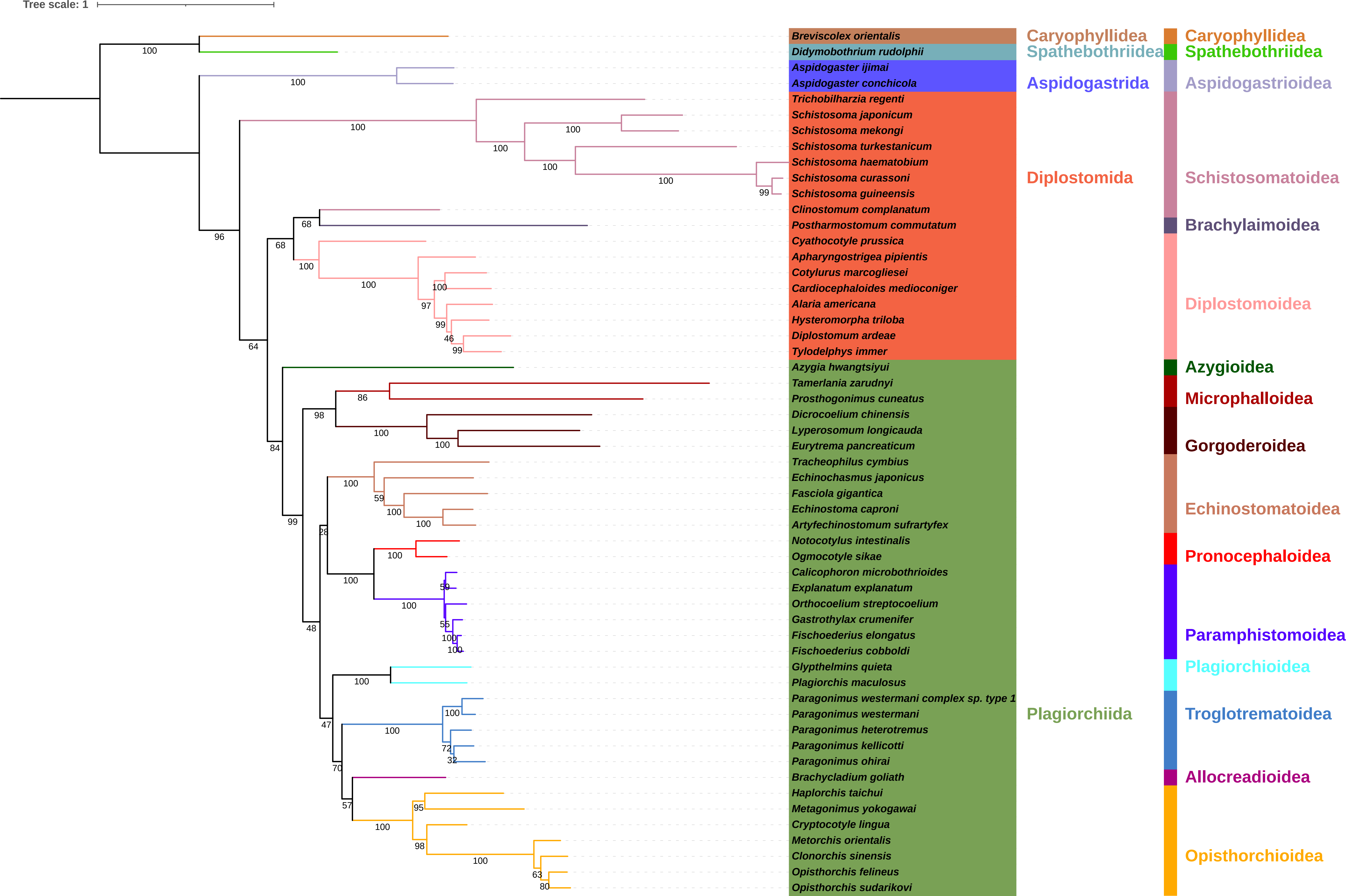

### S30-NUCRAc.pdf

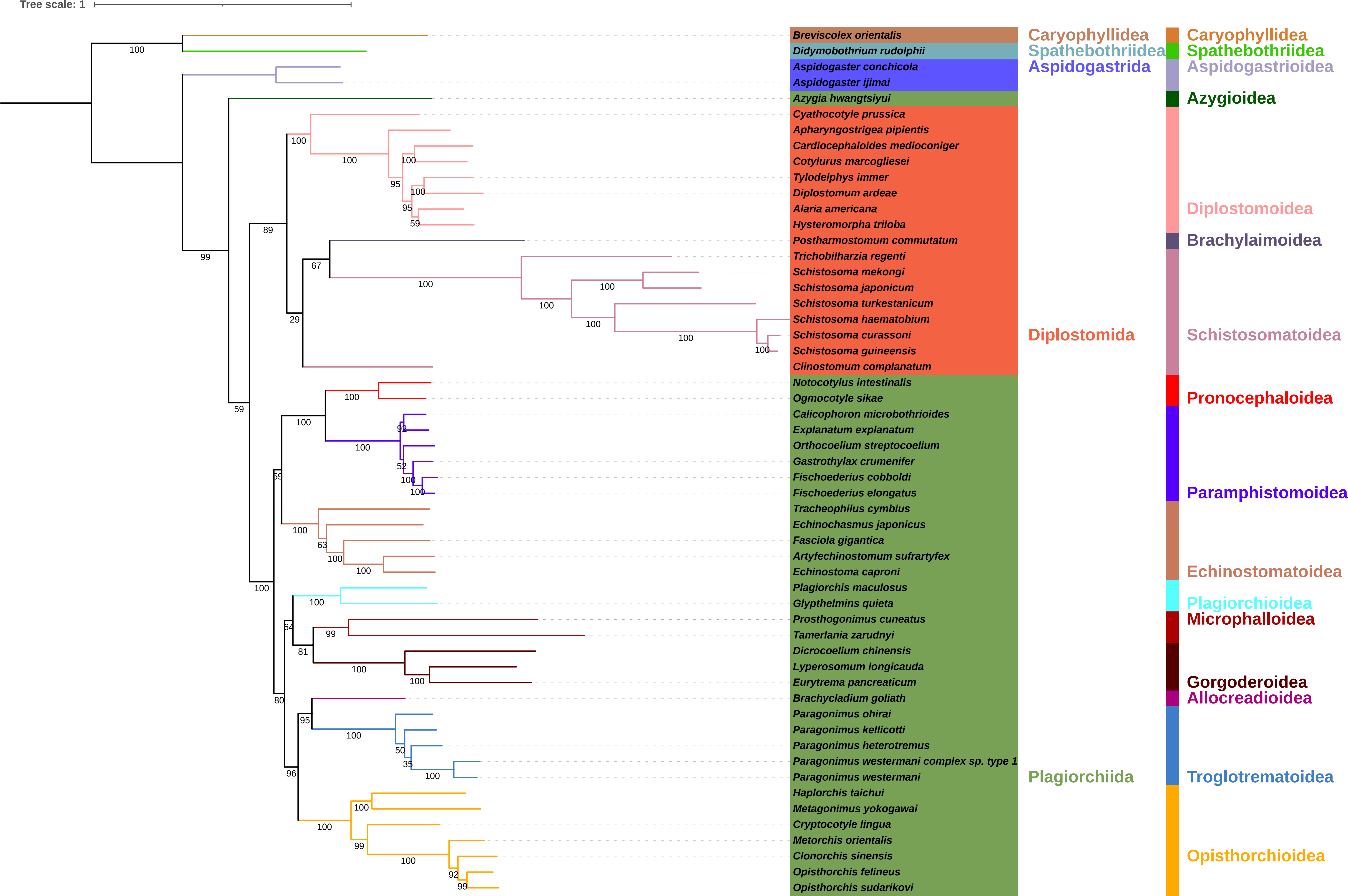

### S31-AAMBc.pdf

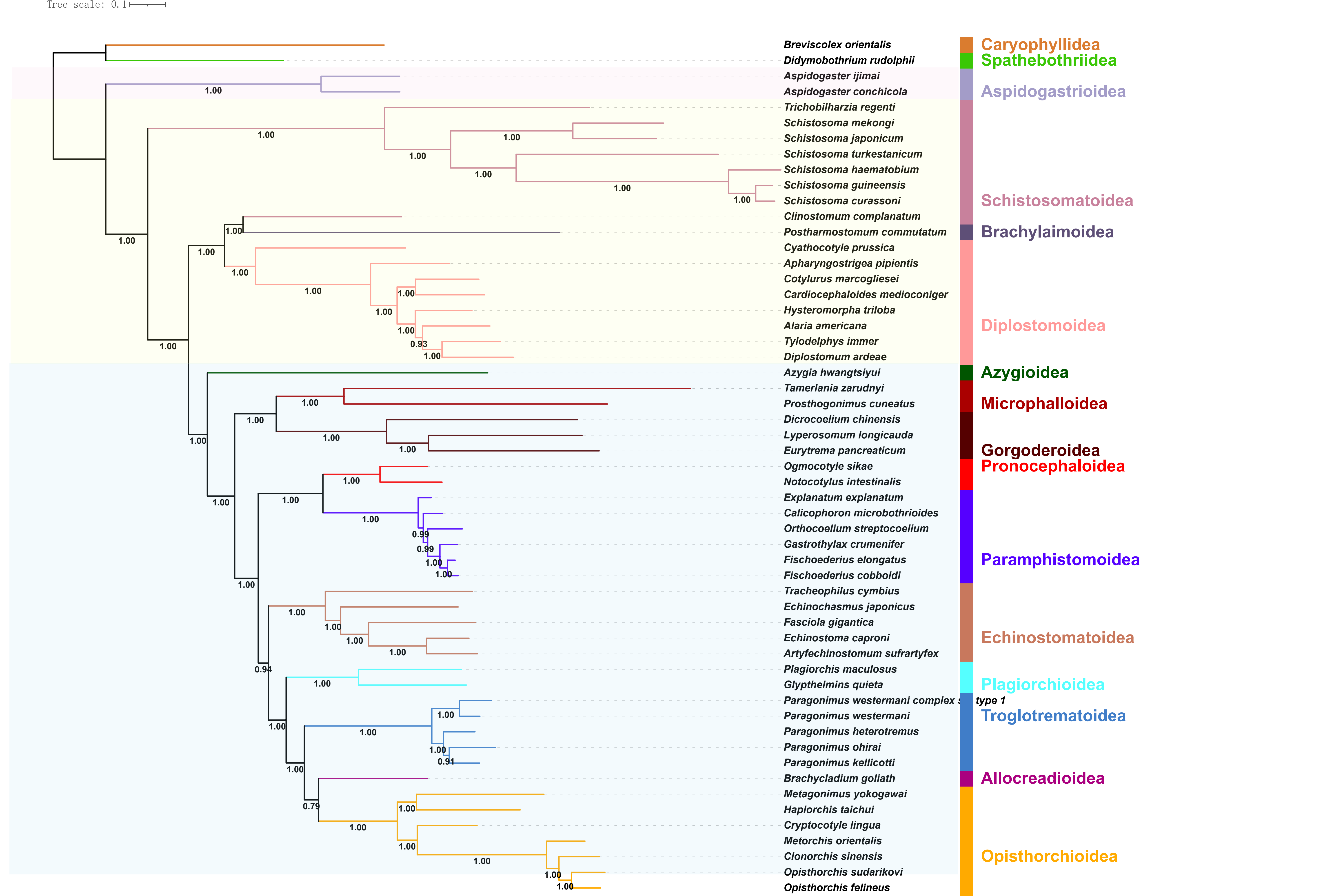

### S32-AAPBc.pdf

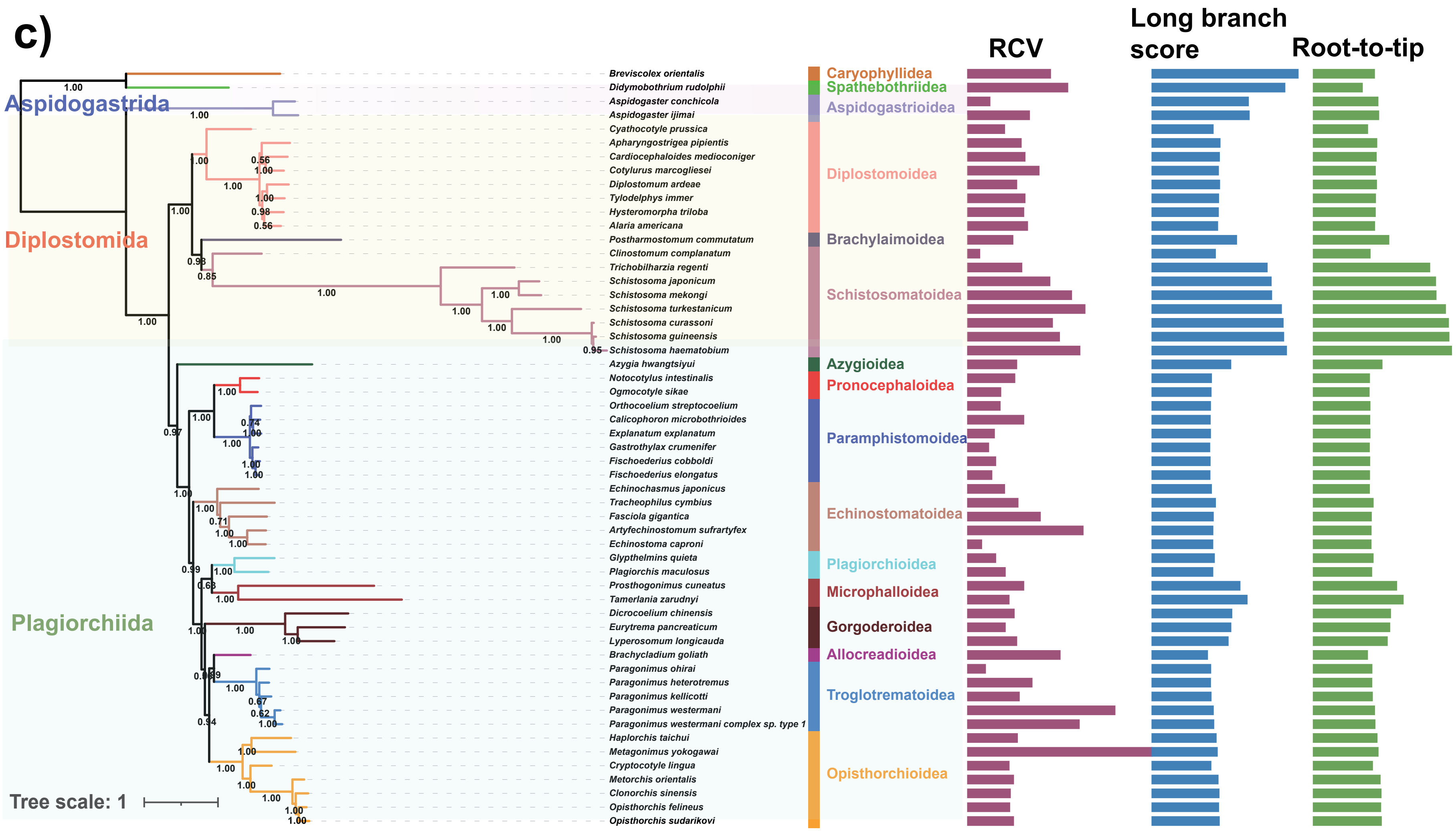

### S33-AASYMc-He.pdf

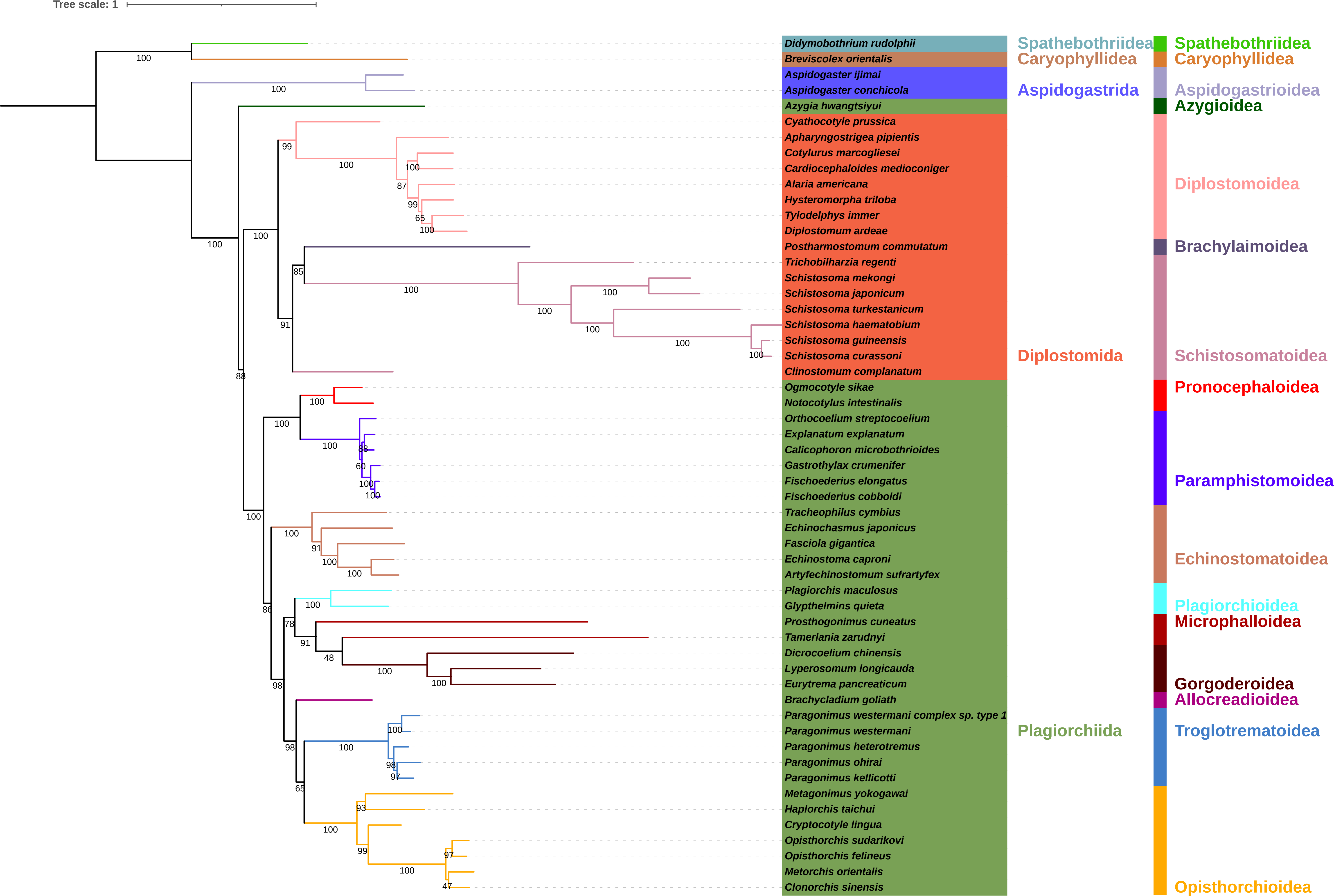

### S34-NUCIQc-sCF.pdf

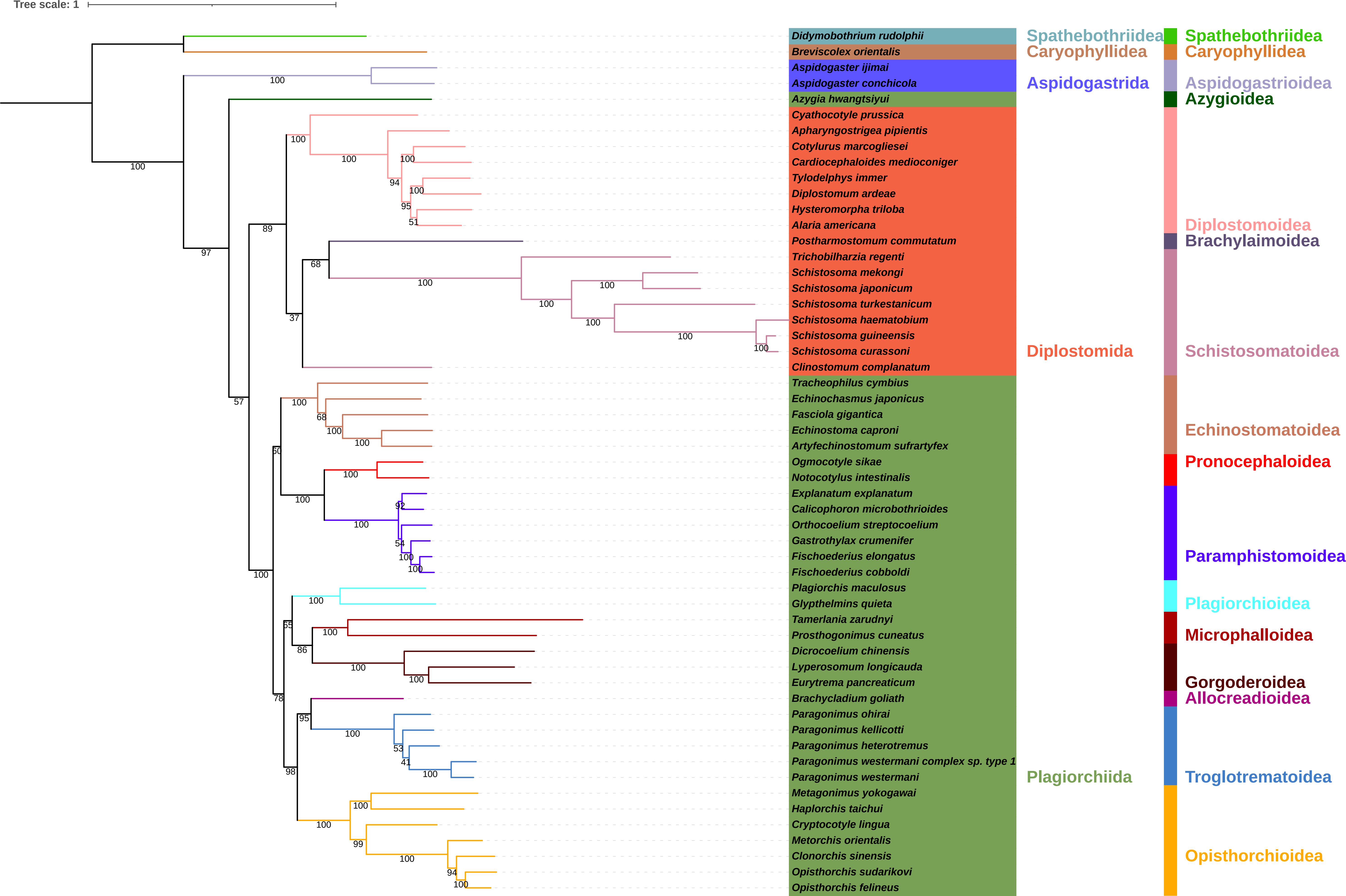

### S35-NUCIQc-gCF.pdf

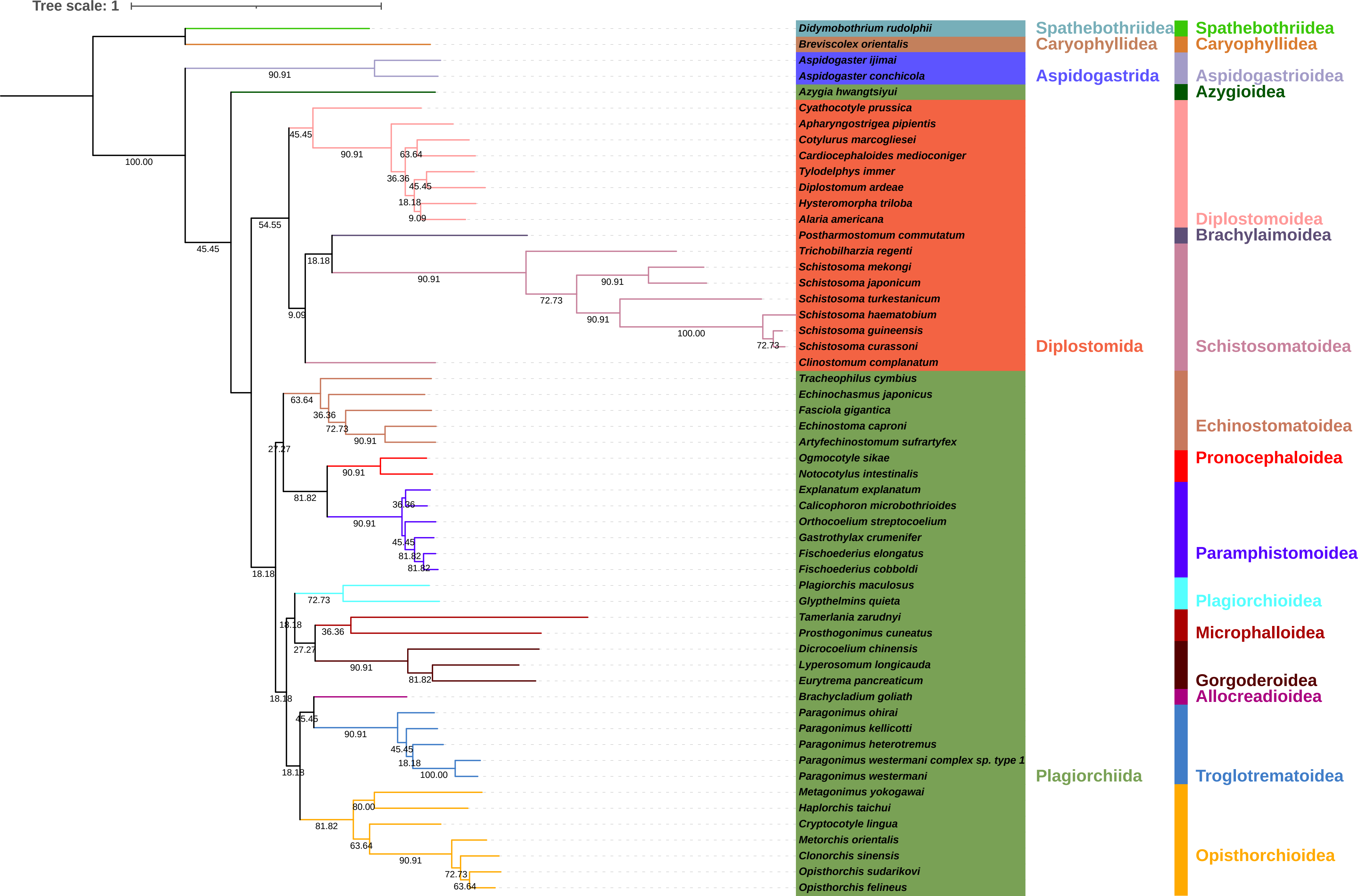

### S36-NUCIQm-sCF.pdf

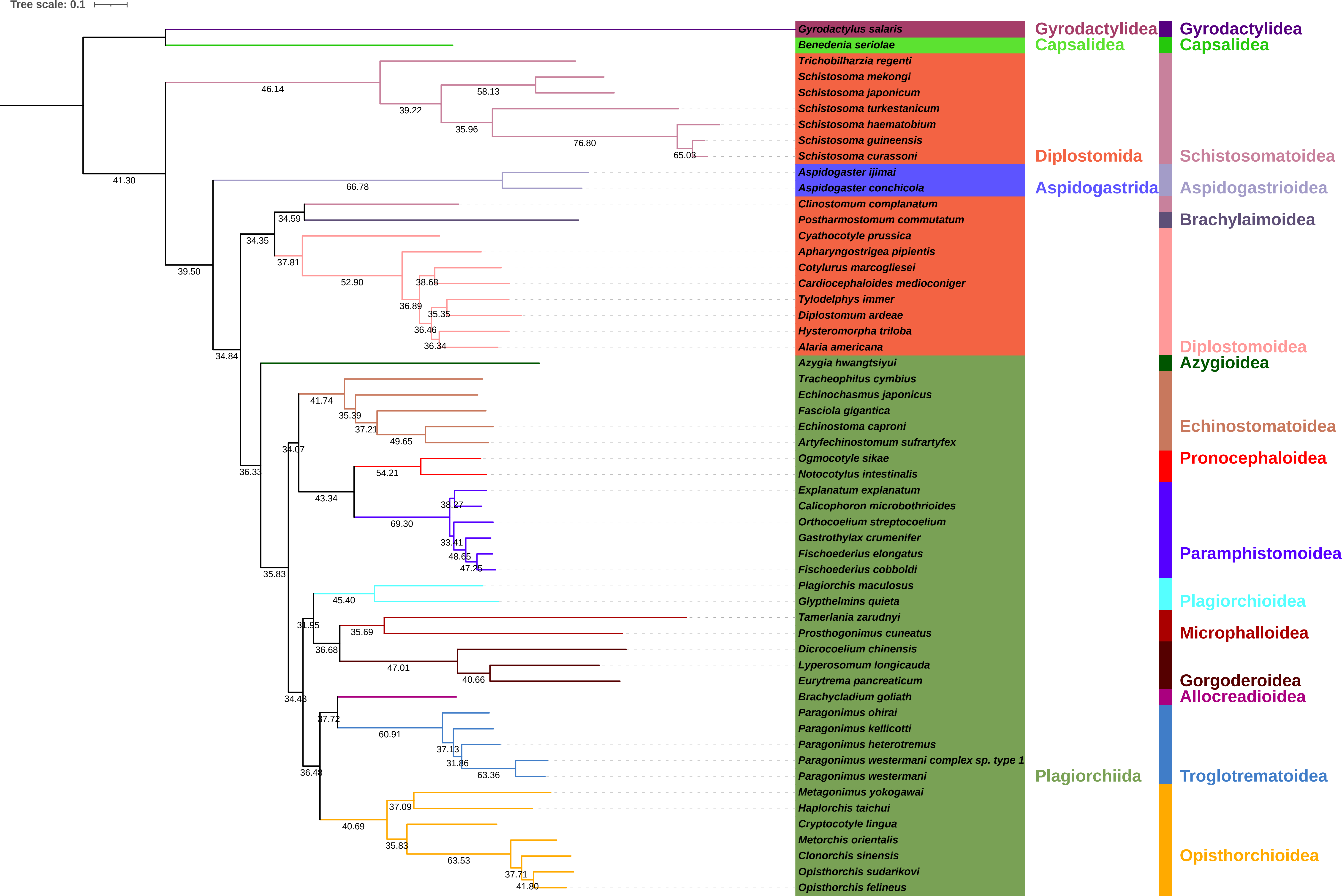

### S37-NUCIQm-gCF.pdf

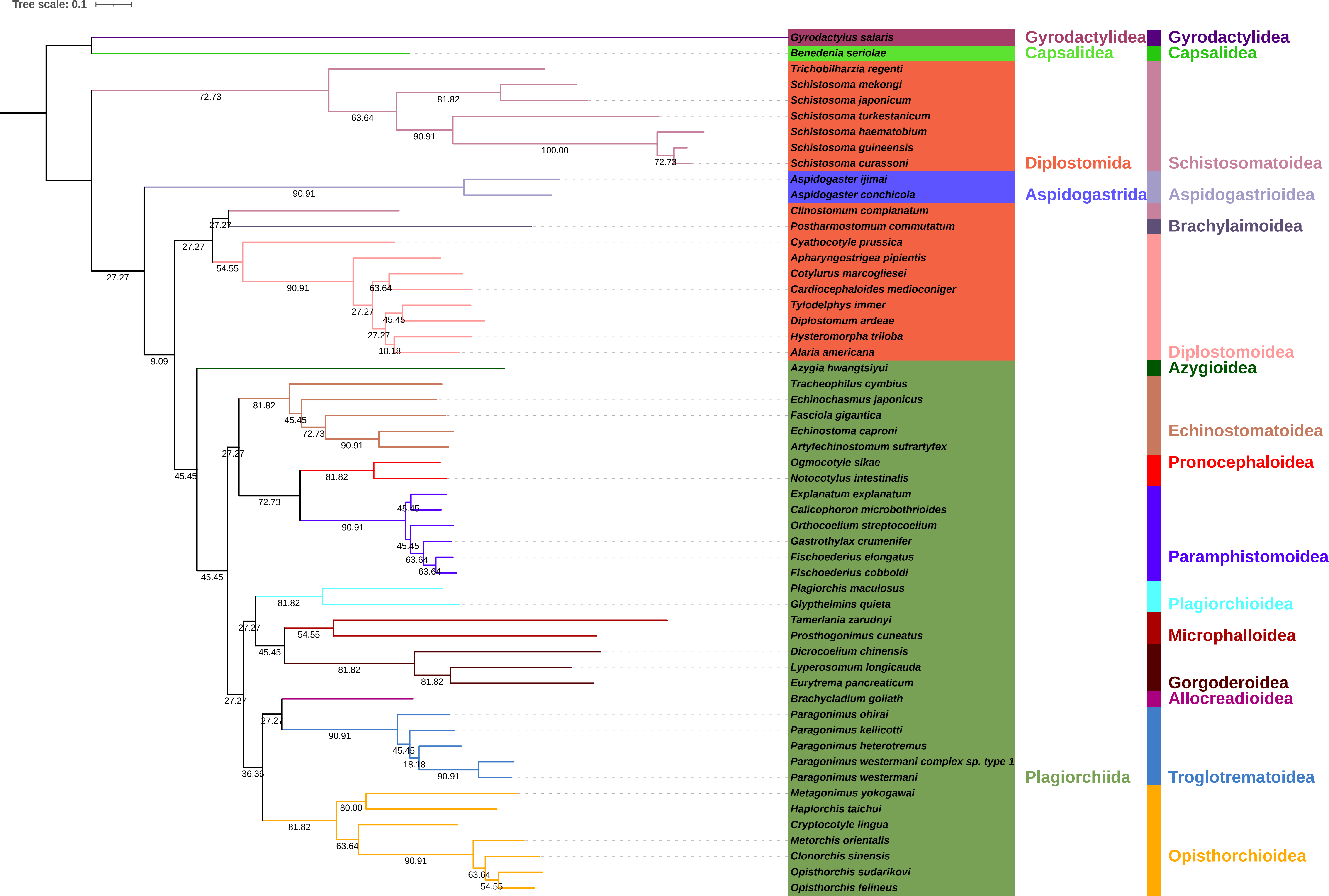

### S38-AAc-cons.pdf

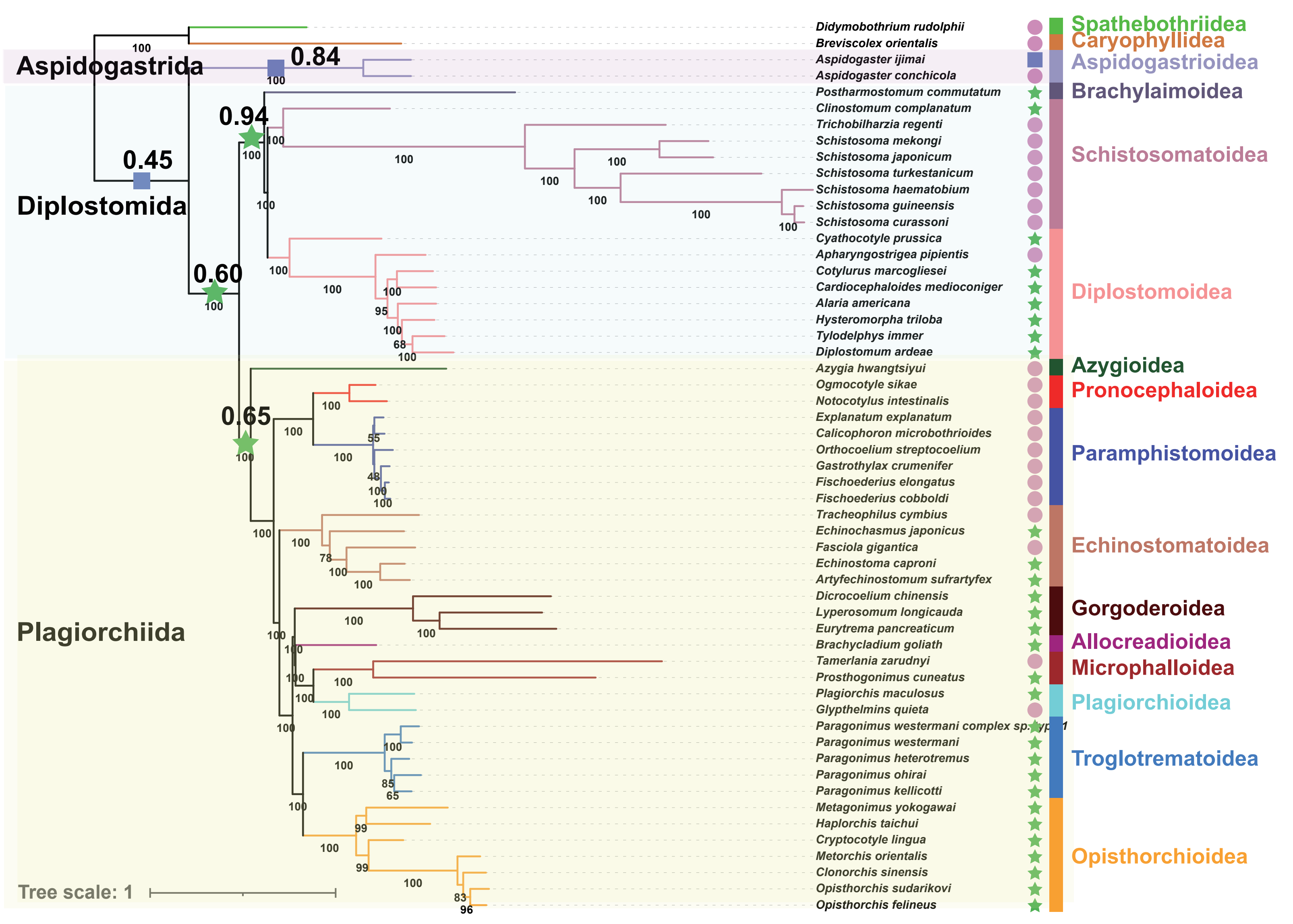

### S39-AAPB-IQn.pdf

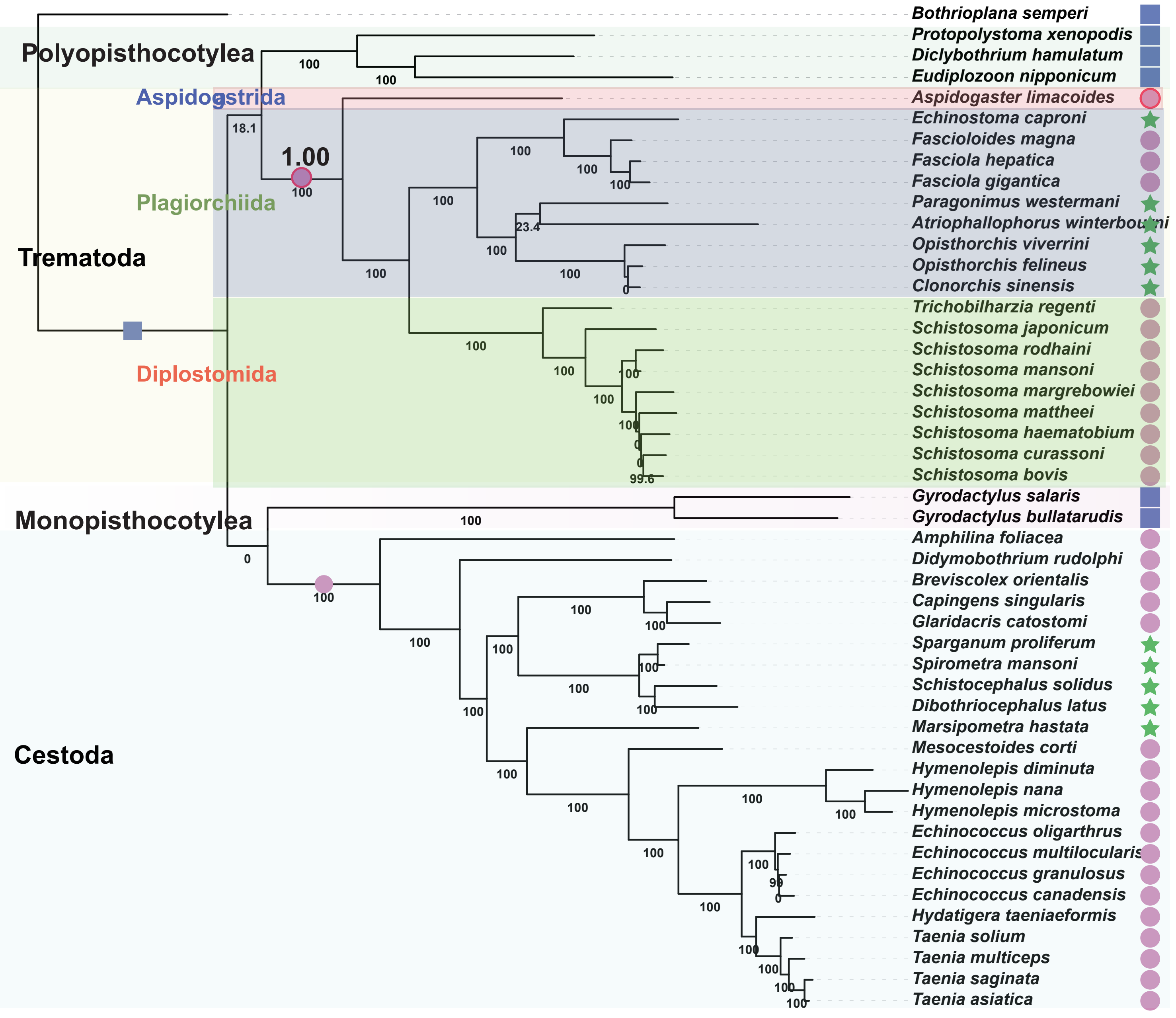

### S40-Mitogenome.pdf

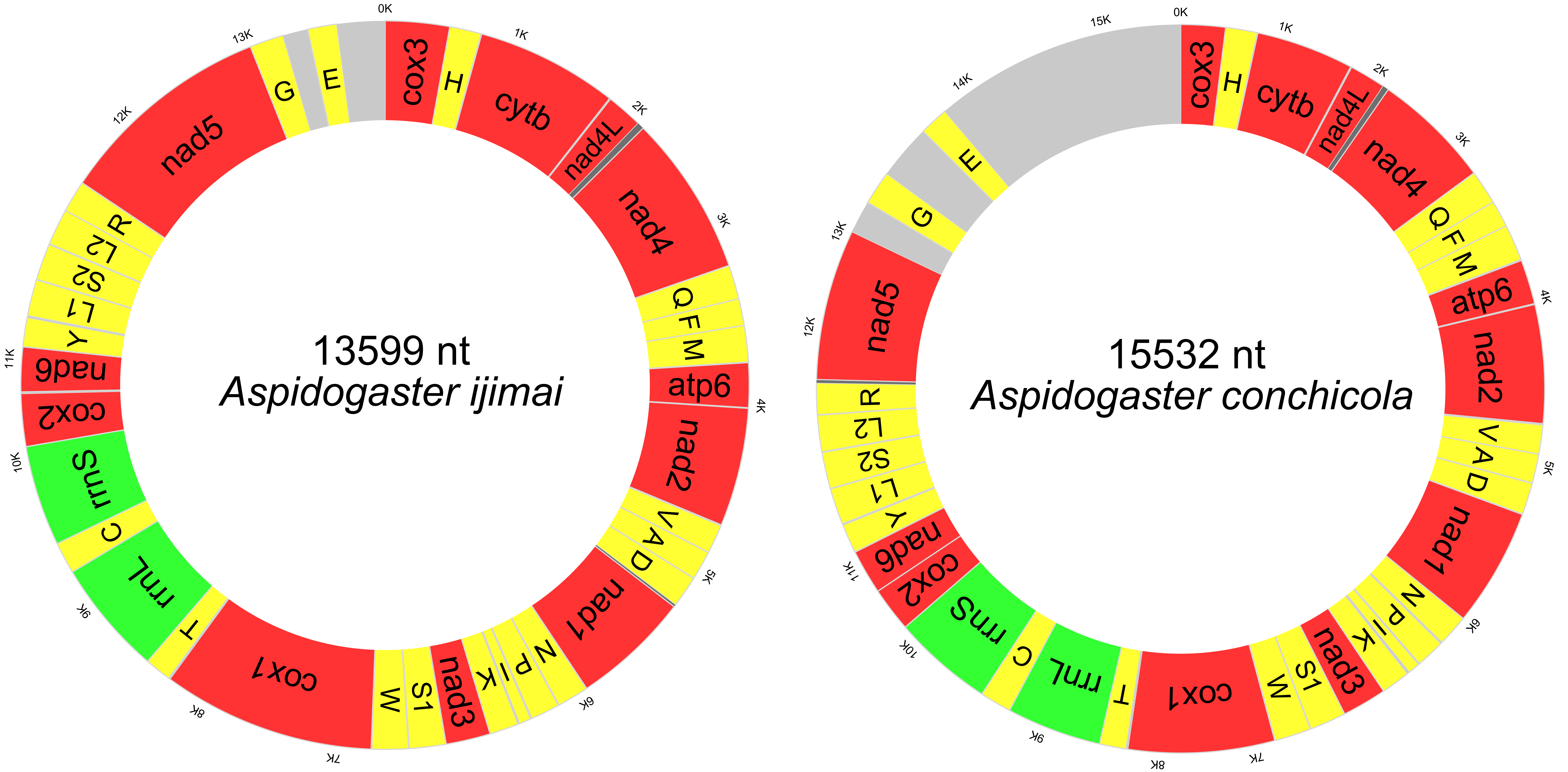

### S41-RSCU.pdf

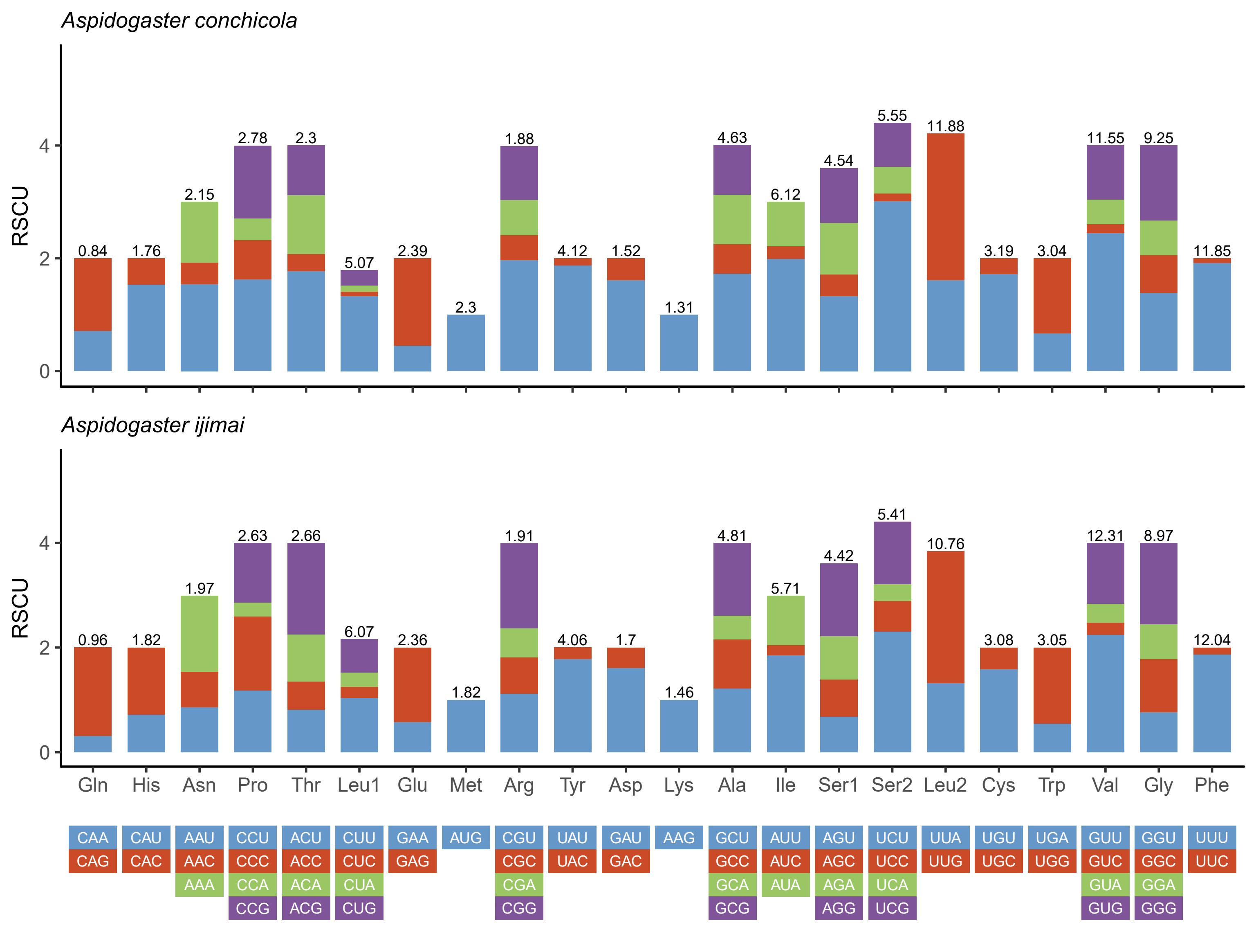

### S42-Stem-loop structures.pdf

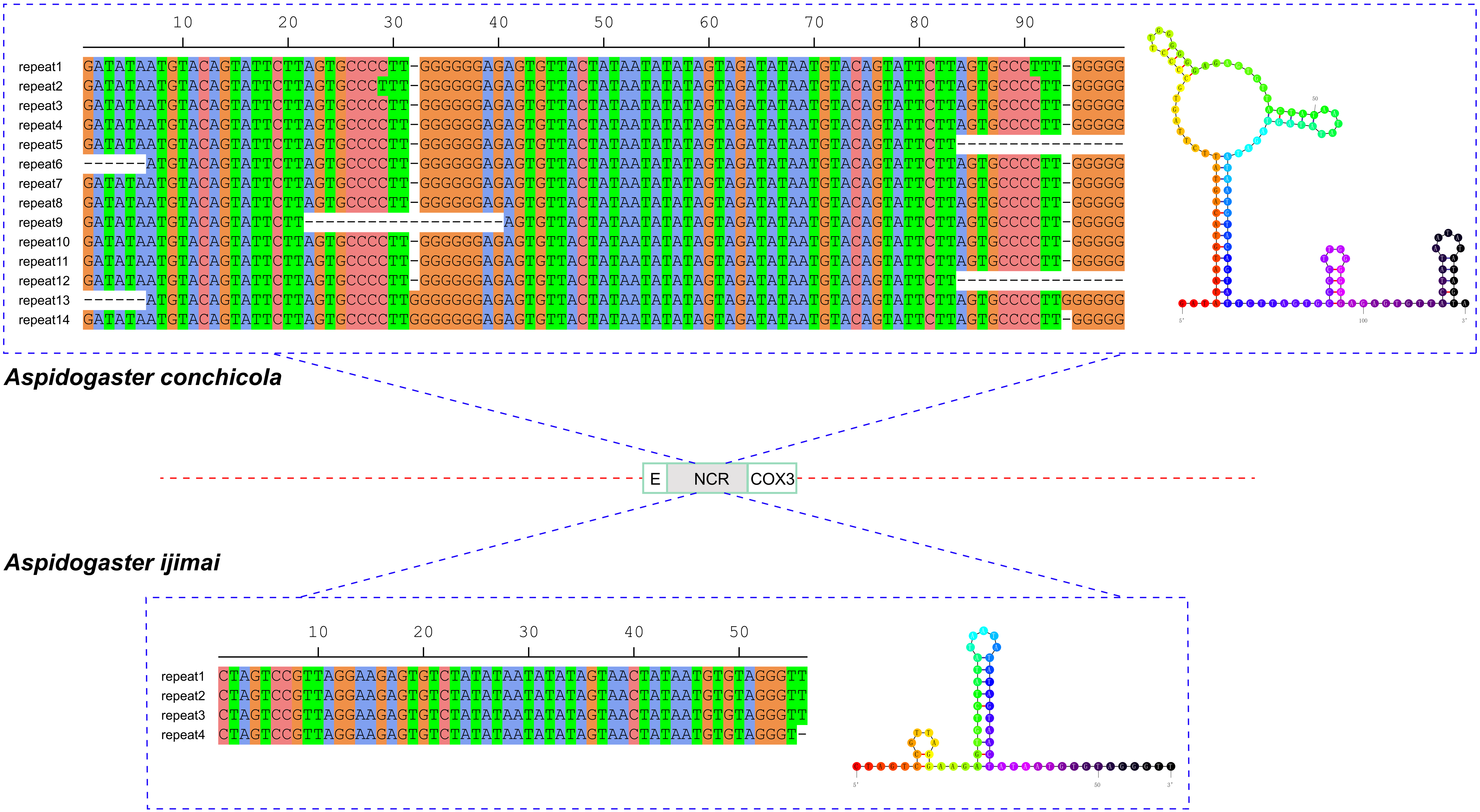

### S43-Nucleotide diversity.pdf

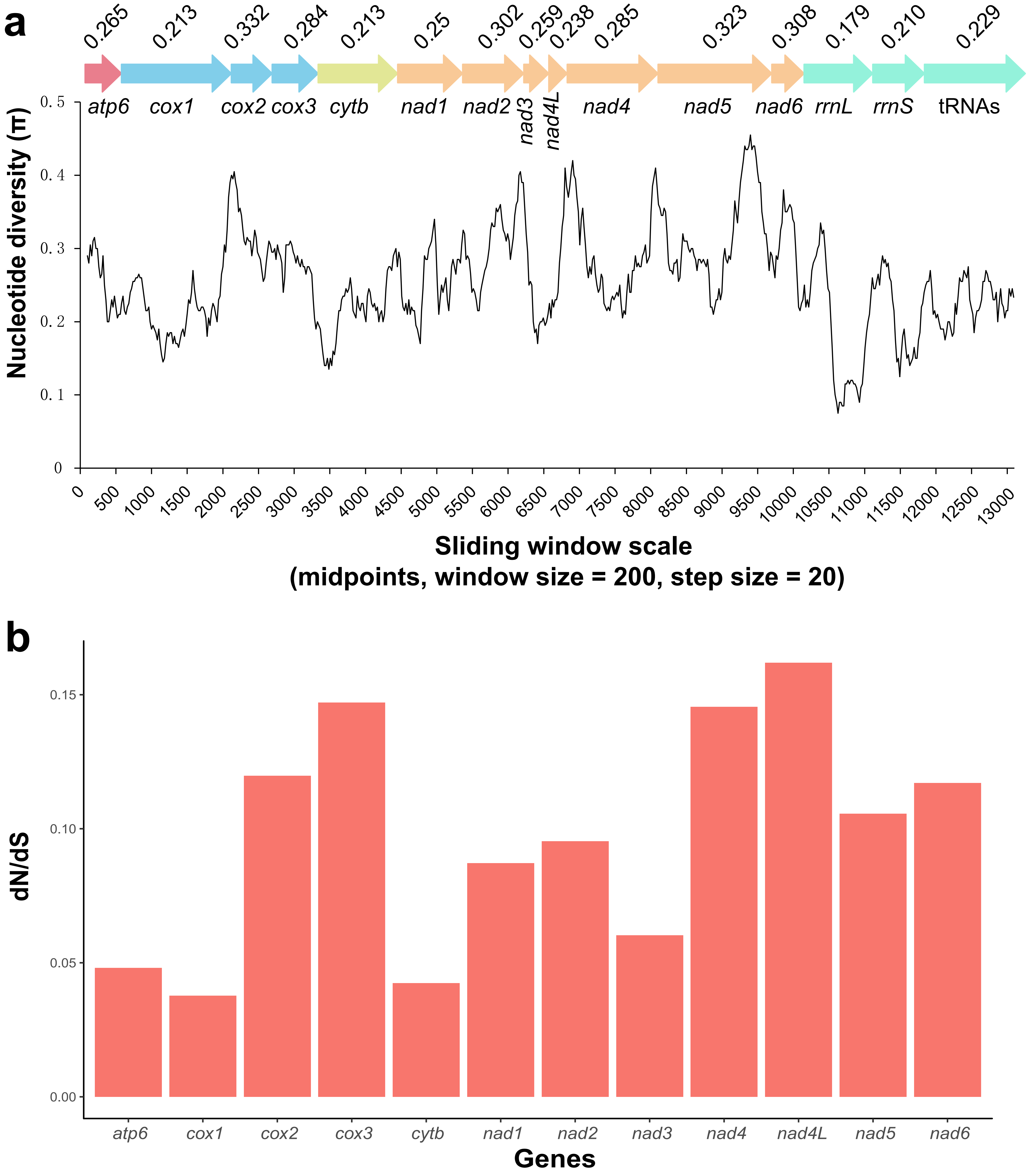

### S44-AAIQPm.pdf

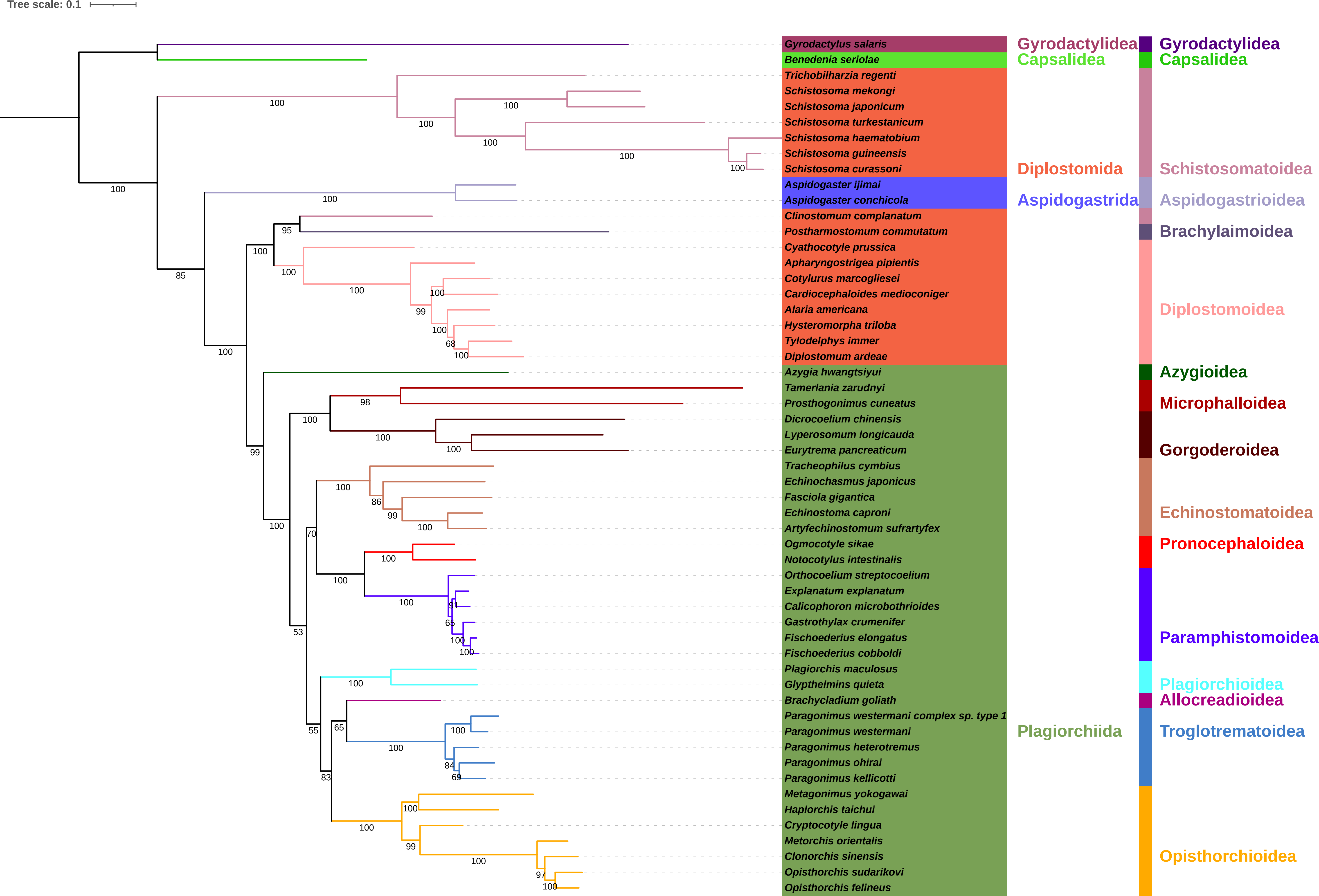

### S45-AAMBPm.pdf

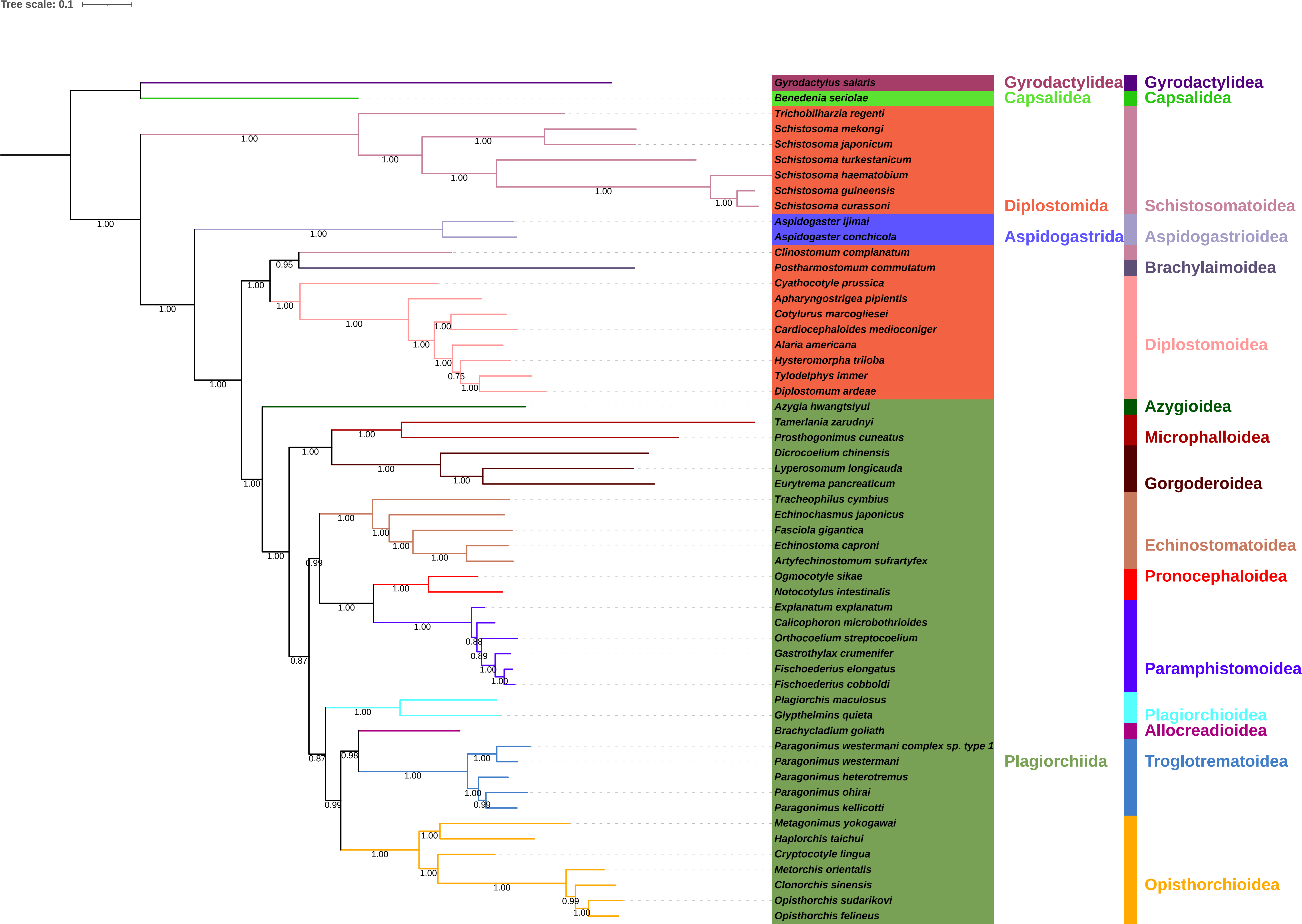

### S46-NUCIQPm.pdf

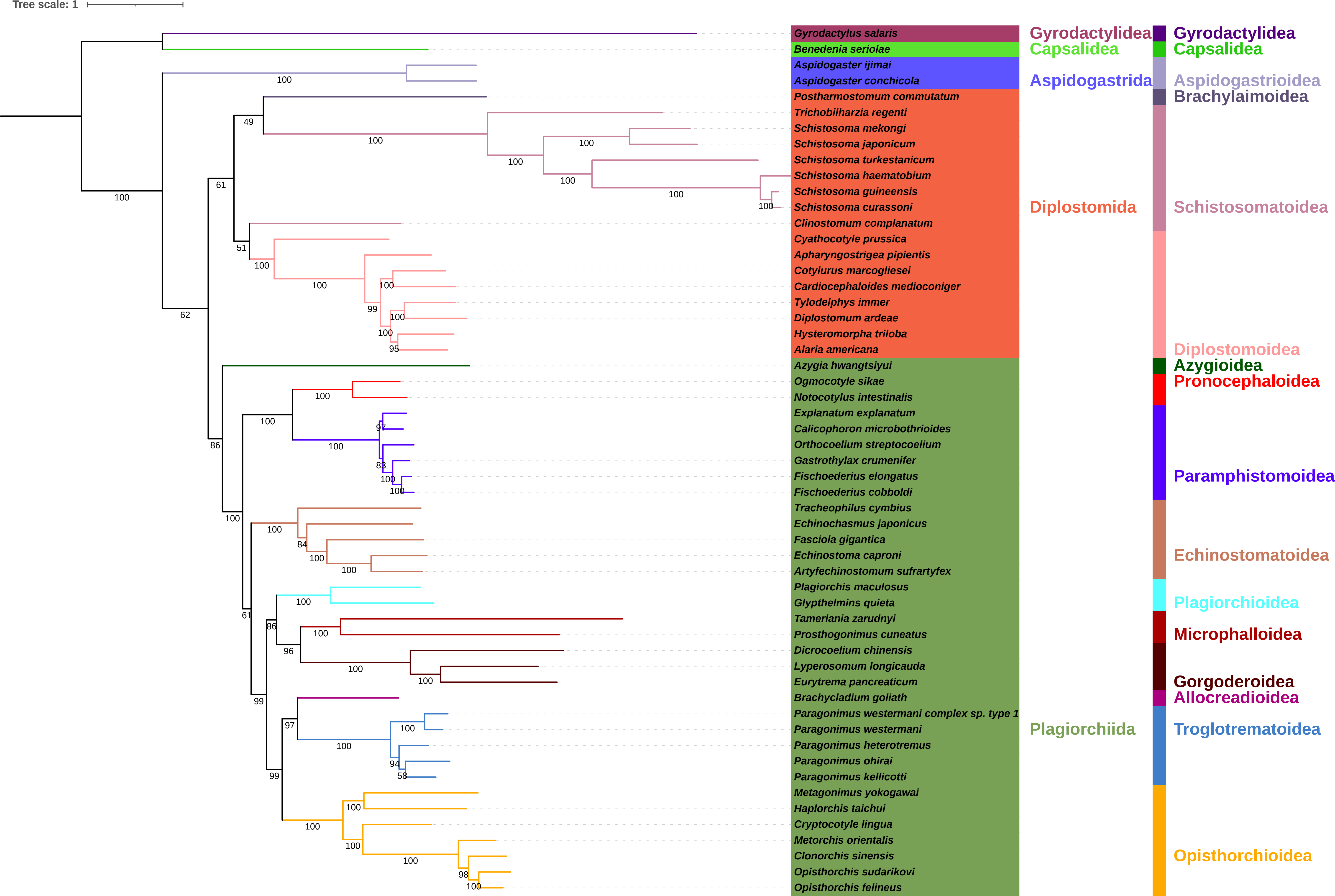

### S47-NUCMBPm.pdf

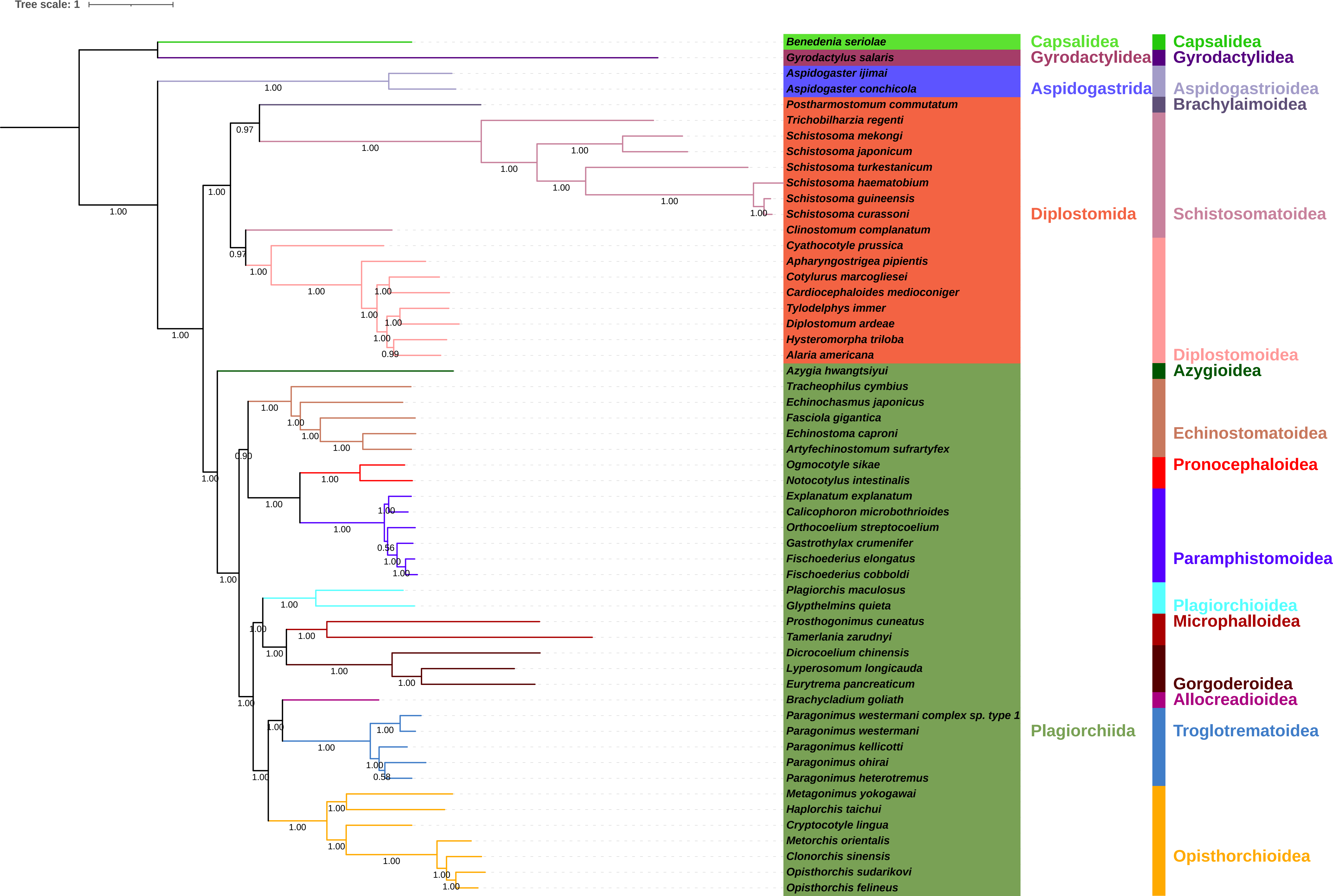

### S48-AAIQPc.pdf

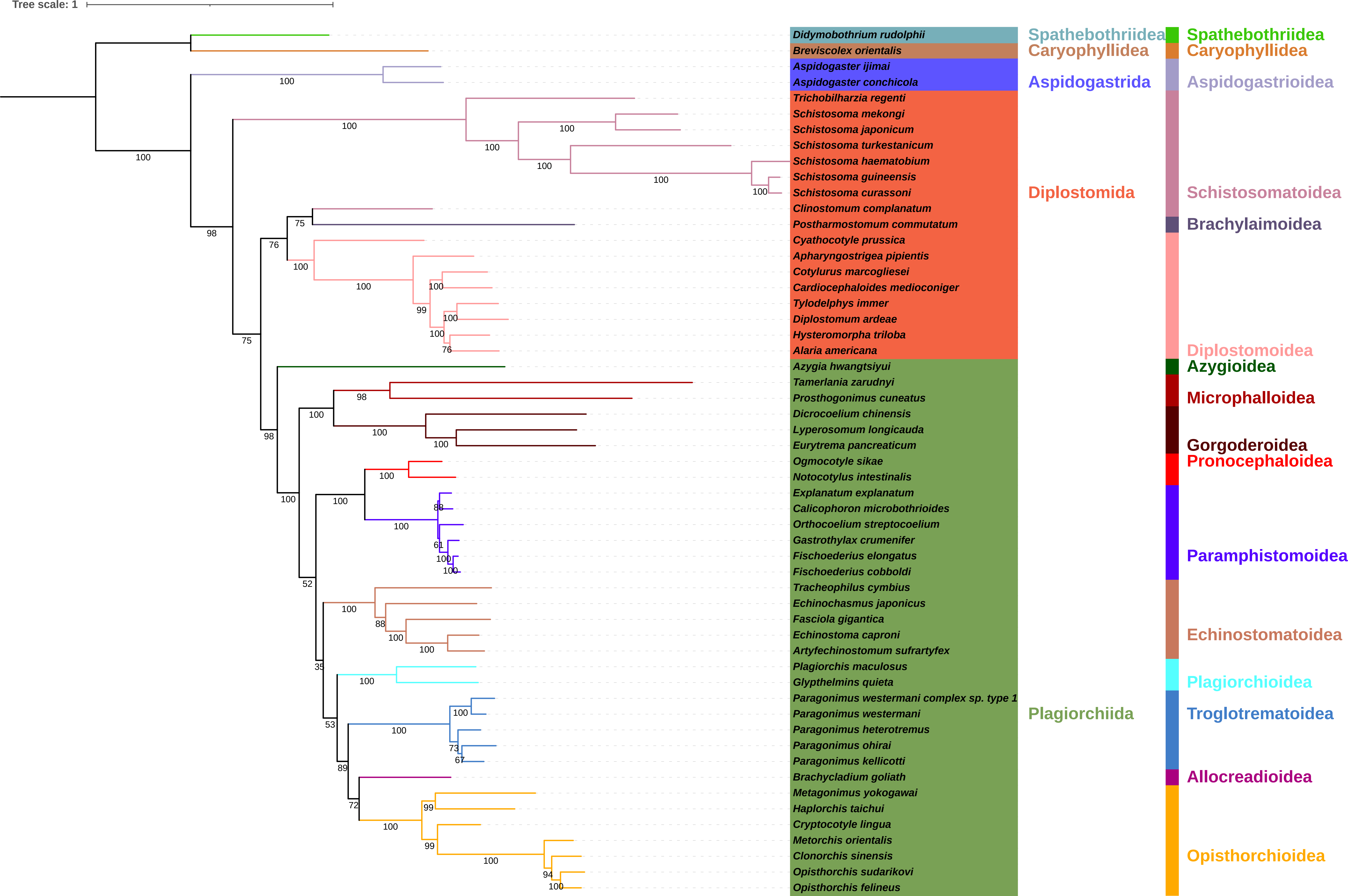

### S49-AAMBPc.pdf

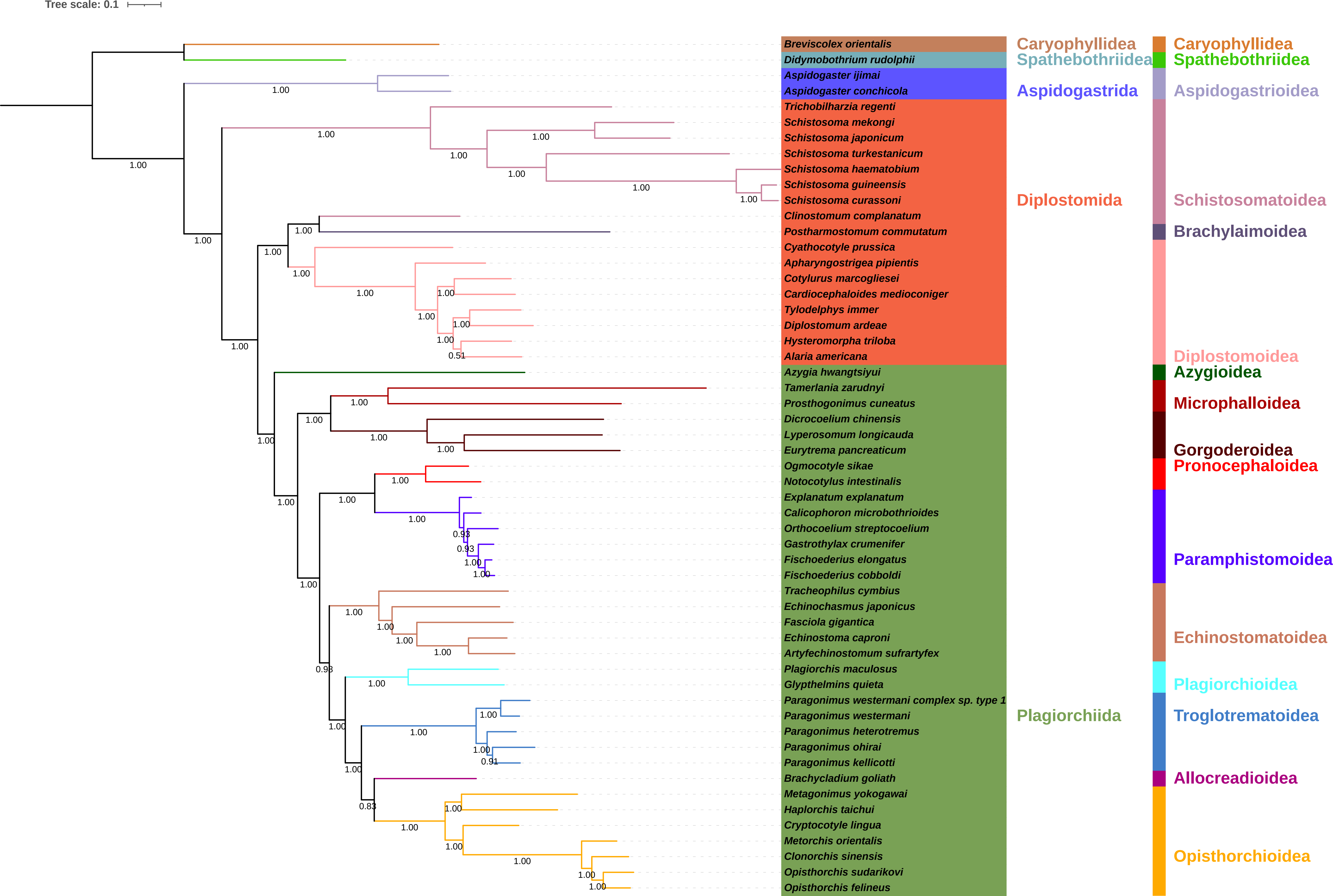

### S50-NUCIQPc.pdf

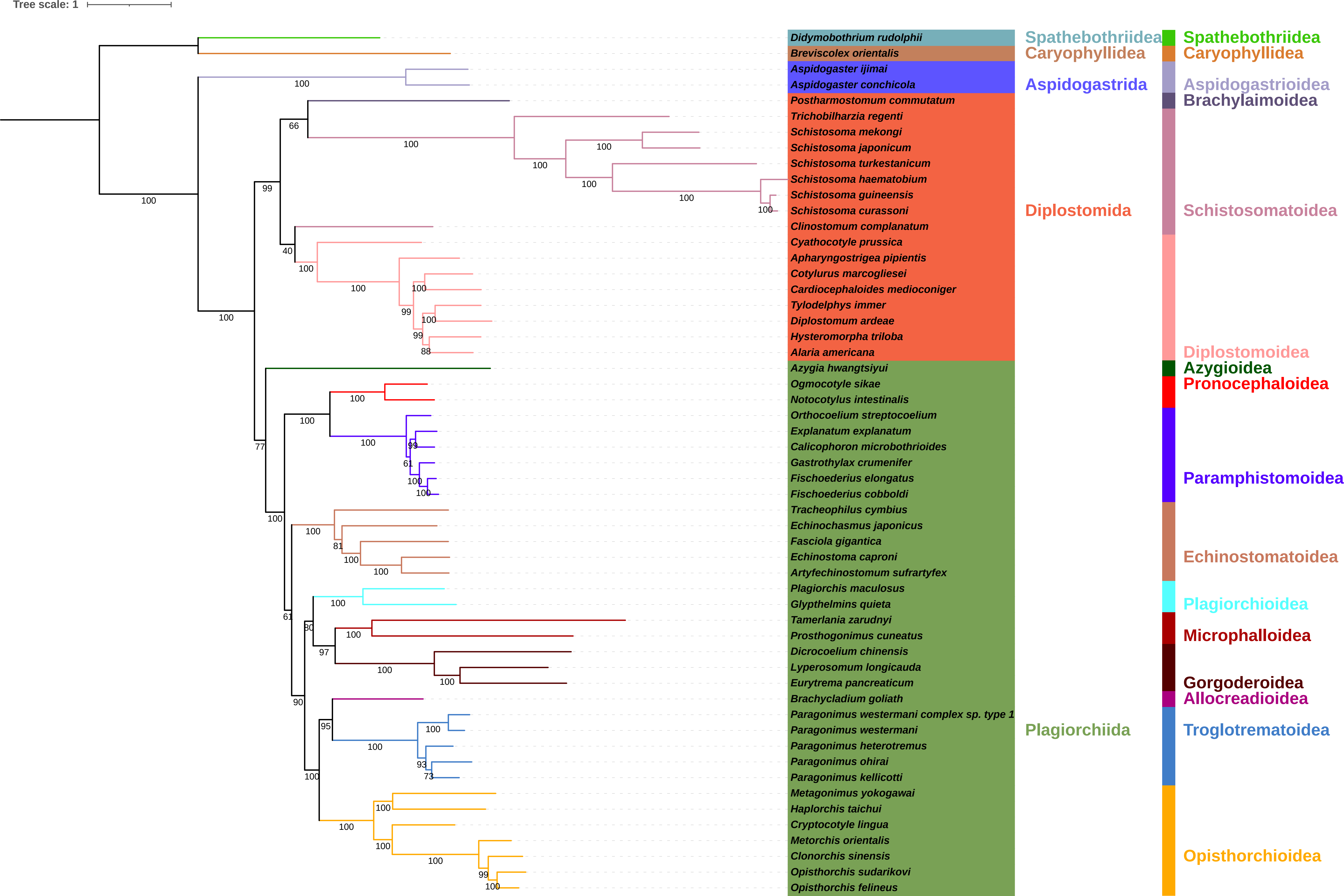

### S51-NUCMBPc.pdf

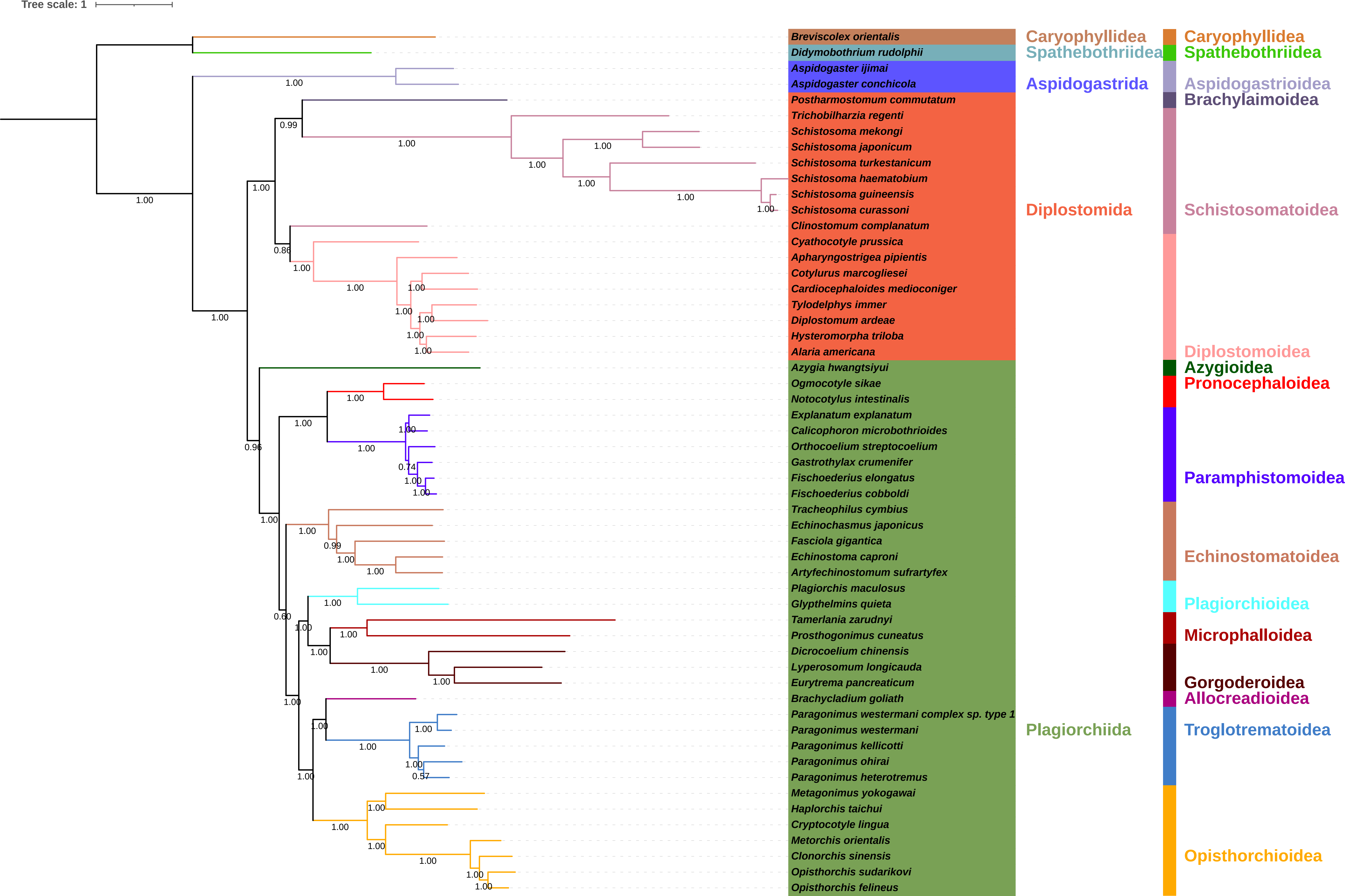

### S52-Comparison of topologies.pdf

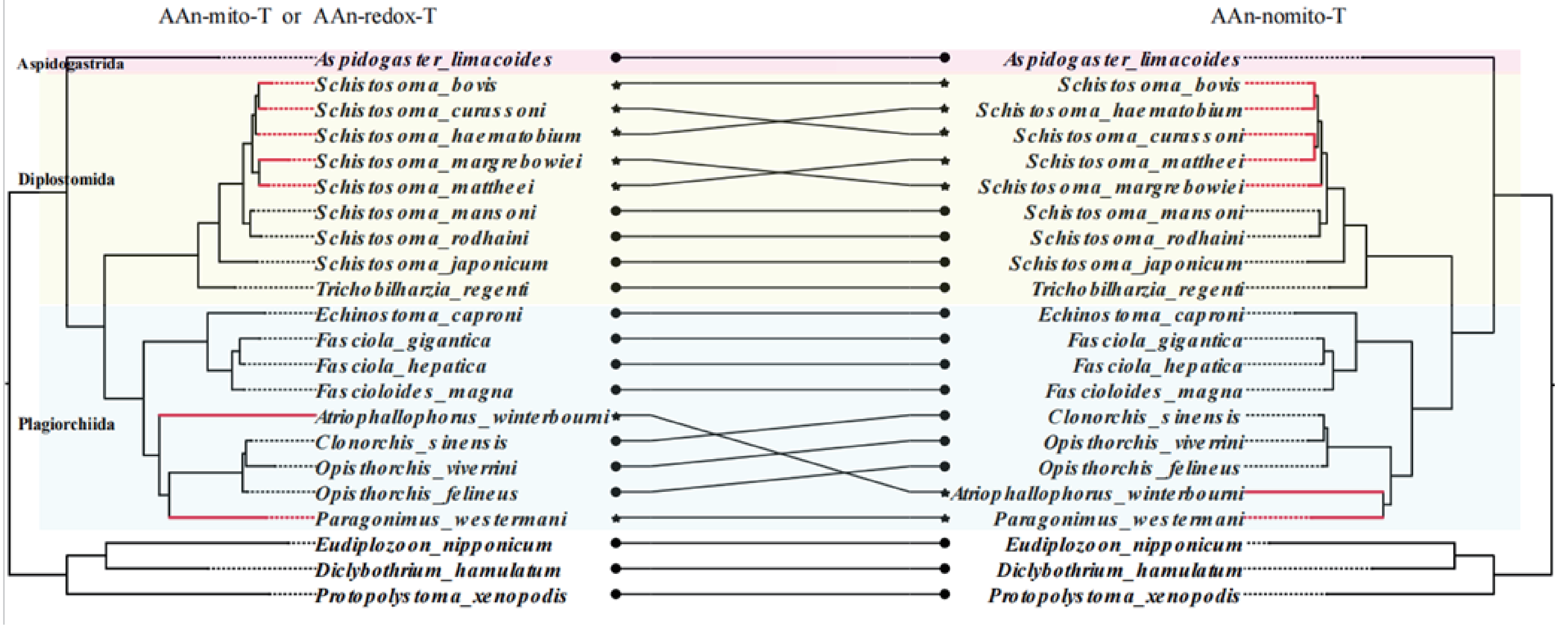
